# Dissecting Response to Cancer Immunotherapy by Applying Bayesian Network Analysis to Flow Cytometry Data

**DOI:** 10.1101/2020.06.14.151100

**Authors:** Andrei S. Rodin, Grigoriy Gogoshin, Seth Hilliard, Lei Wang, Colt Egelston, Russel C. Rockne, Joseph Chao, Peter P. Lee

## Abstract

Cancer immunotherapy, specifically immune checkpoint blockade therapy, has been found to be effective in the treatment of metastatic cancers. However, only a subset of patients achieve clinical responses. Consequently, elucidating immune system-related pre-treatment biomarkers that are predictive with respect to sustained clinical response is a major research priority. Another research priority is evaluating changes in the immune system before and after treatment in responders and non-responders. Our group has been studying immune signaling networks as an accurate reflection of the global immune state. Flow cytometry (FACS, Fluorescence-activated cell sorting) data characterizing immune signaling in peripheral blood mononuclear cells (PBMC) from gastroesophageal adenocarcinoma (GEA) patients were used to analyze changes in immune signaling networks in this setting. Here, we describe a novel computational pipeline developed by us to perform secondary analyses of FACS data using systems biology / machine learning techniques and concepts. It is centered around comparative Bayesian network analyses of immune signaling networks and is capable of detecting true positive signals that conventional methods (such as FlowJo manual gating) might miss. Future studies are planned to validate and follow up immune markers (and combinations / interactions thereof) associated with clinical responses that were identified by this computational pipeline.

## 1. Introduction

Automated analysis of high-dimensional cytometry data is an active bioinformatics research area [1,2]. Increasingly more sophisticated approaches, mostly of the data mining / machine learning variety, have recently been proposed for the primary analysis of such data [3-7]. However, emphasis remains on *primary* data analysis, which is largely reduced to automated gating, clustering, visualization and, subsequently and optionally, cellular identity / population type assignment [8-10]. Identification of clinically useful markers (and, perhaps even more importantly, marker combinations) is a *secondary* data analysis task, belonging to the realm of computational systems biology. To the best of our knowledge, there is currently no systems biology data analysis pipeline aimed specifically at high-dimensional data (including cytometry) in the immuno-oncology domain [11-14]. For example, recent works in the field [15-19] either reduce the markers’ deduction to classification and semi-manual / pairwise combinatorics, or rely on ontological protein-protein interaction networks. Although these analyses are elegant and valid, their completeness and generalizability are uncertain. Notably, such analyses are deficient in automatically inferring higher-order interactions from the data. This is where network-based approaches, such as Bayesian network (BN) modeling, may be useful.

To the best of our knowledge, there has been only one flow cytometry dataset (RAF signaling pathway) subjected to the BN treatment to-date [20-24]. While valuable as a proof-of-concept [20,21], and as a useful platform for further BN methodology development [22-24], it did not lead to wider acceptance of the BN methodology in the field of computational flow cytometry. In this study, we introduce a comprehensive BN-centered analysis strategy aimed at the FACS (fluorescence-activated cell sorting) data analysis in the immuno-oncology context. As a part of this strategy, we introduce a novel technique, based on the “indicator/contrast variable”, for comparing similar but distinct BNs (such as generated in responders vs. non-responders, before vs. after treatment, etc.)

Here we show an application of BN analysis to the gastroesophageal adenocarcinomas (GEA) immuno-oncology data. GAE are comprised of both stomach and esophageal cancers in aggregate, accounting for a large proportion of global cancer-related mortality [25]. In addition to surgery, chemotherapy, radiotherapy, and targeted therapy, immunotherapies such as immune checkpoint blockade (ICB) can serve as promising treatment options for GEA patients. However, the current response rate to ICB in such patients is still very low, with objective response rates less than 20% in all-comers [26]. Therefore, it is critical to identify biomarkers that can effectively predict clinical response to ICB in a subset of patients.

Currently, reliable predictive ICB biomarkers for GEA outside of genomic microsatellite instability (MSI) and Epstein-Barr virus (EBV) detection remain very limited; such potential predictive biomarkers can be identified based on the systemic immunological features that can be measured by using peripheral blood immune cells from GEA patients.

In our study, we have collected peripheral blood mononuclear cell (PBMC) samples from metastatic GEA patients receiving anti-PD-1 (Pembrolizumab) along with radiation therapy. Three immune signaling flow cytometry panels (Adaptive, Checkpoint, Immune) were generated from each sample (see *Materials and Methods: Flow Cytometry* for detailed panel descriptions). Importantly, we had PBMC samples from patients at baseline before the therapy (day 1) and after therapy (day 21). As a result, immunological changes in PBMC samples could be correlated to clinical response to immunotherapy in these patients.

The results of this study are summarized in the three sections below. The first section details application of BN analysis to the three immune signaling flow cytometry panel datasets in all patients at day 1. We further scrutinize the resulting BNs to identify biomarkers (and combinations thereof) predictive for immunotherapy response. In the second section, we perform the more granular analyses, separating the data into four different immune cell types and contrasting resulting networks between responders and non-responders, at day 1 and day 21. In the third section, we discuss the BN validation using statistical criteria, and compare BN modeling with other conventional multivariate analysis methods.

A somewhat unexpected result, described in the third section, was the clear dichotomy between the two groups of analysis methods --- (i) more conventional (in the immuno-oncology context) FlowJo, regression and correlation approaches *vs*. (ii) BNs, multivariate classifiers and distribution distance metrics. In one striking example, the latter methods uniformly pinpointed a strong predictive biomarker that the former methods failed to detect. Therefore, one of the conclusions of this study is that the investigators should probably augment the conventional flow cytometry data analysis methodology with more sophisticated analysis tools.

## 2. Results

### 2.1. Basic computational pipeline, and baseline immune signaling FACS panels analyses

Our basic computational pipeline consists of (1) discretizing FACS measurements (compensated fluorescence intensity values, obtained directly from FlowJo FACS software [27] *via* exported FCS files) into 2-8 equal-frequency bins, (2) performing full BN analysis (i.e., reconstructing the complete BN) for one FACS panel, clinical response status variable, and other available variables using our BN modeling software (see *Materials and Methods: Bayesian networks modeling* for details), (3) isolating a Markov blanket/neighborhood (MN) of the response status variable, (4) analyzing the interplay of (usually no more than 4-8) immune system-related markers within the MN, (5) generating a compact set of rules for predicting response status from the markers’ values, and (6) associating these rules with immune cell population subtypes. Afterwards, results obtained at different immunotherapy timepoints can be compared and contrasted (which is described in the next section, “contrast” analyses).

Three types of immune signaling FACS panels were analyzed, Checkpoint, Innate and Adaptive, with 14 immune markers in each (see *Materials and Methods: Flow cytometry* for details). We will first illustrate the pipeline on the example of Checkpoint immune signaling FACS panel in GEA patients. The data contained 14 relevant PBMC immune marker variables measured (at day 1, before therapy, and day 21, after therapy) in 13 patients undergoing immunotherapy. Four patients exhibited significant (< −25% radius) tumor shrinkage and were labeled responders, with remaining nine labeled non-responders. All 13 patients were pooled together into a single dataset, and the complete dataset was used for three-bin (“low”, “medium” and “high”) equal-frequency discretization of 14 marker variables.

Figure 1 depicts the full reconstructed Checkpoint immune signaling BN at day 1 (pre-treatment). The response status variable is designated “Tsize”, for “absolute tumor size (radius) change”, and is shown as the red node in the graph. It is clear from the network that response status is directly strongly linked with five markers (TIGIT, CD4, CD8, CD160, 4-1BB, designated by dark grey nodes in the network) and, less strongly, with six other markers (TIM3, CD45RA, OX40, CXCR5, KLRG1, LAG3, light grey nodes in the network). Because the edges connecting Tsize to the latter six are, on average, significantly weaker than the edges between the latter six and the former five, we will only consider the former five markers as potential predictors of response to therapy. For example, although there is a direct edge between Tsize and CXCR5, it is very weak compared to the Tsize --- CD4 and CD4 --- CXCR5 edges, and so we conclude that the Tsize --- CXCR5 edge is likely an artifact of the very strong CD4 --- CXCR5 dependence.

**Figure 1.**
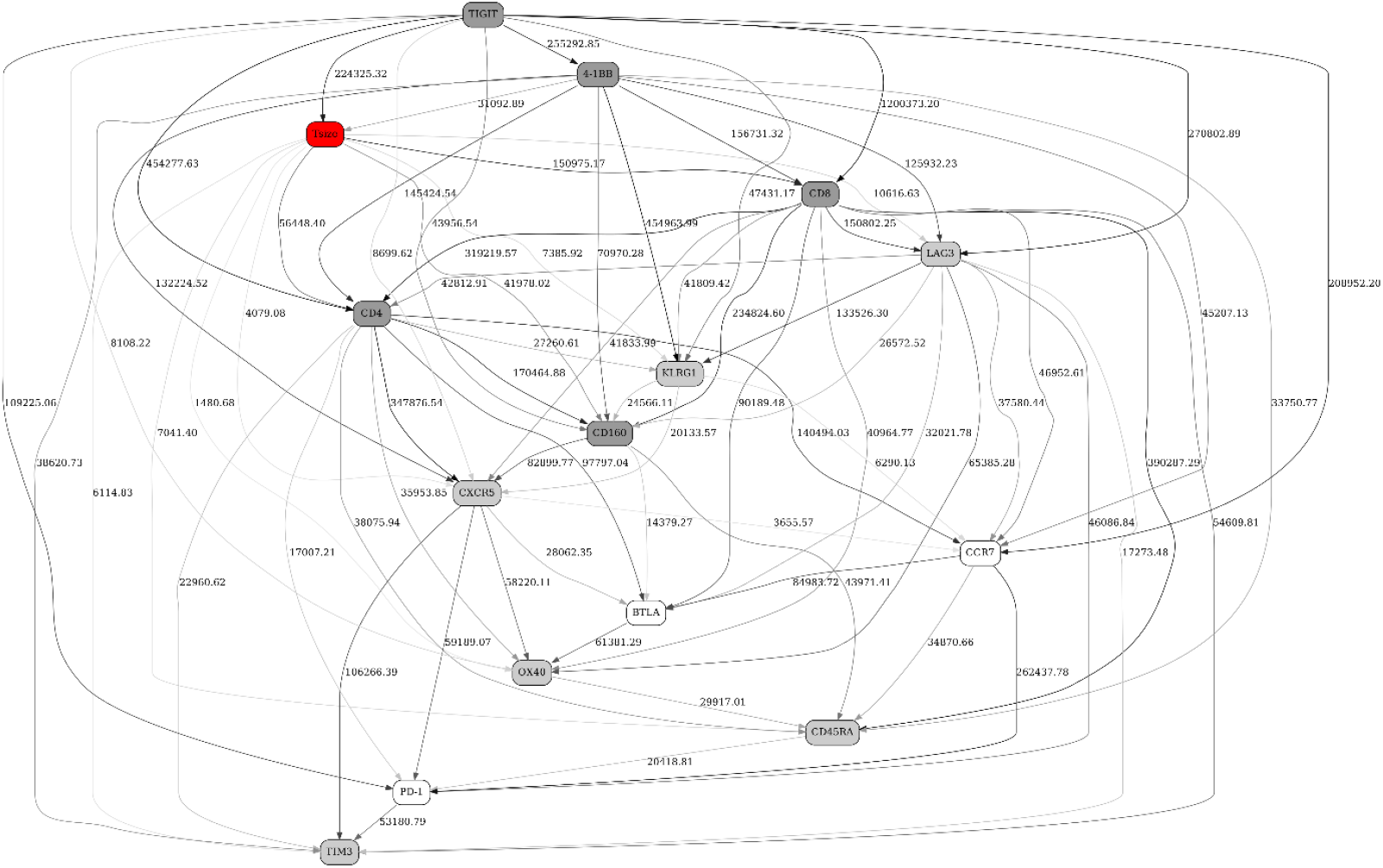
A full BN constructed from FACS data (Checkpoint immune signaling network panel) from 13 patients pre-immunotherapy treatment (day 1). Number next to the edge and edge thickness indicate dependency strength; arrows / edge directionalities distinguish parent/offspring relationships during the BN reconstruction, and do not necessarily imply causation flow (see *Materials and Methods: Bayesian networks modeling* for details). “Tsize” node is the response status variable (4 responders, 9 non-responders). Remaining nodes are immune marker variables. See text for further details.

Recall that the five variables directly influencing response status (TIGIT, CD4, CD8, CD160, 4-1BB) were discretized into three bins, “low”, “medium”, and “high”. Therefore, there are 3^5 = 243 possible configurations for these five variables. This is much better than the 3^14 ∼= 4.8 million configurations in the full dataset before BN-derived dimensionality reduction, but still not feasible for the configuration-by-configuration manual scrutiny. However, we can rank these 243 configurations in order of decreasing frequency in the dataset (which indirectly map to the immune cell population types, from most to least frequent), and select a small number of most frequent configurations (using prespecified, or frequency distribution shape-driven, cutoffs). Then, we can re-sort them in order of decreasing probability of clinical response (tumor shrinkage) given the configuration. We illustrate this approach in Table 1 by showing 15 most frequent configurations, in order of decreasing probability of response --- meaning that responsive patients are associated with higher than randomly expected frequency of top-ranking configurations; these configurations might roughly correspond to particular immune cell population types. (The actual number of configurations, 15, was suggested by the shape of the configuration frequency function, with a noticeable dropoff (∼0.02 to ∼0.017) between the 15^th^ and 16^th^ configurations; the next noticeable dropoff is between the 35^th^ and 36^th^ configurations, and we have chosen the 15 configurations cutoff for presentation compactness).

**Table 1.**
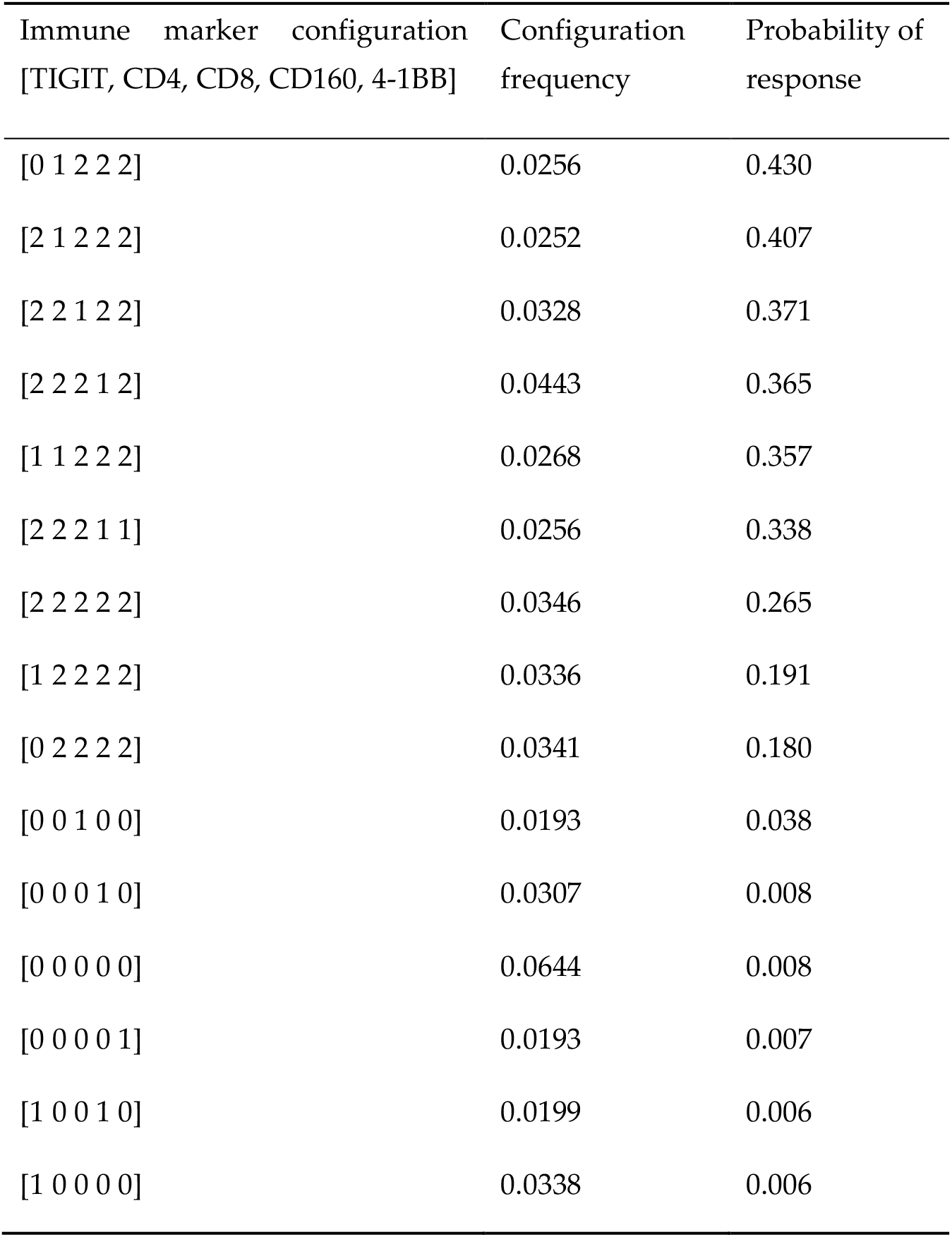
The 15 most frequent predictive immune marker configurations (Checkpoint FACS panel), in descending order of probability of response. First column shows predictive immune marker configurations obtained from the MN of the response status variable, five markers in total (TIGIT, CD4, CD8, CD160, 4-1BB). The values “0”, “1”, “2” in the first column indicate “low”, “medium” and “high” values after discretization. Second column shows configuration frequency; third column – probability of response.

As summarized in Table 1, the analysis results are now compact enough to allow “manual” scrutiny. Note that the above process is completely automated --- the only two adjustable parameters are (i) the number of the potentially predictive variables to be selected from the MN of the response status variable, and (ii) the number of the most frequent marker configurations. In this example, the data itself suggested 5 and 15, respectively.

The above strategy (sorting first by frequency, then by response probability) favors the signals associated with more frequent marker combinations. Alternatively, one can sort by response probability only. Marker combinations associated with very high (or very low) response probabilities are likely to be less frequent. Such strategy is very similar to rare cells subsets discovery algorithms. In general, it is up to the investigator how the conditional probability tables are to be re-sorted once the list of potentially predictive variables is reduced to these inside the MN of response variable. The advantages of the “frequency-then-probability” sorting, as exemplified above, are two-fold: first, higher statistical robustness of discovered signals; second, a (typically) compact set of “rules” describing these signals.

By considering simultaneously the MN of the response variable (Figure 1), and the list of the most frequent marker configurations (Table 1), it is possible to elucidate a set of broad predictive patterns with respect to the clinical response variable. One such pattern, clearly discernible in the marker configuration list (Table 1), is a demarcation line between the top nine and the bottom six configurations. The former correspond to the higher response probability and show, on average, high marker concentrations (more “2”s across the board). The latter correspond to the low response probability and show lower marker concentrations (more “0”s across the board). In addition, Figure 1 suggests TIGIT (and, to a lesser extent, CD8) as the strongest individual predictor(s) of the clinical response. Indeed, when discretized into two bins, high TIGIT values correspond to 0.28572 response probability, while low TIGIT values --- 0.051155 response probability.

We will now consider Innate and Adaptive panels. First, Figure 2 depicts the BN generated from the same 13 patients using Innate immune signaling FACS panel. Out of 14 relevant immune markers, response status (red “Tsize” node in the graph) is directly strongly linked with eight markers (HLA-DR, PD-L1, CD3, CD20, CD83, CD1c, CD14, CD33, dark grey nodes in the network) and, less strongly, with one other marker (CD141, light grey node in the network).

**Figure 2.**
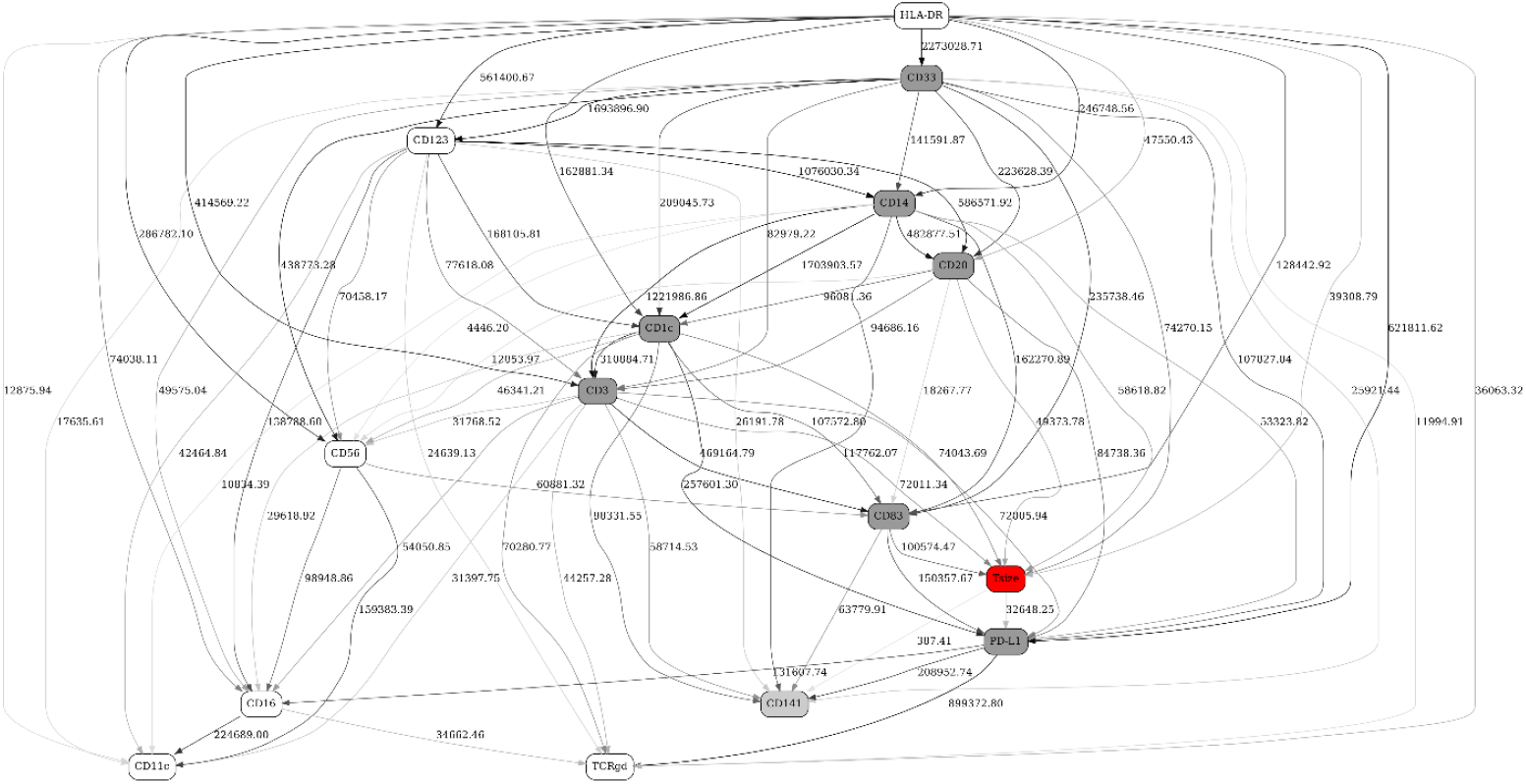
A full BN constructed from FACS data (Innate immune signaling network panel) from 13 patients pre-immunotherapy treatment (day 1). See Figure 1 legend for further details.

An 8-marker configuration frequency table is shown in Table 2. The actual number of configurations shown (20) was suggested by the shape of the configuration frequency function, which had the first noticeable dropoff at 20.

**Table 2.** The 20 most frequent predictive immune marker configurations (Innate FACS panel), in descending order of probability of response. First column shows predictive immune marker configurations obtained from the MN of the response status variable, eight markers in total (HLA-DR, PD-L1, CD3, CD20, CD83, CD1c, CD14, CD33). The values “0”, “1”, “2” in the first column indicate “low”, “medium” and “high” values after discretization. Second column shows configuration frequency; third column – probability of response.

| Immune marker configuration<br>[HLA-DR, PD-L1, CD3, CD20,<br>CD83, CD1c, CD14, CD33] | Configuration<br>frequency | Probability of<br>response |
| --- | --- | --- |
| [2 1 0 1 2 1 2 2] | 0.0182 | 0.648 |
| [2 2 2 1 2 2 1 0] | 0.0096 | 0.639 |
| [2 1 1 1 2 1 2 2] | 0.0308 | 0.604 |
| [2 2 2 1 2 2 1 1] | 0.0098 | 0.586 |
| [2 1 1 2 2 1 2 2] | 0.0163 | 0.369 |
| [2 2 2 2 2 2 2 2] | 0.0269 | 0.357 |
| [1 2 2 2 1 2 1 0] | 0.0081 | 0.249 |
| [1 2 2 2 1 2 1 1] | 0.0176 | 0.246 |
| [2 1 2 1 2 1 0 2] | 0.0086 | 0.202 |
| [2 2 2 2 2 1 2 2] | 0.0430 | 0.141 |
| [2 2 2 2 2 1 1 2] | 0.0090 | 0.138 |
| [1 2 2 2 1 2 0 0] | 0.0080 | 0.138 |
| [1 2 2 2 1 2 0 1] | 0.0078 | 0.133 |
| [0 0 0 0 0 0 1 0] | 0.0147 | 0.063 |
| [0 0 0 0 0 0 1 1] | 0.0074 | 0.037 |
| [0 0 0 0 0 0 0 1] | 0.0086 | 0.026 |
| [0 0 0 0 1 0 0 0] | 0.0158 | 0.015 |
| [0 0 0 0 0 0 0 0] | 0.0603 | 0.014 |
| [0 0 0 0 0 0 2 1] | 0.0085 | 0.013 |
| [1 0 0 0 0 0 0 0] | 0.0187 | 0.007 |

Similar to Table 1 (Checkpoint FACS panel), the most striking feature of the marker configuration list (Table 2) is a clear demarcation line between the top 13 and the bottom 7 configurations. The former correspond to the higher response probability and show, on average, high marker concentrations (more “2”s across the board). The latter correspond to the low response probability and show lower marker concentrations (more “0”s across the board). However, in contrast to the Checkpoint immune signaling network results, no particular markers (out of 8) stand out as the strongest individual predictors. In conclusion, on average, high values of HLA-DR, PD-L1, CD3, CD20, CD83, CD1c, CD14 and CD33 predict favorable response to immunotherapy.

Finally, Figure 3 depicts the BN generated from the same 13 patients using Adaptive immune signaling FACS panel. Out of 14 immune markers, response status (red “Tsize” node in the graph) is directly strongly linked with four markers (CXCR3, CCR4, CD8, CXCR5, dark grey nodes in the network) and, less strongly, with five other markers (CCR10, ICOS, CD73, CD4, CD45RA, light grey nodes in the network).

**Figure 3.**
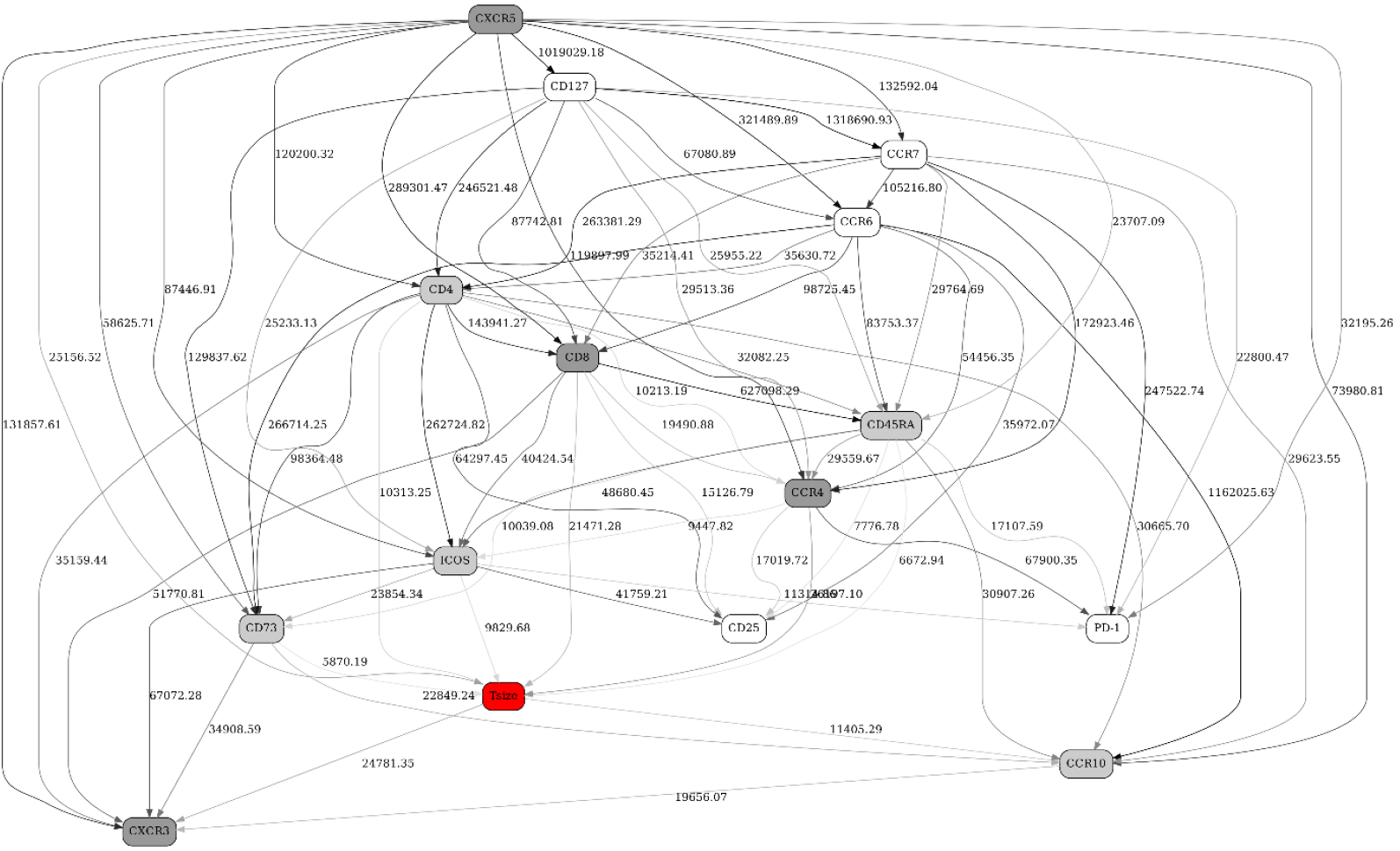
A full BN constructed from FACS data (Adaptive immune signaling network panel) from 13 patients pre-immunotherapy treatment (day 1). See Figure 1 legend for further details.

A 4-marker configuration frequency table is shown in Table 3. The actual number of configurations shown (10) was suggested by the shape of the configuration frequency function, which had the first noticeable dropoff at 10. Similar to the Tables 1 and 2, the top rows (higher probability of response) show, on average, high marker concentrations (more “2”s across the board), and the bottom rows (low probability of response) show lower marker concentrations (more “0”s across the board). Notably, the 5^th^ row presents an almost perfect “borderline” case --- medium concentrations for all markers (all “1”s), and an intermediate probability of response (0.08891941). No particular markers (out of 4) stand out as the strongest individual predictors. In conclusion, on average, high values of CXCR3, CCR4, CD8 and CXCR5 predict favorable response to immunotherapy.

**Table 3.** The 10 most frequent predictive immune marker configurations (Adaptive FACS panel), in descending order of probability of response. First column shows predictive immune marker configurations obtained from the MN of the response status variable, four markers in total (CXCR3, CCR4, CD8, CXCR5). The values “0”, “1”, “2” in the first column indicate “low”, “medium” and “high” values after discretization. Second column shows configuration frequency; third column – probability of response.

| Immune marker configuration<br>[CXCR3, CCR4, CD8, CXCR5] | Configuration<br>frequency | Probability of<br>response |
| --- | --- | --- |
| [2 1 2 2] | 0.0352 | 0.406 |
| [2 2 2 2] | 0.1520 | 0.377 |
| [2 2 1 2] | 0.0401 | 0.347 |
| [1 2 2 2] | 0.0384 | 0.226 |
| [1 1 1 1] | 0.0404 | 0.089 |
| [0 0 1 0] | 0.0351 | 0.039 |
| [0 1 0 0] | 0.0377 | 0.018 |
| [0 0 0 1] | 0.0412 | 0.008 |
| [0 0 0 0] | 0.1240 | 0.008 |
| [1 0 0 0] | 0.0289 | 0.005 |

### 2.2. Separation into major immune cell types, and “contrast” (before/after immunotherapy, responders/non-responders) analyses in Checkpoint and Adaptive immune signaling networks

In the above analyses, we have used all markers available on the corresponding FACS panels, with all immune cell types and subtypes pooled together. To achieve a higher analysis granularity, we have further separated each FACS dataset into four subsets, according to the four major immune cell types: (1) Naïve CD4+ T cells, (2) Naïve CD8+ T cells, (3) Non-naïve CD4+ T cells, (4) Non-naïve CD8+ T cells. These were stratified using CD4, CD8, CCR7 and CD45RA markers, according to the following membership rules: (1) 1011, (2) 0111, (3) 1010+1001+1000, (4) 0110+0101+0100, where “1” means high, and “0” – low marker concentration of CD4, CD8, CCR7 and CD45RA, in that order. The cutoff points between “high” and “low” values of compensated immunofluorescence intensity were defined separately for the Checkpoint and Adaptive networks (due to different calibrations) and were as follows: Checkpoint: CD4=1500, CD8=750, CCR7=150, CD45RA=400; Adaptive: CD4=1000, CD8=1000, CCR7=150, CD45RA=1000.

Subsequently, we have generated BNs for 32 sub-datasets: (responders/non-responders) x (before/after immunotherapy) x (Checkpoint / Adaptive) x (Naïve CD4+ T cells / Naïve CD8+ T cells / Non-naïve CD4+ T cells / Non-naïve CD8+ T cells). Markers CD4, CD8, CCR7 and CD45RA were naturally excluded from these networks. 8-bin discretization was used throughout, to increase specificity (decrease edge density in the networks). Comparisons of the resulting networks allowed us to evaluate changes in immune signaling networks in responders and non-responders before (day 1) and after (day 21) immunotherapy, stratified by a major immune cell type. In order to quantify and automate these comparisons, we have introduced “indicator/contrast variables”, as illustrated in the following example (Checkpoint network, Non-naïve CD4+ T cells sub-dataset):

Figure 4 depicts four BNs obtained using 8-bin equal-frequency discretization --- (a) day 1, non-responders only, (b) day 1, responders, (c) day 21, non-responders, (d) day 21, responders.

**Figure 4.**
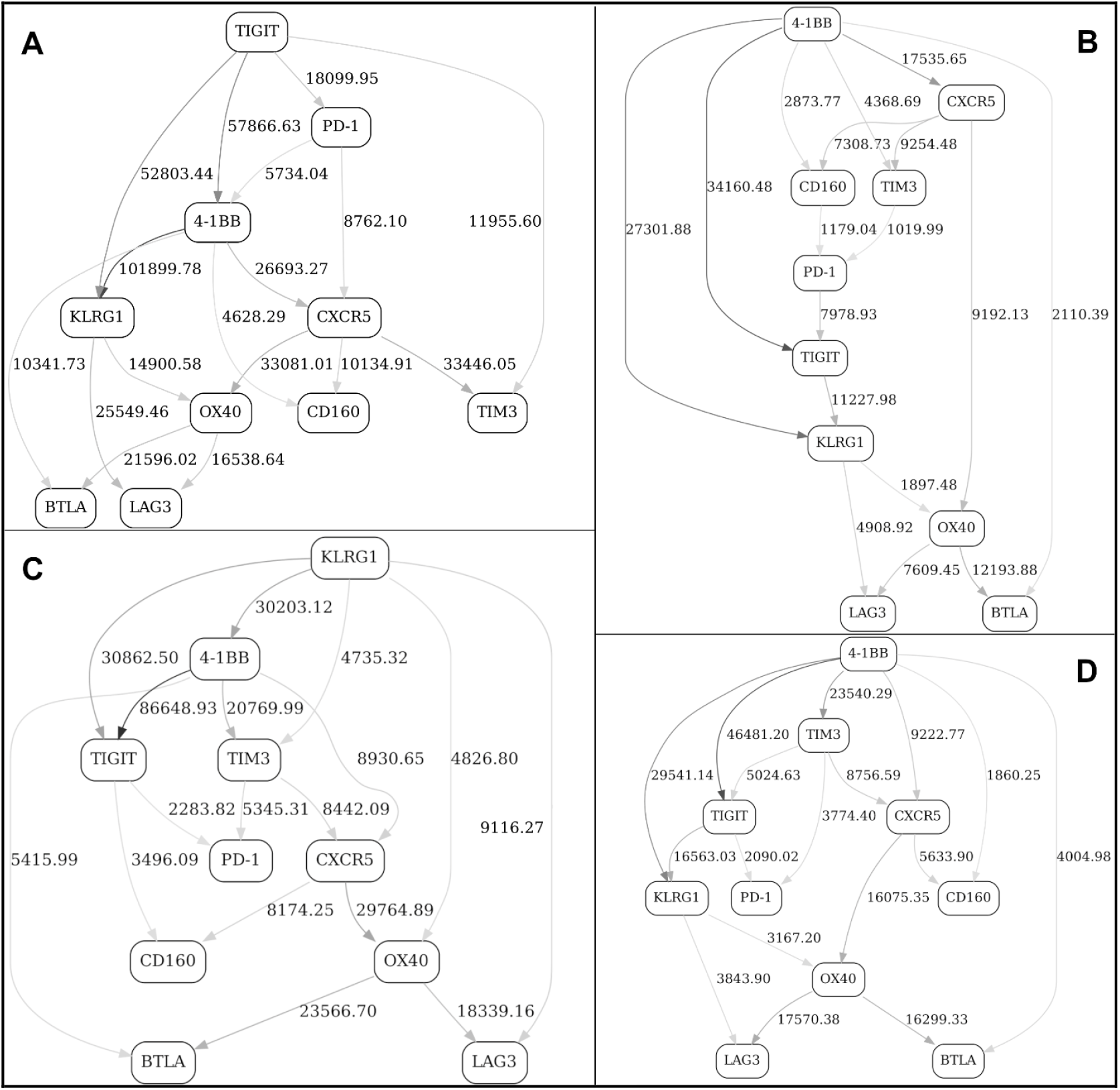
BNs constructed from FACS data (Checkpoint immune signaling network panel, Non-naïve CD4+ T cells sub-dataset) from 13 patients. (a) day 1, non-responders only, (b) day 1, responders, (c) day 21, non-responders, (d) day 21, responders. See Figure 1 legend for further details.

At this stage, it is difficult to efficiently compare/contrast these networks in order to tease out the most significant changes, short of manually enumerating the topological differences. Therefore, we have introduced a “Contrast” variable with four distinct states (day 1 / non-response, day 1 / response, day 21 / non-response and day 21 / response) and included it in the pooled, or “supergraph” BN (Figure 5). In Figure 5, four markers (KLRG1, BTLA, OX40 and, to a lesser extent, PD-1) were observed in the MN of the “Contrast” variable (red node in the graph), suggesting that it is the behavior of these four markers (reflected in the corresponding MNs in the original four stratified networks, Figure 4abcd) that differentiate the responder / non-responder / day 1 / day 21 states.

**Figure 5.**
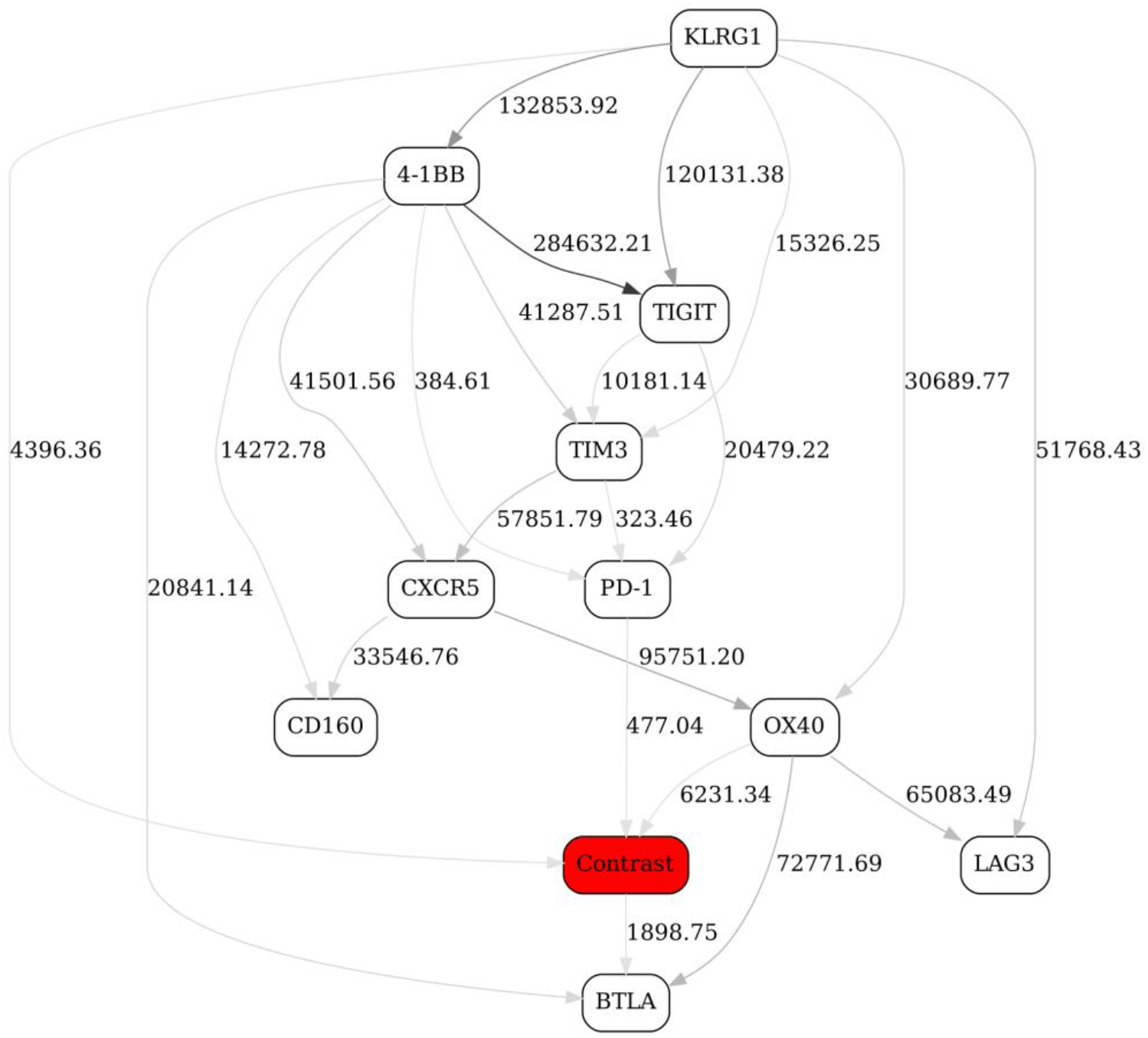
A BN constructed from FACS data (Checkpoint immune signaling network panel, Non-naïve CD4+ T cells sub-dataset) from 13 patients. The “Contrast” node is the indicator variable with four states, (day 1 / non-response, day 1 / response, day 21 / non-response and day 21 / response). See Figure 1 legend for further details.

To look even deeper, we have subsequently introduced two separate contrast variables (“Response” and “Day”, red nodes in the graph), and constructed four contrast networks shown in Figure 6: (a) at day 1, responders vs. non-responders, (b) at day 21, responders vs. non-responders, (c) non-responders, at day 1 vs. day 21, and (d) responders, at day 1 vs. day 21.

**Figure 6.**
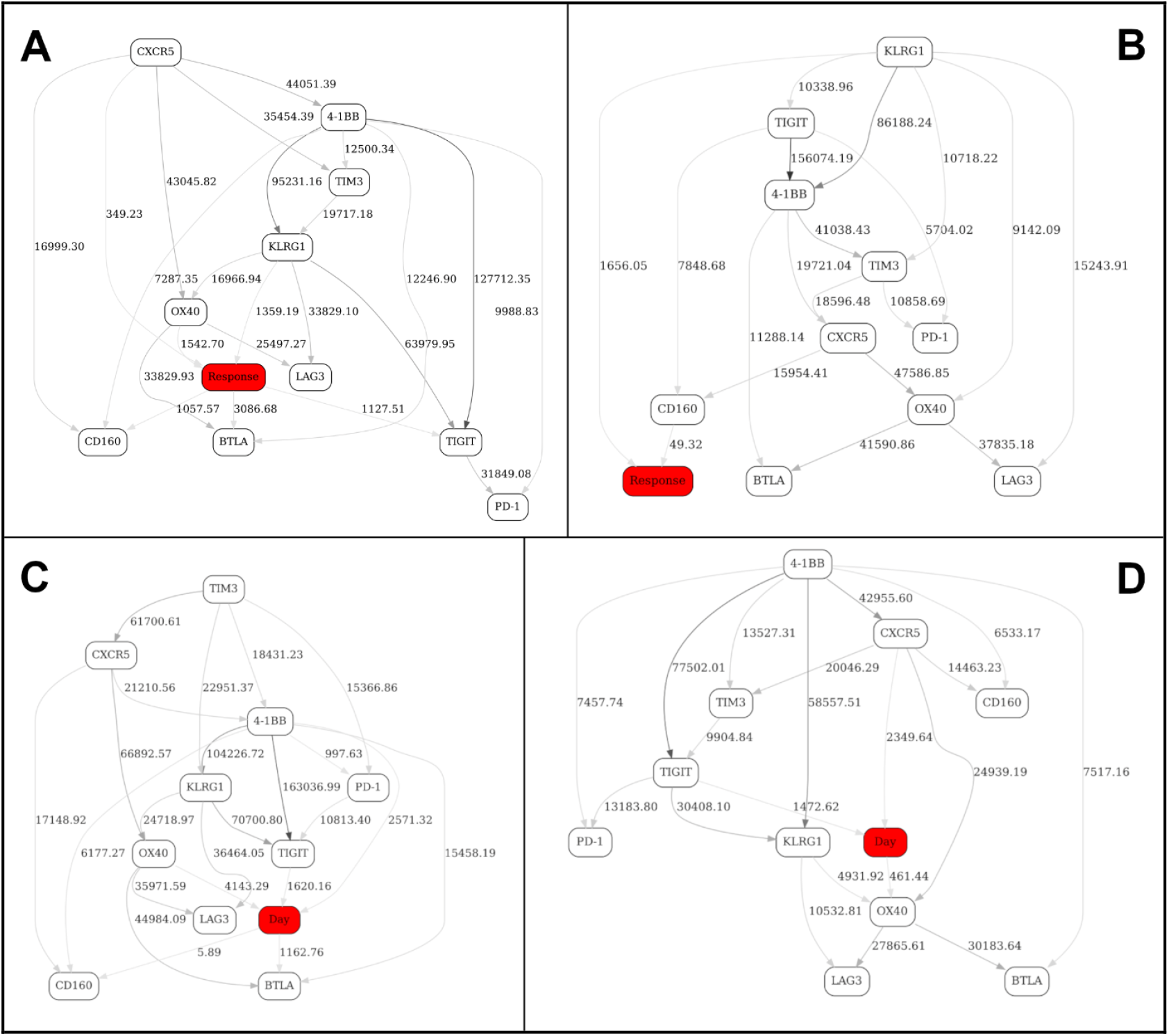
Contrast BNs constructed from FACS data (Checkpoint immune signaling network panel, Non-naïve CD4+ T cells sub-dataset) from 13 patients. (a) at day 1, responders vs. non-responders, (b) at day 21, responders vs. non-responders, (c) non-responders, at day 1 vs. day 21, and (d) responders, at day 1 vs. day 21. “Day” node is the indicator variable with two states (day 1, day 21). “Response” node is the indicator variable with two states (response, non-response). See Figure 1 legend for further details.

We were especially interested in Figure 6b that compares/contrasts responders and non-responders at day 21 (post-treatment). It appears that there is just one marker with a strong network edge in the MN of the “response” variable, KLRG1. By looking at KLRG1’s MNs in the stratified networks (Figure 4cd), we conclude that the major difference between the Checkpoint network in responders and non-responders at day 21 (after immunotherapy treatment) is a relatively less prominent dependency (correlation) between KLRG1 and TIGIT, and an absence of dependency between KLRG1 and TIM3 in responders (compared to non-responders). This is an example of an interaction, and change thereof, that would have been exceedingly difficult to pinpoint without the BN comparison analysis along the above lines. In other words, our analytical strategy automatically identified the single most important topological difference between the immune signaling networks reflecting two different states (responders vs. non-responders after immunotherapy). (We need not, of course, concentrate on only the one top difference.)

The results of remaining seven sub-dataset analyses (that is, Adaptive immune signaling network panel x Non-naïve CD4+ T cells sub-dataset, and both Adaptive and Checkpoint panels x Naïve CD4+ T cells, Naïve CD8+ T cells and Non-naïve CD8+ T cells sub-datasets) are shown in S1 Appendix (together with the itemized figure captions/legends). To illustrate another case of a “contrast” network analysis, consider, for example, S1 Figure 52, which contrasts Adaptive Non-naïve CD8+ T cells networks at day 1 between responders and non-responders. The strongest network edge in the MN of the “response” variable points to ICOS, and after looking at the ICOS node’s MNs in the corresponding stratified networks (S1 Figure 46 - day 1, responders, and S1 Figure 47 - day 1, non-responders) we conclude that there are interactions between ICOS and other markers (notably, CD127) that are present in responders but not in non-responders. Again, just like in a previous example, we were able to automatically single out one strongest interaction (in this case, between ICOS and CD127) that distinguishes between the two different network states.

The above analyses illustrate the process of identifying markers and also higher-order marker interactions that distinguish immune signaling networks in responders and non-responders before and after immunotherapy treatment. The more theoretical question remains --- does the “contrast variable” approach comprehensively capture the topology changes between the networks being compared? In a BN, a presence of an edge is nothing more than a presence of a node in the parent set of another node. Any time an edge is introduced, we are in fact looking at the corresponding change in the joint distribution of the two nodes in question: therefore, the “contrast” node is necessarily dependent on the joint distribution of the two nodes that gain or lose an edge. Consequently, appearance / disappearance of an edge A->B is reflected in that the contrast C gets “pulled in” as A&B -> C. This is because, in order for contrast C to be independent, we must have P(C | A&B) = P(C). However, P(A&B | C) P(C) = P(A&B) P(C | A&B), and so P(C | A&B) = P(A&B | C) P(C) / P(A&B). Thus, the only way C can be independent of A&B is if P(A&B | C) = P(A&B), which we know is not true because it is given that P(A&B | C=0) ≠ P(A&B | C=1). Therefore, the variations that are not “detected” by contrast variable must be either noise or structural variants from the same BN equivalence class. Moreover, the appearance/disappearance of an edge is not the only inter-BN difference that will be detected *via* contrast variables. For example, the same edge could represent a direct or an inverse relationship in two graphs, so one would not necessarily know the difference by just looking at the edges, but it would be a distribution difference that would be captured by a contrast variable all the same. In fact, all differences associated with going from one graph to another are necessarily nothing but results of conditioning on the contrast variable in the supergraph --- for example, a non-response graph G0 (Figure 4c) is simply a supergraph G (Figure 6b) conditioned with contrast C (“Response”) = 0. Therefore, we can expect that all differences will necessarily be encapsulated by the contrast variable connectedness.

### 2.3. Validation of the BNs using statistical criteria, and comparison of the BN results with other multivariate analysis methods

While the BNs constructed in this study (Figures 1-6, S1 Figures 1-63) are informative in a sense that (i) they suggest the markers most likely to influence the response status and immune network changes and (ii) they can be compared/contrasted among themselves, it is difficult to directly assign the measure of statistical confidence to a particular edge strength in the network. Conversely, it would be instructive to compare the BN analysis with other multivariate analysis methods --- namely, multivariate classifiers, and variable selection (importance ranking) methods. In this section, we will illustrate BN validation and comparison with alternative multivariate analysis methods using one particular BN example --- responders vs. non-responders at day 1, Adaptive immune signaling network panel, naïve CD4+ T cells (Figure 7, same as S1 Figure 43). Our interest in this specific subset (naïve CD4+ T cells) was heightened by the observations that, according to the BN analysis, CXCR3 appeared to be a very strong response predictor in naïve CD4+ T cells (Figure 7), and also by the recent literature suggesting an important role of CXCR3 in response to immunotherapy [28, 29 and references therein]. (All standard machine learning and statistical analyses in this section were carried out using Scikit-learn machine learning toolkit [30], with default parameters unless otherwise noted.)

**Figure 7.**
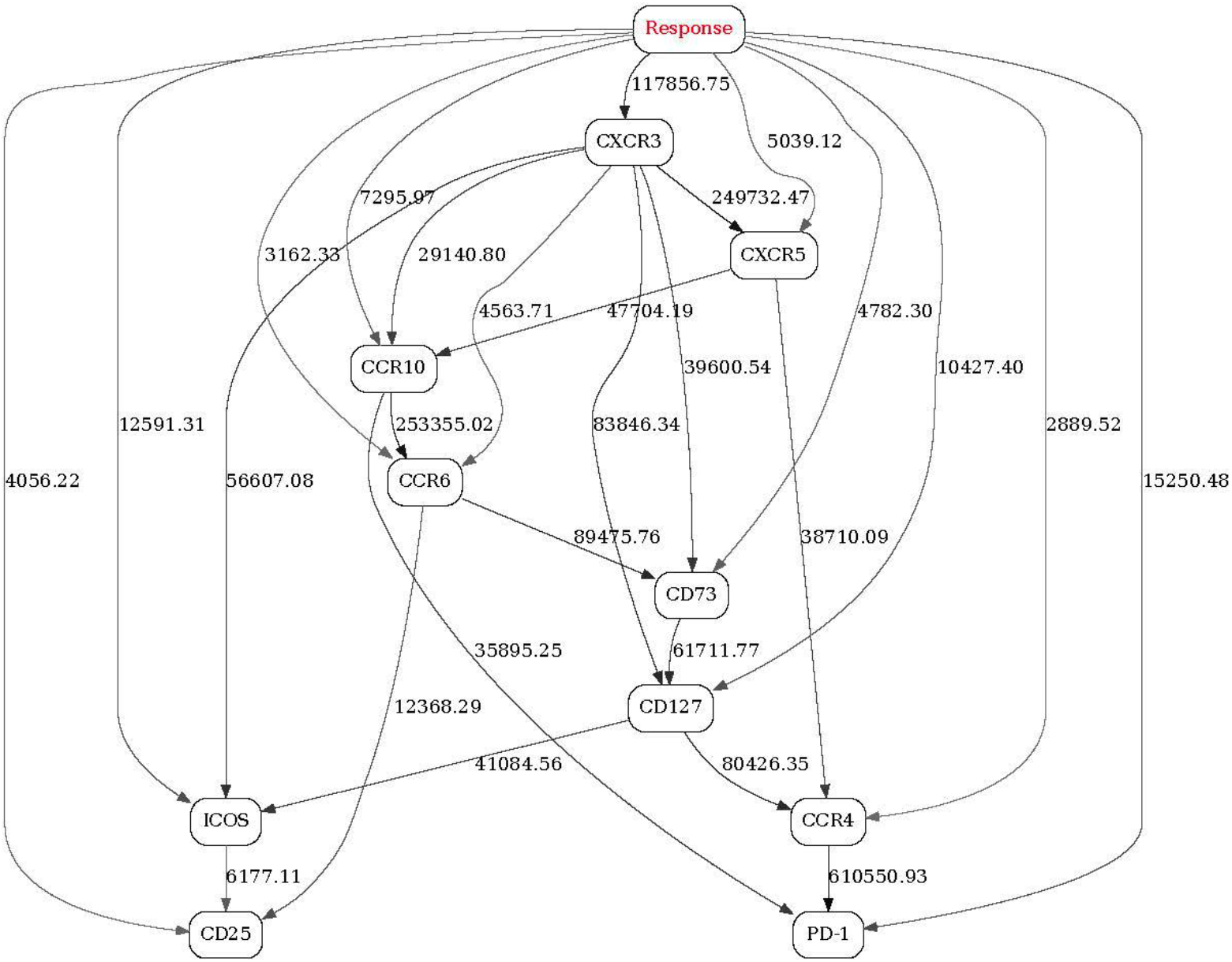
Contrast BN constructed from FACS data (Adaptive immune signaling network panel, Naïve CD4+ T cells) from 13 patients pre-immunotherapy treatment (day 1). “Response” (red node in the graph) is the indicator variable with two states (response, non-response). See Figure 1 legend for further details.

In order to quantify, in a statistically objective fashion, the association strength between CXCR3 (as wel,l as other nine markers, see Figure 7) and Response status, we used univariate logistic regression, point-biserial correlation, and distribution distance metrics. Application of univariate logistic regression and point-biserial correlation (Table 4) did not show significant differences between CXCR3 and other markers in their predictive ability with respect to Response status. Similarly, results from the FlowJo manual gating analysis [27] showed no indication that CXCR3 was differentially expressed in the naïve CD4+ T cells between responders and non-responders at day 1 (Figure 8).

**Table 4.** Univariate logistic regression and point-biserial correlation results. (Adaptive immune signaling network panel, Naïve CD4+ T cells, responders vs non-responders at day 1). Classification accuracy was assessed using 10-fold cross-validation.

| Variable | Logistic regression classification accuracy (responders *vs* non-responders) | Point-biserial correlation with response status |
| --- | --- | --- |
| CXCR3 | 68.34% | 0.32 |
| CCR10 | 67.80% | -0.12 |
| CD73 | 67.50% | 0.12 |
| CCR6 | 67.72% | 0.05 |
| CD25 | 67.82% | -0.15 |
| ICOS | 67.80% | 0.06 |
| CXCR5 | 67.67% | 0.10 |
| PD-1 | 69.12% | -0.21 |
| CD127 | 67.80% | 0.26 |
| CCR4 | 67.96% | -0.18 |
| Response | 100.00% | 1.00 |

**Figure 8.**
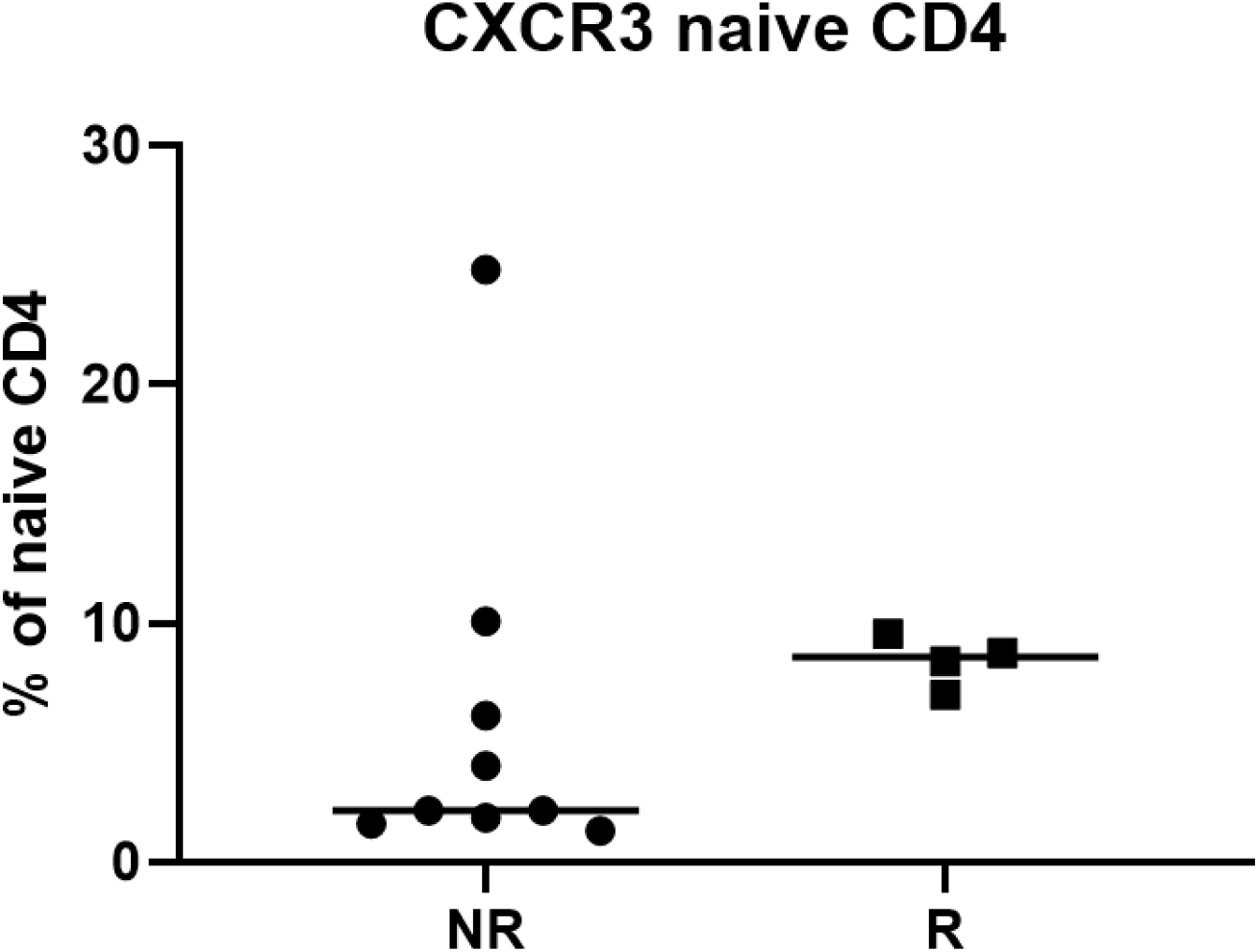
“Traditional” flow cytometry assessment of CXCR3 expression. Naïve CD45RA+ CD4+ CD3+ T cells were manually gated using FlowJo flow cytometry analysis software. The fraction (%) of naïve CD4+ T cells manually judged to express CXCR3 across patient responders (R) and non-responders (NR) are shown. Unpaired two-tailed t-test resulted in p-value of 0.5496, indicating no significant difference between responders and non-responders.

Subsequently, we used Earth Mover’s Distance (EMD) [31] and the Energy Distance (ED) [32] distribution distance metrics to compare differences in effective cumulative distribution functions (ECDFs) of the CXCR3 and other nine markers’ fluorescence intensity values between responders and non-responders. Previously, EMD has been shown to be objective and unbiased in comparing biomarker expression levels in cell populations [33]. As can be seen in Figure 9 and Table 5, CXCR3 stands out as the strongest, by far, discriminator between responders and non-responders. (To provide the sense of scale, EMD is roughly equal to the area between the two ECDFs --- therefore, a difference between, for example, EMD of 55.660 and 22.011, as shown in Table 5, is highly significant; similarly, ED is a linear function of the Cramer Distance, with the same scale sensitivity). Figure 10 further illustrates the significant difference between the probability distribution functions (PDFs) of CXCR3 fluorescence intensity values in responders and non-responders, in contrast with the “traditional” FlowJo result output (Figure 8).

**Table 5.**
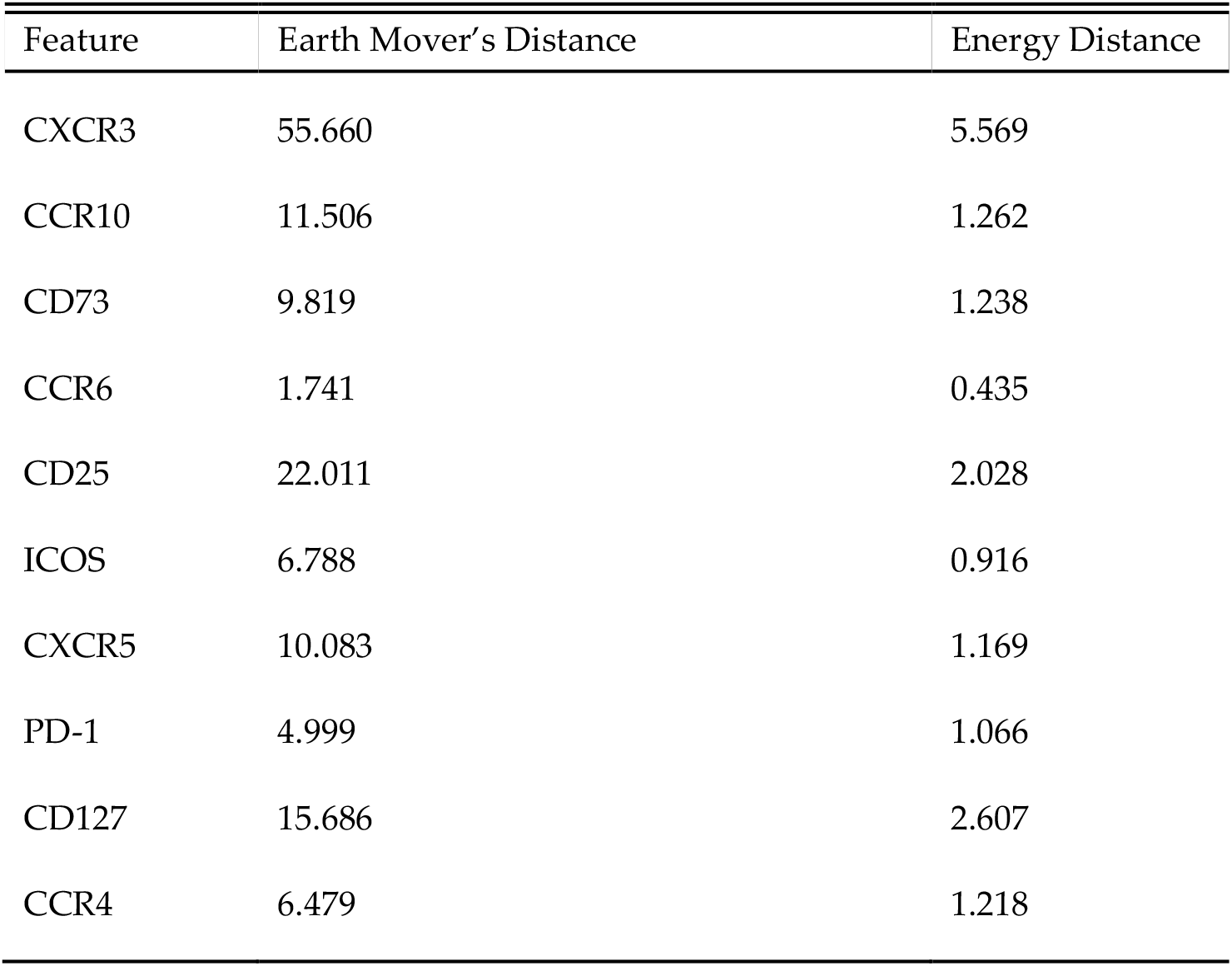
EMD and ED measures (computed from the ECDFs shown in Figure 9) between responders and non-responders for CXCR3 and other nine markers. (Adaptive immune signaling network panel, Naïve CD4+ T cells, day 1).

| Feature | Earth Mover's Distance | Energy Distance |
| --- | --- | --- |
| CXCR3 | 55.660 | 5.569 |
| CCR10 | 11.506 | 1.262 |
| CD73 | 9.819 | 1.238 |
| CCR6 | 1.741 | 0.435 |
| CD25 | 22.011 | 2.028 |
| ICOS | 6.788 | 0.916 |
| CXCR5 | 10.083 | 1.169 |
| PD-1 | 4.999 | 1.066 |
| CD127 | 15.686 | 2.607 |
| CCR4 | 6.479 | 1.218 |

**Figure 9.**
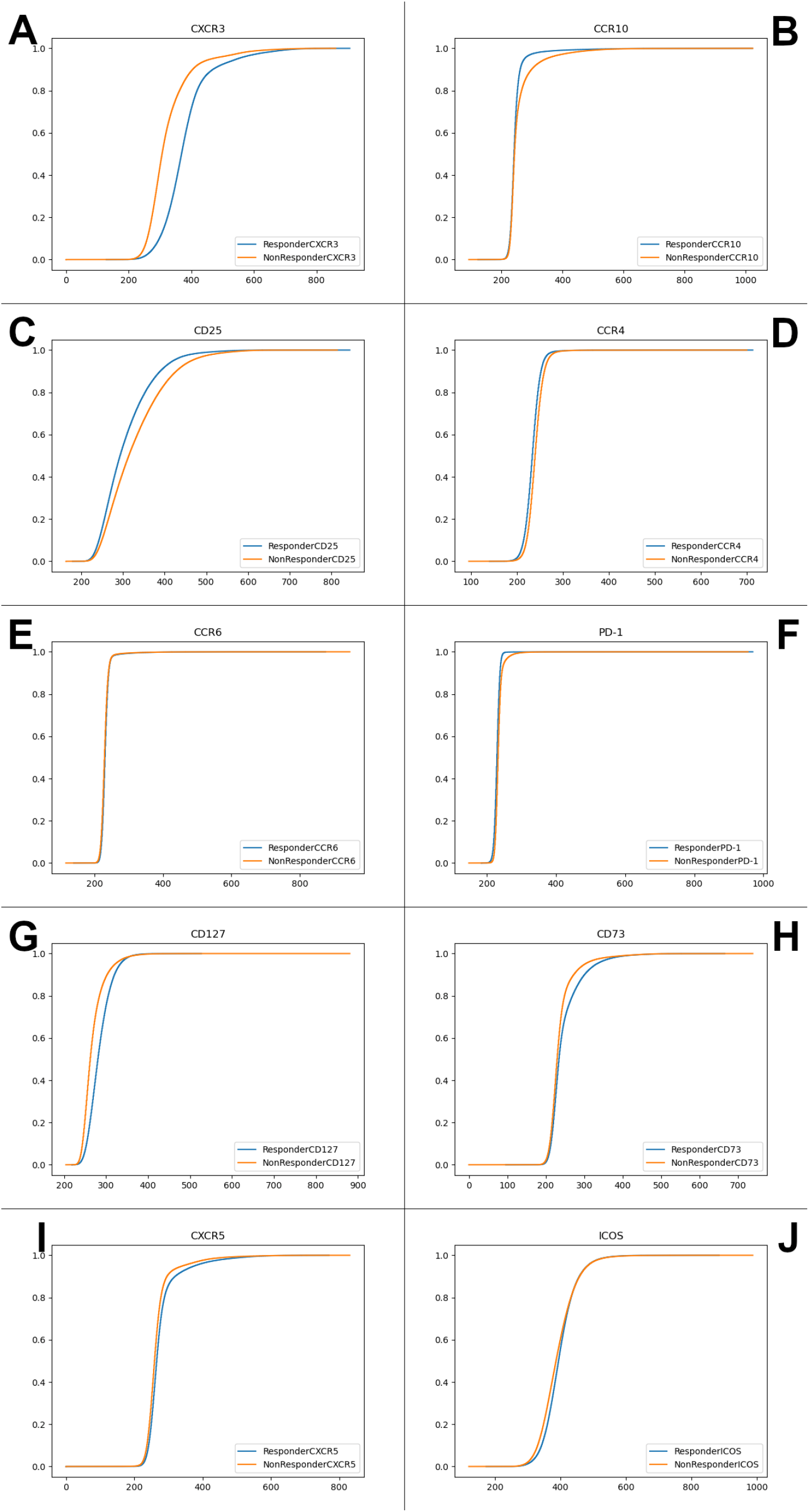
ECDFs for CXCR3 (Figure 9A) and other nine markers (Figure 9B-J) compensated fluorescence intensities, compared in responders and non-responders. (Adaptive immune signaling network panel, Naïve CD4+ T cells, day 1).

**Figure 10.**
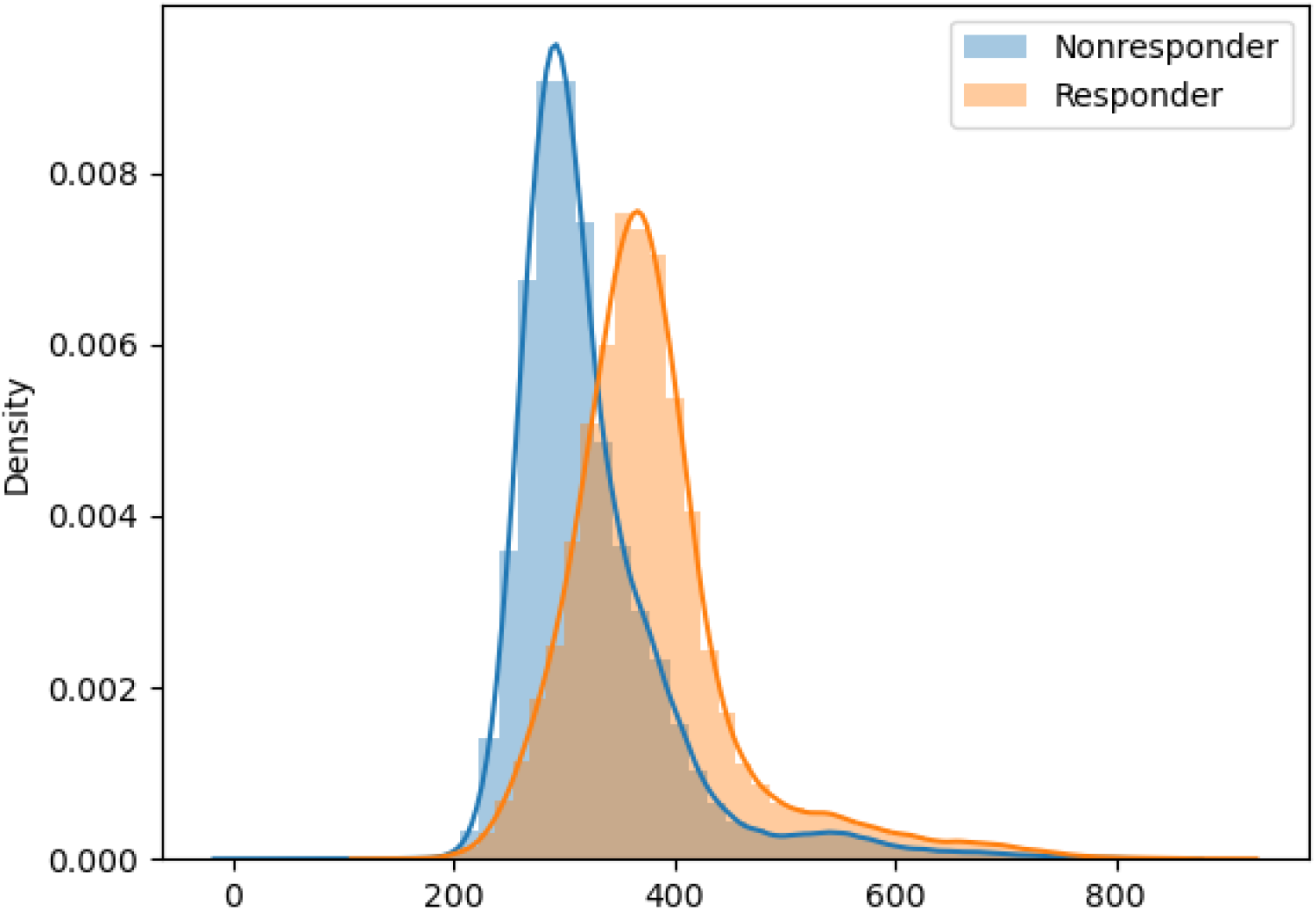
PDFs for CXCR3 compensated fluorescence intensities, compared in responders and non-responders. (Adaptive immune signaling network panel, Naïve CD4+ T cells, day 1).

Clearly, there is a conflict. Notable discrepancy between BN and EMD/ED/distributional results (strong CXCR3 – Response association) on the one hand, and FlowJo results (no noticeable CXCR3 – Response association) on the other hand can be attributed to the fact that the former methods capture (via multi-bin discretization with BNs, and full-distribution assessment with EMD/ED) the information that manual gating and simpler statistical analyses simply do not “see”. To further investigate this, we applied select multivariate classifiers to the same data that was used to build the BN in Figure 7. First, we tested various generalized linear models implementing different kinds of regularization and solvers. CXCR3 and, to a lesser extent, CCR10 appeared to be the strongest signals (Table 6). We followed up with the Random Forests (RF) ensemble classifier augmented by the Shapley value-based explainer (TreeSHAP) [34], to look at both individual feature importance and pairwise feature importance. The results (Figure 11) suggested that CXCR3 was indeed the most significant single predictor for response. Additionally, its significance showed partial dependence on the value of CCR10, the same interplay that was observed in the BN (Figure 7).

**Table 6.** Application of different variants of generalized linear models (Adaptive immune signaling network panel, Naïve CD4 cells, responders vs non-responders at day 1). “lbfgs” stands for Limited-memory Broyden–Fletcher–Goldfarb–Shanno algorithm; “liblinear”, coordinate descent algorithm; L1, L1 regularization; L2, L2 regularization. Classification accuracy was assessed using 5-fold cross-validation. Extended decimals are shown to indicate that the results were similar but not identical.

| Model | Classification accuracy | Coefficients, in order of: ['CXCR3', 'CCR10', 'CD73', 'CCR6', 'CD25', 'ICOS', 'CXCR5', 'PD-1', 'CD127', 'CCR4']] - [Intercept] |
| --- | --- | --- |
| L2, lbfgs logistic regression | 79.8128% | [[ 1.3640, -1.0545, 0.3220, 0.1558, -0.2632, -0.2438, -0.6944, -0.5492, 0.4632, -0.2202]] [-1.0805] |
| L2, liblinear logistic regression | 79.8107% | [[ 1.3641, -1.0547, 0.3220, 0.1558, -0.2632, -0.2438, -0.6943, -0.5492, 0.4633, -0.2202]] [-1.0806] |
| L1, liblinear logistic regression | 79.8102% | [[ 1.3636, -1.0542, 0.3219, 0.1557, -0.2631, -0.2437, -0.6939, -0.5591, 0.4632, -0.2201]] [-1.0803] |

**Figure 11.**
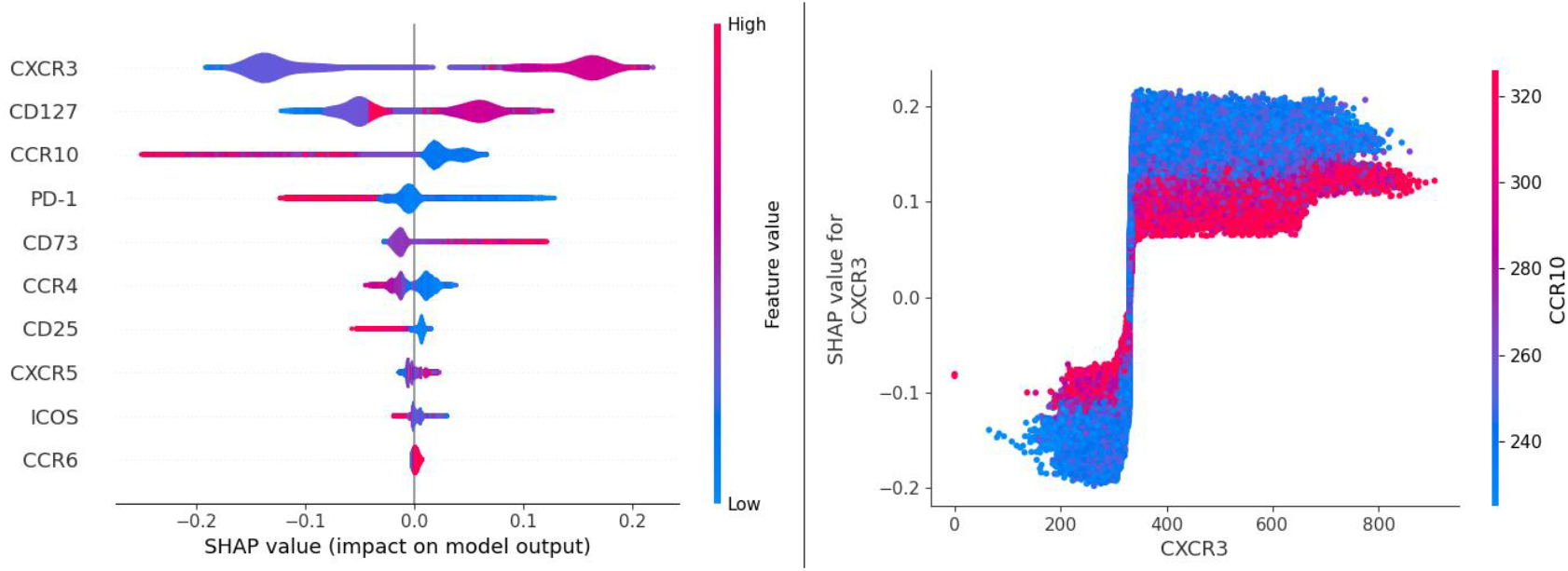
SHAP (Shapley Additive exPlanations) output of the RF model (Adaptive immune signaling network panel, Naïve CD4 cells, responders vs non-responders at day 1). The left panel shows a set of “beeswarm” plots reflecting individual variable importance (variables are ranked in descending order) and the impact on the model output (probability of response) depending on the value of the feature (marker). For instance, high values of CXCR3 (red color) act as a positive predictor (right side of the central axis) of response. The right panel shows an example of the interplay between two features, here CXCR3 and CCR10 (low values of CCR10 increase the predictive power of CXCR3). CCR10 was found to be the strongest modulator of CXCR3.

To summarize, BN, multivariate logistic regression and RF suggested that the CXCR3 was the strongest predictor for response. This was supported by the distribution distance metrics. On the contrary, “traditional” FlowJo data analysis workflow did not detect this signal. Finally, BN and RF analyses also provided an added value of capturing pairwise and higher-level interactions, although BN modeling was superior in interpretability.

## 3. Discussion

In this study, we developed a novel secondary data analysis strategy aimed at FACS data in the immuno-oncology context. It is based on BN modeling, maximum entropy discretization, multi-stage marker configuration frequency sorting, and pooling sub-dataset BNs while introducing “contrast” variables. The end output of our computational pipeline consists of the following components: (a) a compact subset of potentially predictive (with respect to a clinical outcome or some other phenotype) markers, typically in the single digit (no more than 4-8 markers) range, (b) a graphical causal model of the interactions between these predictors and other variables, together with the corresponding relative interaction strengths, (c) a compact list of the most frequent marker combinations, together with the corresponding outcome/phenotype probabilities, and (d) a shortlist of markers whose behavior (as reflected in their corresponding MNs in the stratified sub-networks) is significantly different between the varying immune signaling network states (responders / non-responders, before / after immunotherapy). These markers and combinations can subsequently be associated with the immune cell types.

One advantage of this analytical strategy is that it is fully automated and generalizable to other similar datasets, of varying data structure and dimensionality. At the same time, it allows for certain amount of flexibility in the analysis process. First, different discretization approaches can be tried, depending on the investigators’ understanding of what “high” and “low” might mean for a particular marker concentration. Second, more (or fewer) potential predictors can be selected from the outcome/phenotype variable’s MN, depending on the BN’s local sparseness and the general pattern of conditional independencies in the network. (This, however, is a complicated issue, and we plan to investigate it further as part of our ongoing BN methodological research.) Third, more (or fewer) potential predictor configurations (and corresponding cell types) can be scrutinized at later analysis stages, depending, for example, on the size and coverage/purpose of the concrete FACS panel, or on whether investigators are interested in rare cell types that are very strongly associated with particular outcomes/phenotypes.

There are two additional interrelated methodological caveats that are intrinsic to the BN modeling in general: dependency strength validation, and inter-network comparisons. Throughout this study, we use edge strengths to estimate statistical support for the networks and their features. Edge strength is conceptually similar to the marginal likelihood ratio test, in that it is proportional to the ratio of the BN model scoring function value when the edge is present in the BN to the BN model scoring function value when the edge is absent in the BN. Edge strengths do not directly translate into p-values; therefore, once a particular dependency/association is suggested by the BN, and is of interest to the investigators, one should follow up with “traditional” statistical tools. This said, edge strengths are directly comparable to each other within a single BN. However, it is not immediately clear whether they are directly comparable to each other *between* the BNs, even if the corresponding datasets are similar in their dimensionality. There are reasons to believe that edge strengths are linear functions of the dataset size (e.g., number of cells in one sample in the FACS context), but this aspect requires further investigation. For now, we would not necessarily advise comparing edge strengths between the networks (e.g., between Figures 1, 2, 3 in this study) unless the BN-generating datasets are very similar in their size and structure. Instead, when possible, the data should be pooled, and indicator/contrast variables (distinguishing the networks to be compared) introduced (e.g., Figures 5, 6 in this study).

One somewhat unexpected result of our study was a clear divergence between two groups of analytical approaches --- FlowJo and simpler analysis methods *vs* BNs and multivariate classifiers. Using one example dataset (Adaptive immune signaling network panel, Naïve CD4+ T cells, responders vs non-responders at day 1) we have demonstrated that there exists a strong predictive signal, CXCR3, that is reliably “picked up” by the latter, but not by the former. Since many FACS analyses in current immunology research and recent literature rely on the FlowJo data analysis pipeline, we are concerned that strong signals (such as CXCR3 in our example) might be missed. This could be attributed to the intrinsic limitations of the manual gating process [4, 9], including such interrelated factors as (i) subjectivity of the manual gating procedure, (ii) discretization in two bins, (iii) loss of the higher-order information, and (iv) linearity assumptions throughout. Therefore, we suggest that in the future investigators should augment the “standard” FlowJo (or similar manual gating) data analysis pipeline with the direct analysis of the compensated fluorescence intensity data, preferably by a variety of methods, and preferably including at least some non-linear ones. In that, our conclusions dovetail with other recent recommendations in the field [33].

One of the obvious limitations of this study was the low number of patients (14). However, the primary purpose of the study was to propose and develop a novel analytical framework, and to see if it compares favorably with the existing tools. Indeed, our analysis pipeline has successfully identified a number of potentially strong predictive biomarkers (and their combinations), including one biomarker (CXCR3) that was clearly missed by the more conventional methodology.

It should also be emphasized that the principal advantage of the BN + MN approach is the inherent interpretability of the analysis results, in which it compares favorably with even the relatively sophisticated methods such as RF (although the recent progress in increasing ensemble classifiers’ interpretability is very promising [34]).Conversely, the BN + MN approach can be more constructive than the commonly used dimensionality reduction / clustering / visualization techniques, in that (i) it presents the structures (full networks and MNs) that are directly mechanistically interpretable, in contrast to the more vaguely defined similarity “clusters” and “trees”, and (ii) it allows to immediately identify, rank and prioritize the important features within such structures. This is not to diminish the value of such dimensionality reduction methods; rather, the BN + MN approach presents an alternative, and oftentimes more natural and straightforward, path from the “pretty pictures” to the actual candidate markers and their interactions. In general, we agree with the recent literature [38] in that while data visualization and clustering algorithms are helpful with the broad exploration of the flow cytometry data, it is the novel supervised machine learning techniques that hold the most promise for a seamless automatization of the analysis of large flow cytometry datasets. This becomes even more relevant in light of the ongoing advent of increasingly higher-resolution (e.g., 40-color) flow cytometry panels [39]. By its very design, our “BN => MN funnel” approach is easily scalable to hundreds of parameters.

In conclusion, since only some cancer patients tend to be responsive to immunotherapy treatments such as ICB, development of reliable predictive biomarkers for treatment response is urgently needed to select patients. While recent findings have shown some promise in tumor tissue-based immunological biomarkers [14], peripheral blood biomarkers are more accessible and less invasive [17]. In this study, peripheral blood FACS data, combined with our analytical strategy, have been successfully used to suggest such biomarkers, and gain insight into their interactions and dynamics before and after treatment. Presently, we plan to validate and follow up immune markers (and combinations / interactions thereof) associated with clinical responses that were identified by us in this study, starting with CXCR3 and CXCR3-CCR10 interaction. In the future, we intend to generalize and apply our analysis tools to other cancer types, treatments, and patient cohorts.

## 4. Materials and Methods

### 4.1. Flow cytometry

Peripheral blood mononuclear cells (PBMCs) from patients consented to an IRB-approved protocol were isolated from heparinized blood by Ficoll-Paque density centrifugation and cryopreserved in FSB with 10% DMSO. Cryopreserved PBMCs were thawed and stained with antibodies for the following flow cytometry panels:

Checkpoint panel: CD4, CD8, CD45RA, KLGR1, CCR7, CXCR5, 4-1BB, BTLA4, LAG3, OX40, CD160, TIGIT, PD1, TIM3;

Innate panel: CD3, CD14, CD16, CD20, CD33, CD56, CD11c, CD141, CD1c, CD123, CD83, HLA-DR, TCRgδ, PD-L1;

Adaptive panel: CD4, CD8, CCR10, CCR6, CD73, ICOS, CXCR3, CXCR5, CD45RA, CCR4, CCR7, CD25, CD127, PD1.

Flow cytometry was performed using Fortessa Flow Cytometers and flow cytometry data was analyzed using FlowJo software (BD Biosciences) [27].

### 4.2. Bayesian networks modeling

BN modeling is a systems biology method that constructs a graphical visualization of a joint multivariate probability distribution of the random variables in the dataset.

While recent BN modeling software implementations are highly scalable [35], combining scalability with mixed variable types (e.g., continuous and discrete) is less straightforward; an argument can be made [35,36] that discretization of continuous variables is a better solution, in practical terms, than imposing mixed distribution models. Therefore, in this study we were using full sample equal-frequency discretization which also allowed us to adjust the under/over-fitting balance (i.e., specificity vs. sensitivity). Typically, discretizing into a smaller number of bins (2-3) increases sensitivity (higher edge density), and discretizing into a larger number of bins (4-8) increases specificity (lower edge density). Our BN implementation [35] is based on a hybrid “sparse candidates” + “search-and-score” algorithm with random restarts until convergence. In the resulting network structures, we would typically “zoom in” on the immediate Markov neighborhood (MN) of a specific variable (such as an immune system-related marker, or a patient’s clinical response status), which corresponds to all the nodes (variables) directly linked to a specific variable in question. (MN is a simplified version of the Markov Blanket --- the latter takes into account parent-offspring relationship directionality --- meaning, for the practical purposes, that “two degrees of separation”, or more, are sometimes needed for fully assessing the conditional independence relationships around a specific variable/node.) Numbers next to the edges and edge “thickness” in the resulting BN figures specify relative edge strengths (which are marginal likelihood-based). Further details on the BN modeling in general and our implementation thereof can be found in [35,37].

## Supplementary Materials

### Appendix S1

Bayesian networks obtained from the remaining sub-datasets in the present study (i.e., Adaptive immune signaling network panel x Non-naïve CD4 sub-dataset, and both Adaptive and Checkpoint panels x Naïve CD4, Naïve CD8 and Non-naïve CD8 sub-datasets).

S1 Figures 1-9 -- Checkpoint, Naïve CD4;

S1 Figures 10-18 – Checkpoint, Non-naïve CD8;

S1 Figures 19-27 – Checkpoint, Naïve CD8;

S1 Figures 28-36 – Adaptive, Non-naïve CD4;

S1 Figures 37-45 – Adaptive, Naïve CD4;

S1 Figures 46-54 – Adaptive, Non-naïve CD8;

S1 Figures 55-63 – Adaptive, Naïve CD8.

Within each group, the nine networks are as follows:

Day 1, responders

Day 1, non-responders

Day 21, responders,

Day 21, non-responders,

Day 1 and day 21, responders, with “day” contrast variable,

Day 1 and day 21, non-responders, with “day” contrast variable,

Day 1, responders and non-responders, with “response” contrast variable,

Day 21, responders and non-responders, with “response” contrast variable, “Supergraph” (all data pooled together, with a four-state “contrast” variable).

## Data Availability Statement

Principal datasets analyzed in this study have been deposited to Dryad (doi:10.5061/dryad.fxpnvx0ng). https://doi.org/10.5061/dryad.fxpnvx0ng

All intermediate/auxiliary datasets will be made available by the authors, without undue reservation, to any qualified researcher.

## Acknowledgments

The authors are grateful to Arthur D. Riggs and Sergio Branciamore for many stimulating discussions and useful suggestions.

## Author Contributions

**Andrei S. Rodin**; Conceptualization, formal analysis, funding acquisition, investigation, methodology, project administration, resources, supervision, visualization, writing (original draft preparation). **Grigoriy Gogoshin**; Conceptualization, data curation, formal analysis, investigation, methodology, software, visualization, writing (original draft preparation). **Seth Hilliard**; Conceptualization, data curation, formal analysis, investigation, methodology, software, visualization, writing (original draft preparation). **Lei Wang**; Conceptualization, data curation, formal analysis, investigation, methodology, writing (original draft preparation). **Colt Egelston**; Conceptualization, data curation, formal analysis, investigation, methodology. **Russell C. Rockne**; Conceptualization, funding acquisition, methodology, visualization, writing (review and editing). **Joseph Chao**; Conceptualization, data curation, funding acquisition, investigation, project administration, resources, writing (original draft preparation). **Peter P. Lee**; Conceptualization, funding acquisition, investigation, methodology, project administration, resources, supervision, visualization, writing (original draft preparation).

