## Supplementary material for "Dissecting Response to Cancer Immunotherapy by Applying Bayesian Network Analysis to Flow Cytometry Data": S1 Appendix

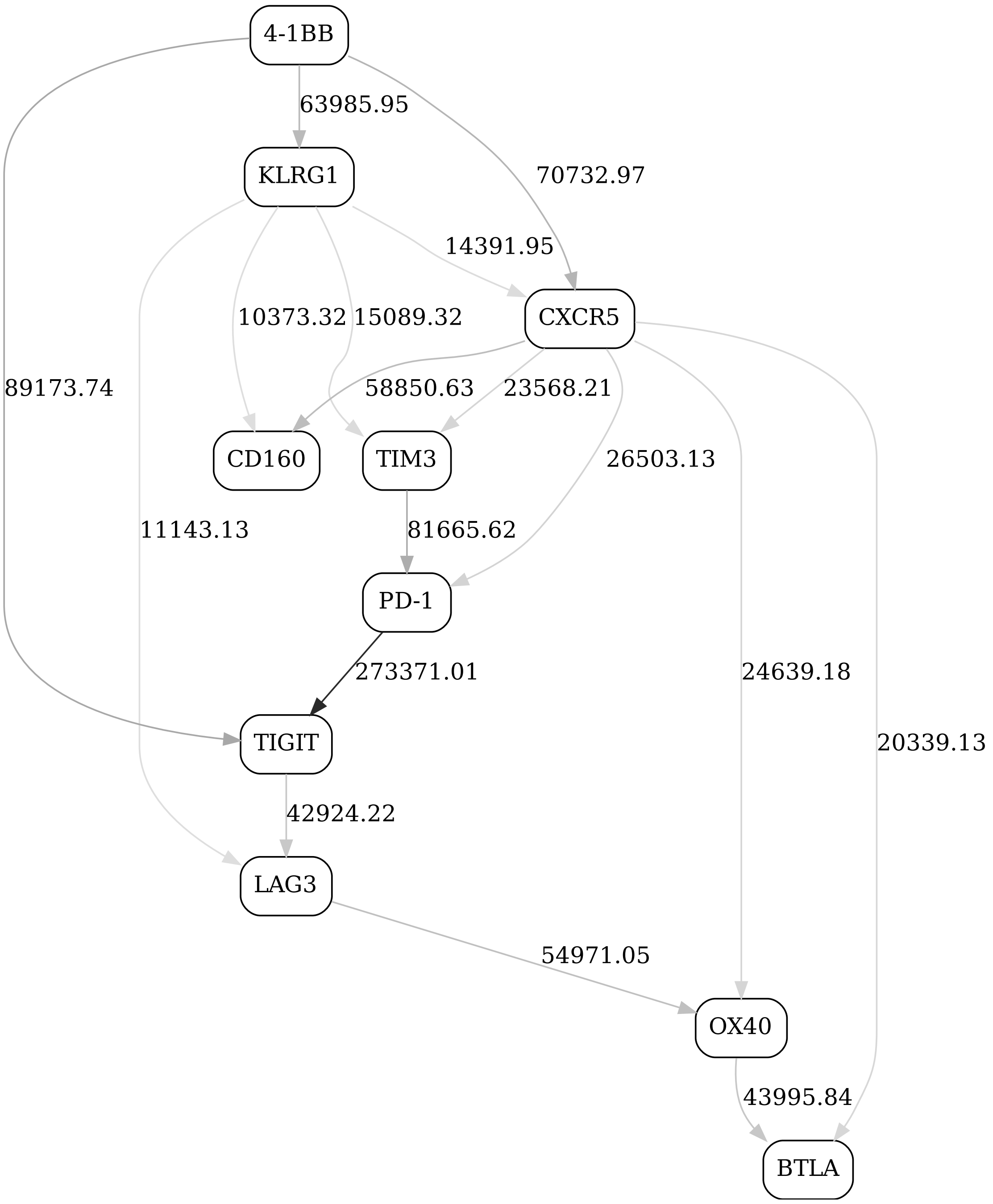


Figure 1: Checkpoint immune signaling network panel, naive CD4, day 1, responders

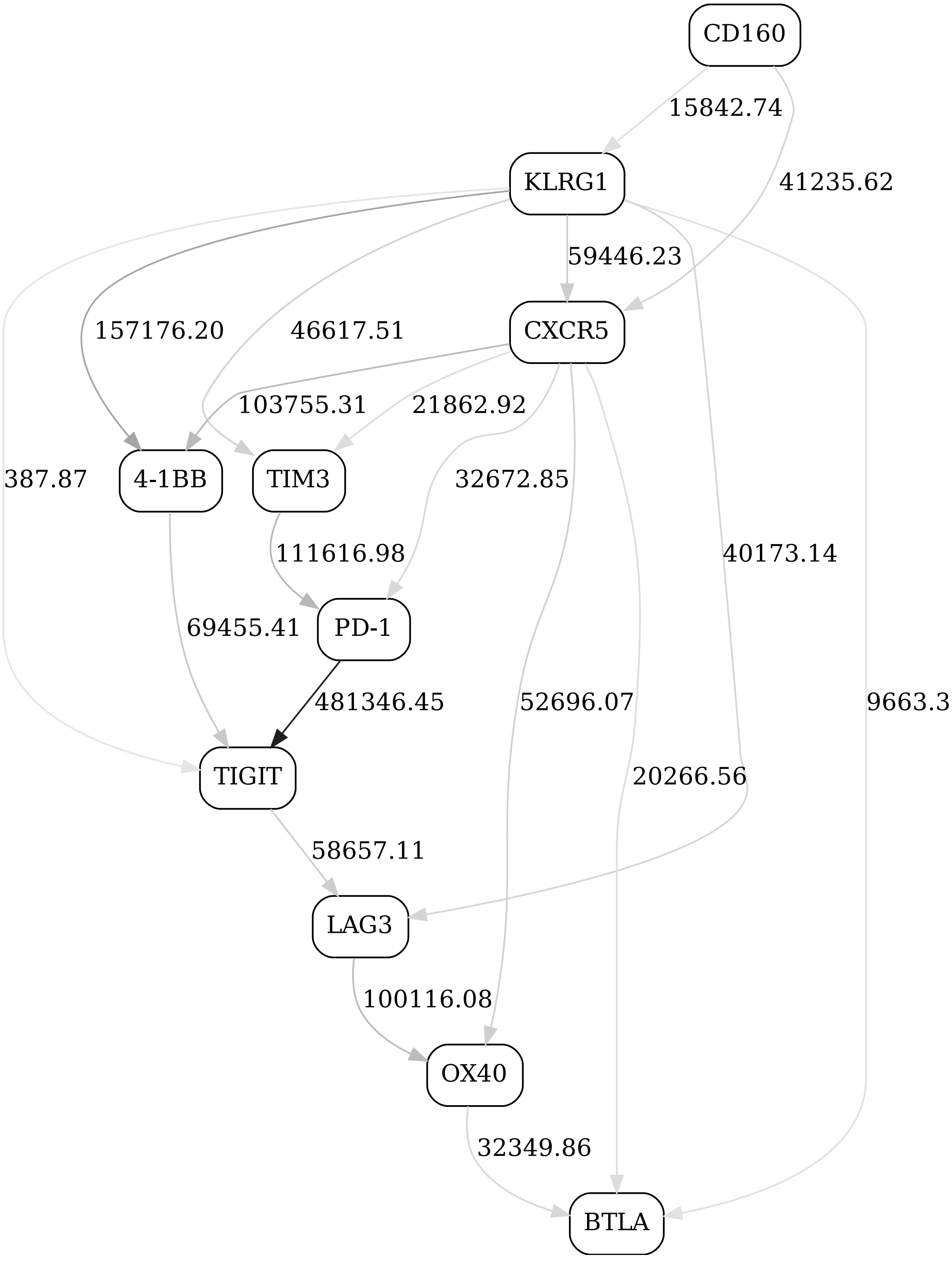


Figure 2: Checkpoint immune signaling network panel, naive CD4, day 1, non-responders

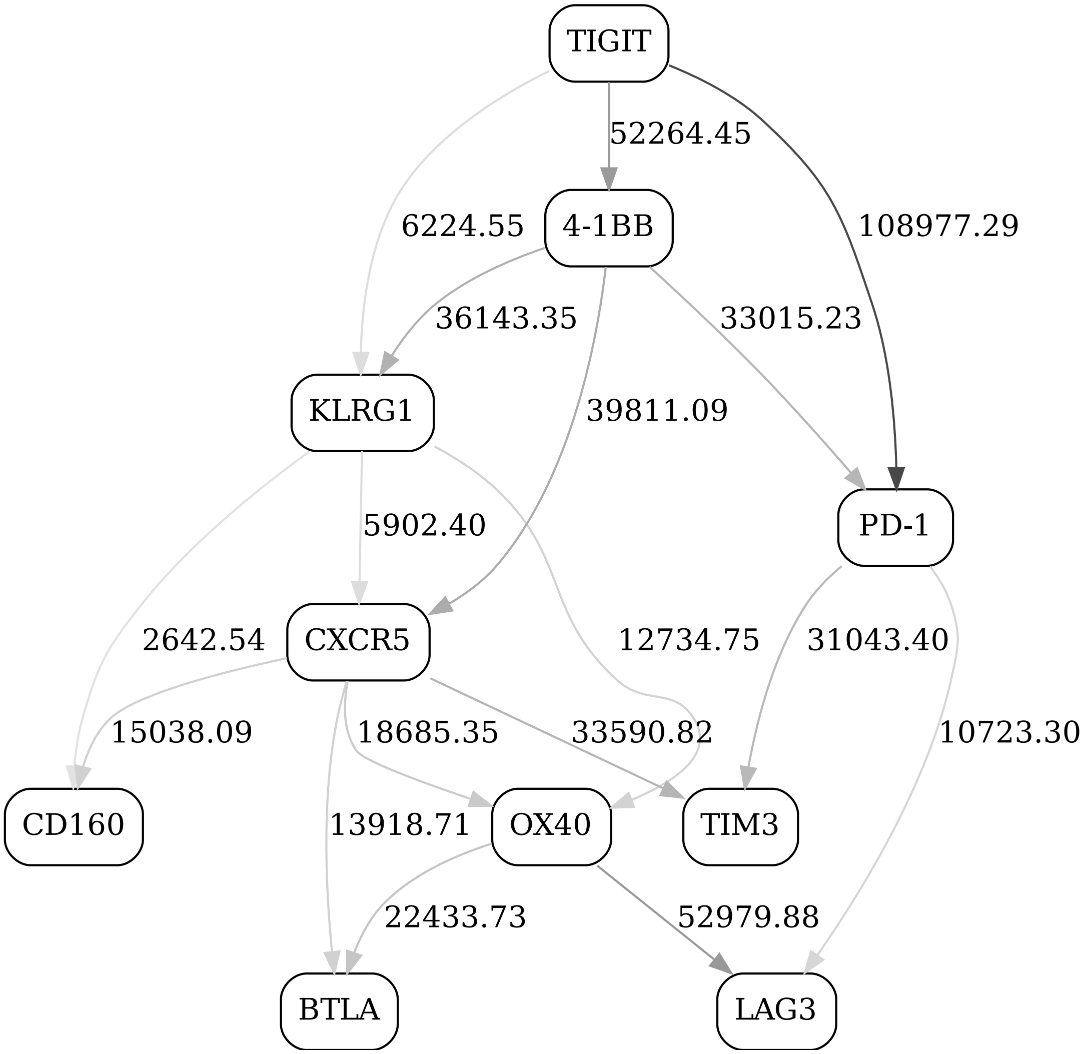


Figure 3: Checkpoint immune signaling network panel, naive CD4, day 21, responders

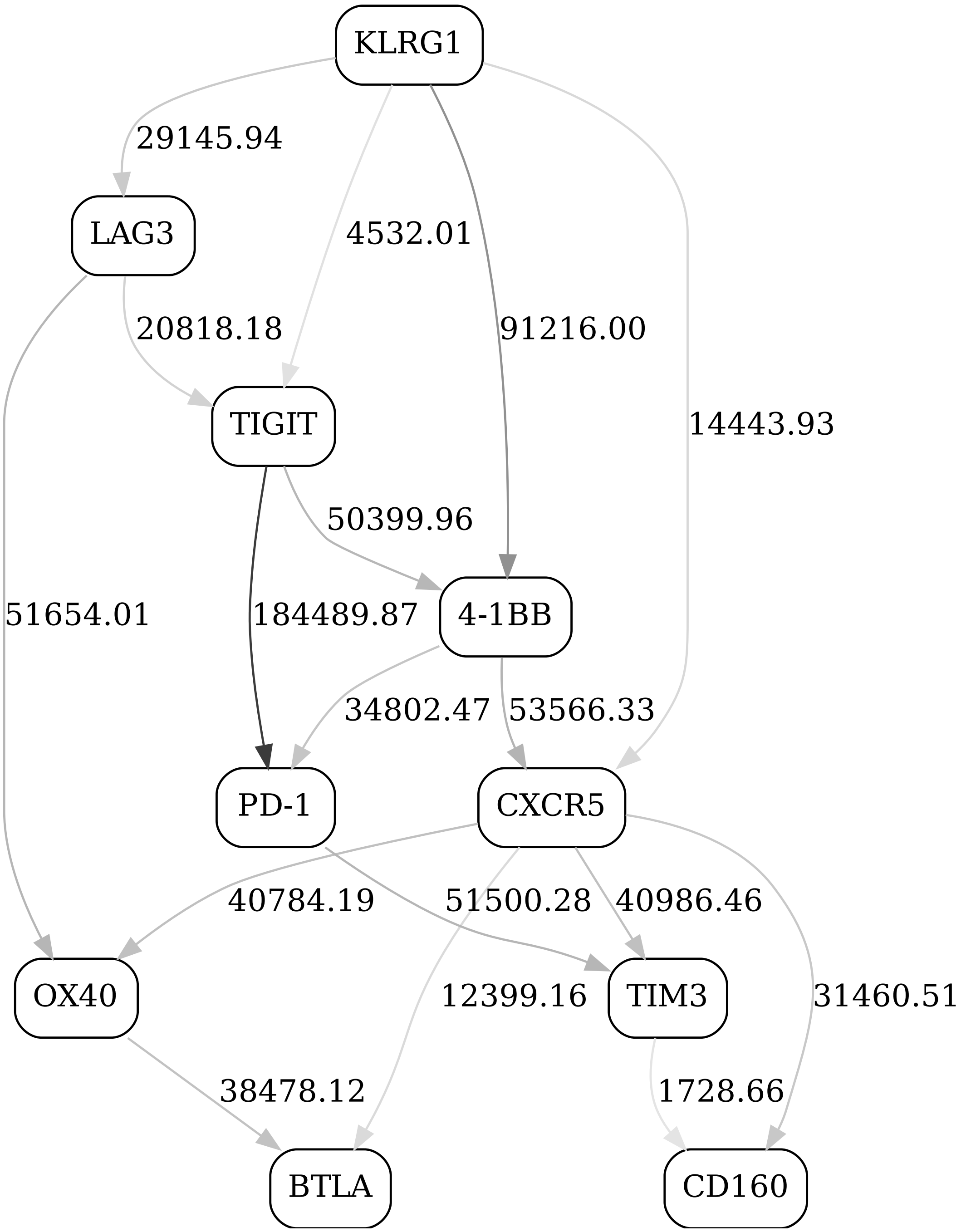


Figure 4: Checkpoint immune signaling network panel, naive CD4, day 21, non-responders

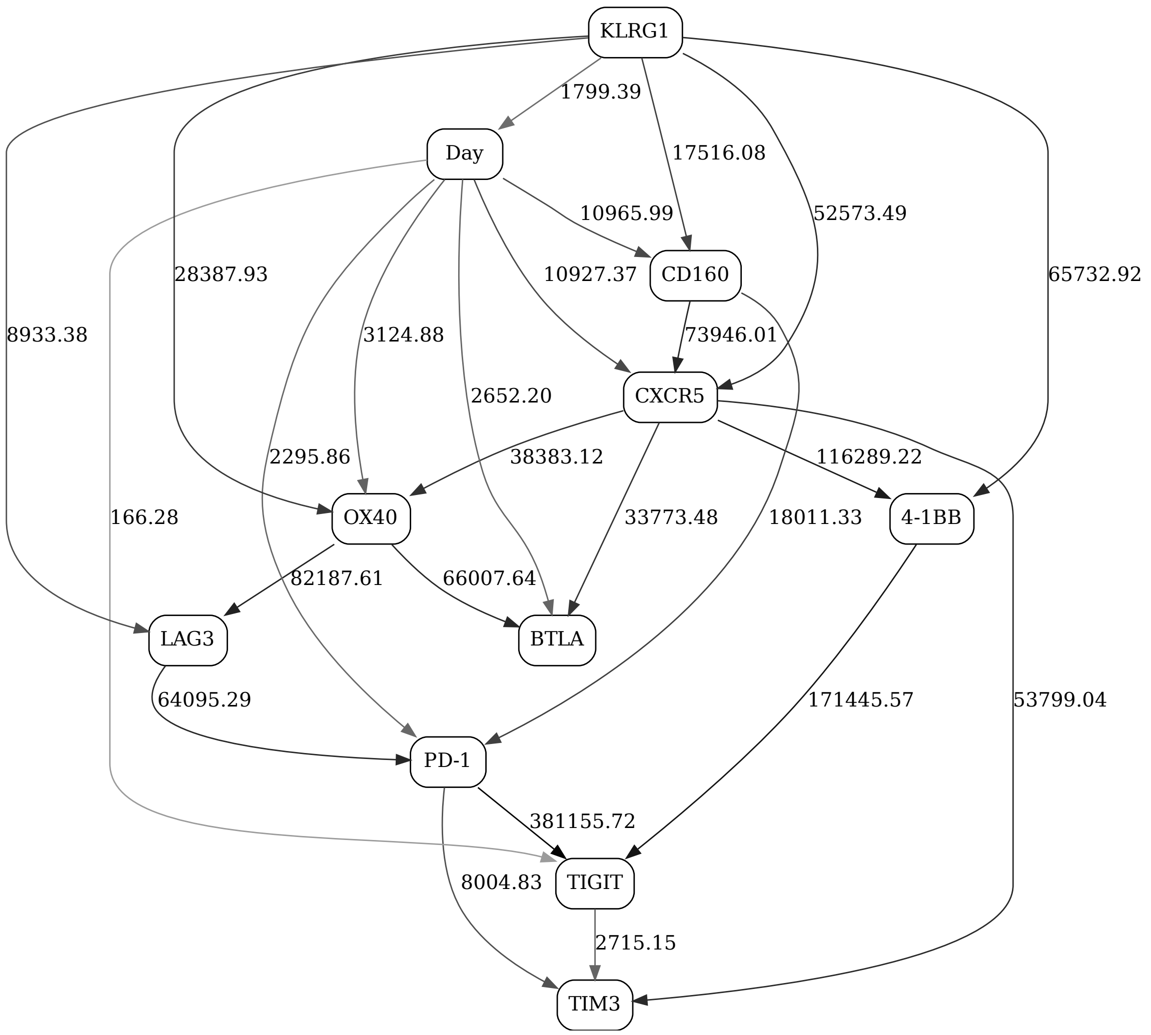


Figure 5: Checkpoint immune signaling network panel, naive CD4, day contrast (2-state "Day" variable), responders

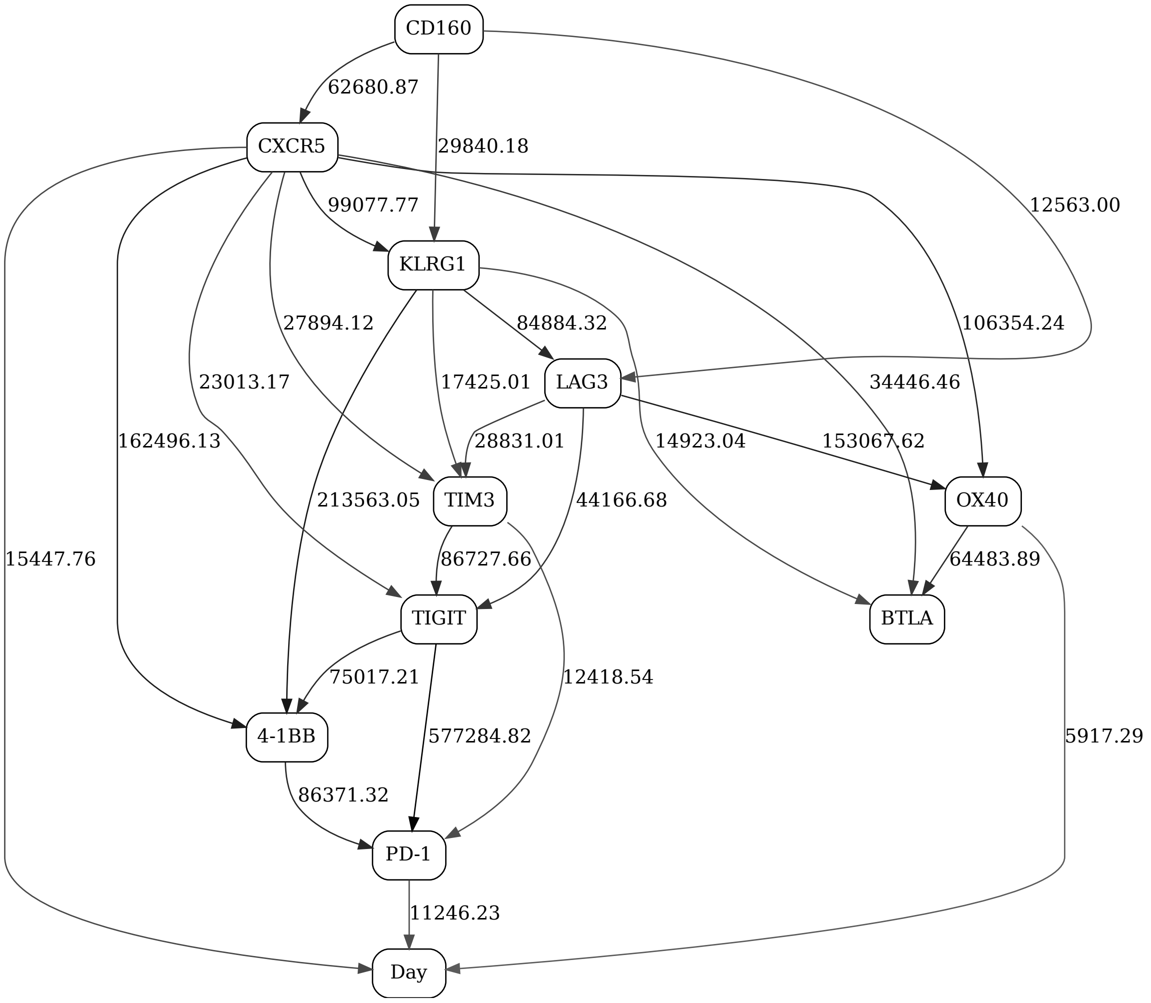


Figure 6: Checkpoint immune signaling network panel, naive CD4, day contrast (2-state "Day" variable), non-responders

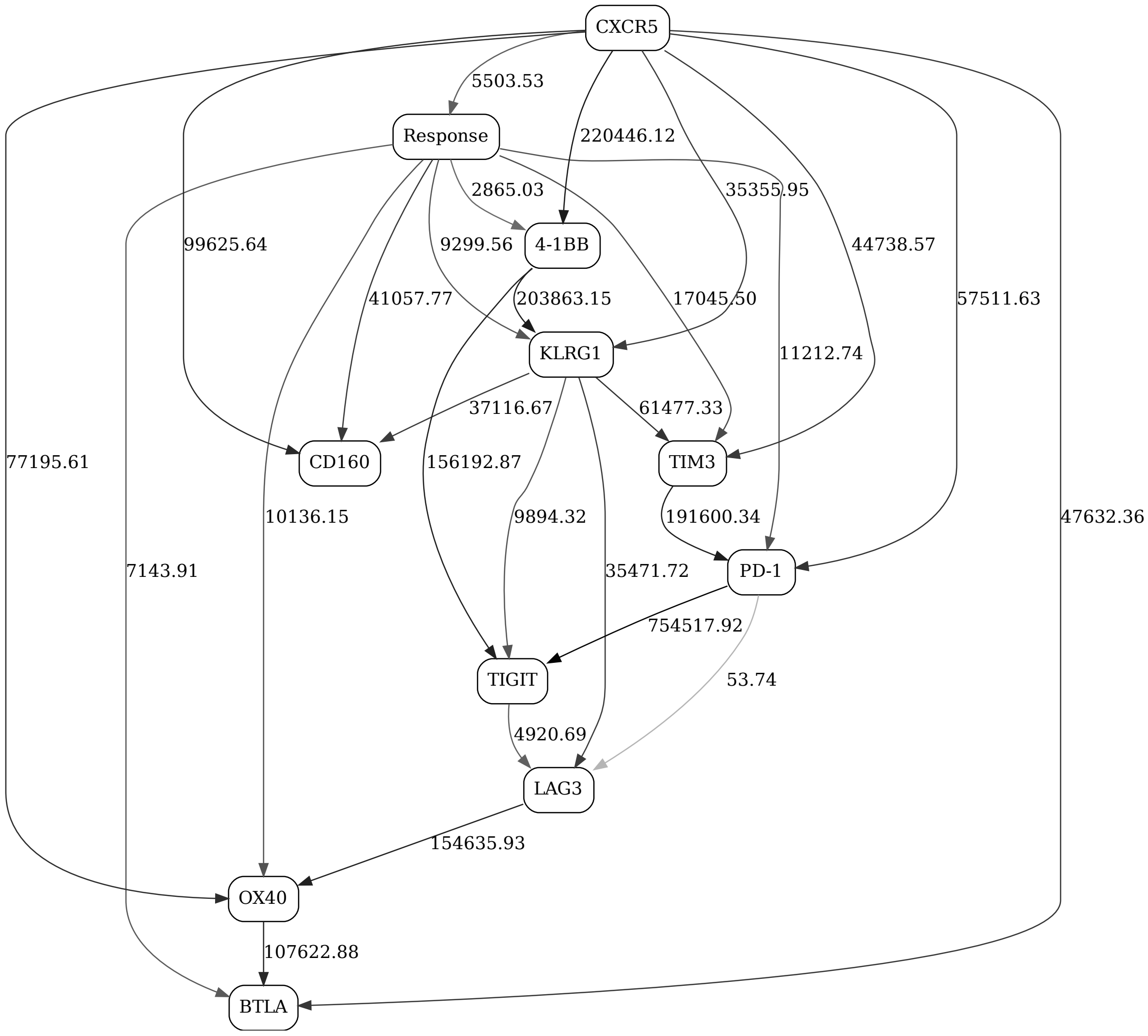


Figure 7: Checkpoint immune signaling network panel, naive CD4, day 1, response contrast (2-state "Response" variable)

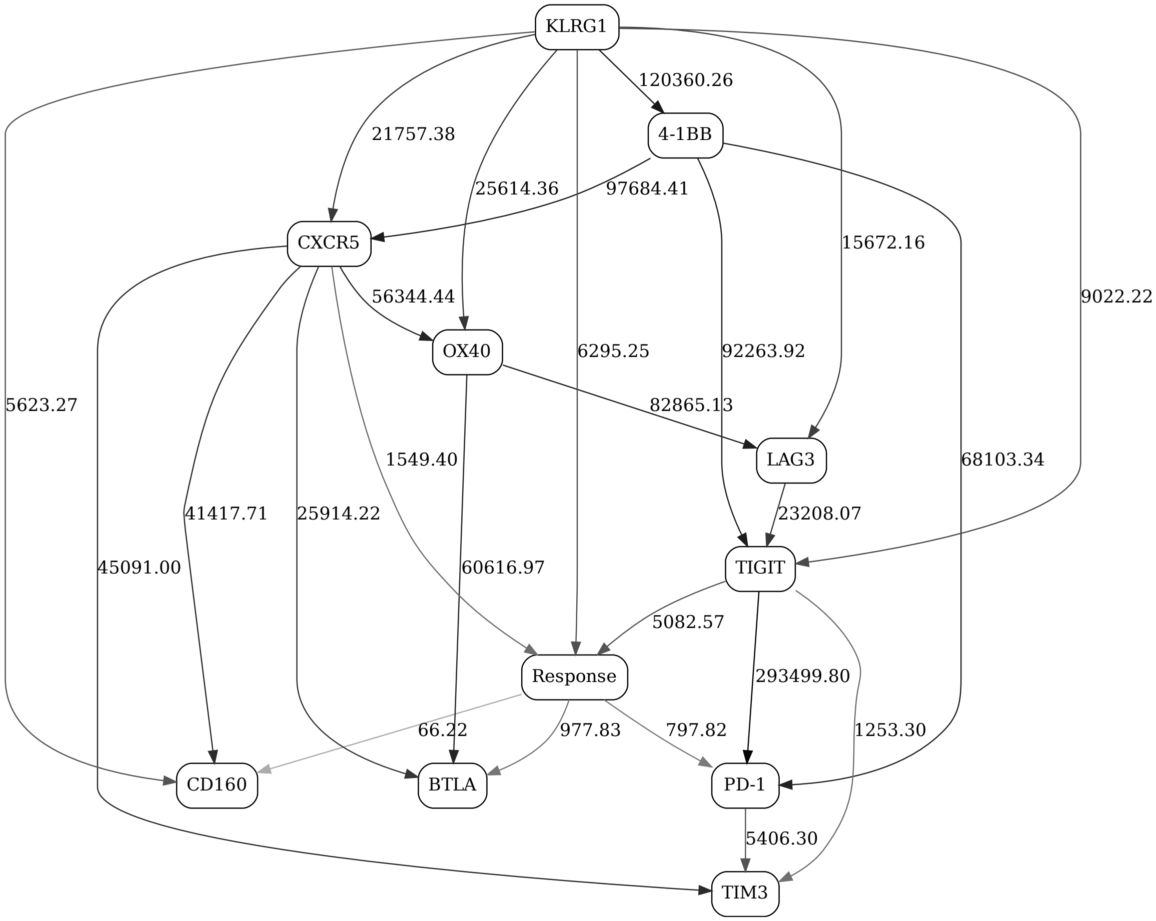


Figure 8: Checkpoint immune signaling network panel, naive CD4, day 21, response contrast (2-state "Response" variable)

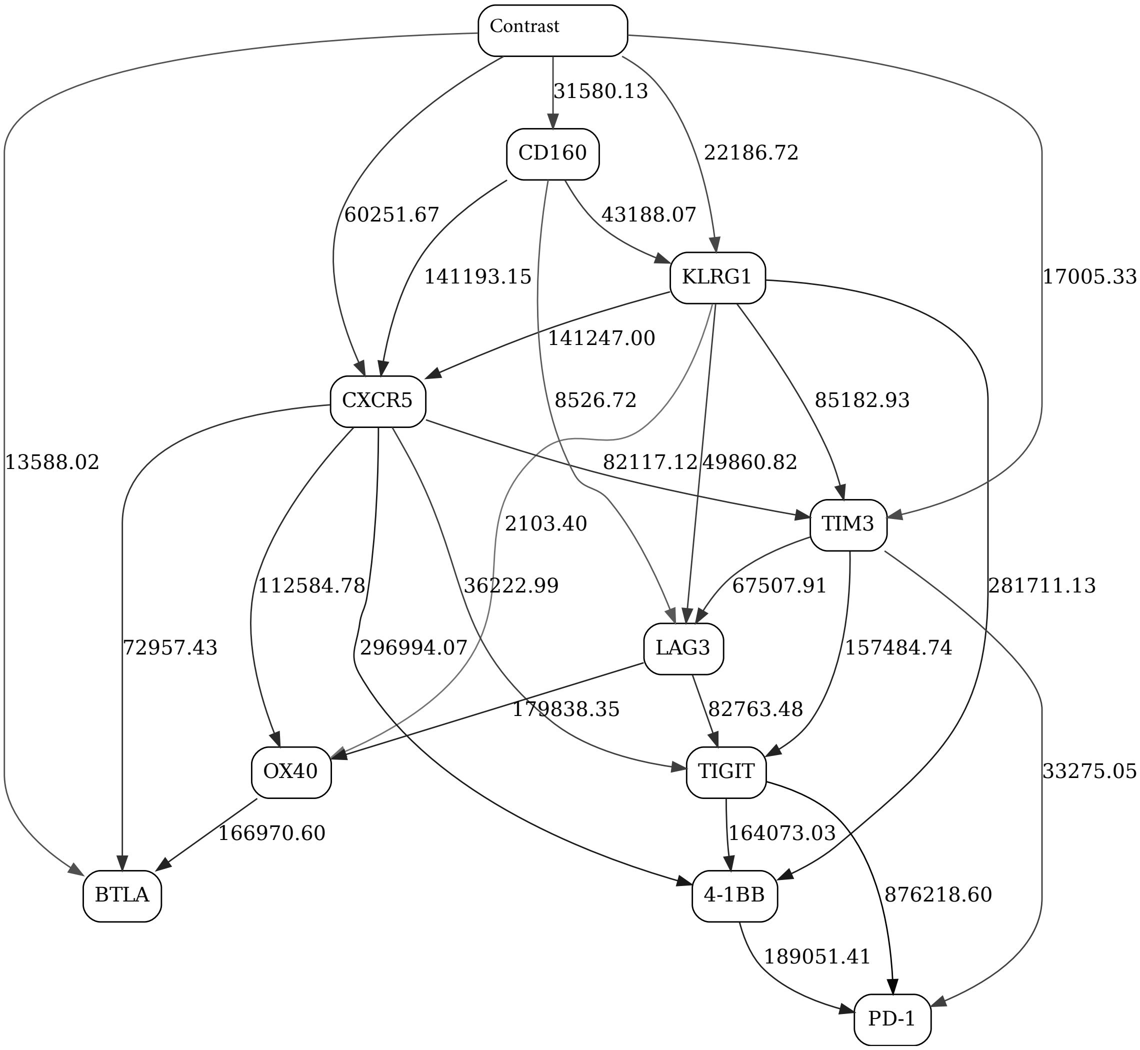


Figure 9: Checkpoint immune signaling network panel, naive CD4

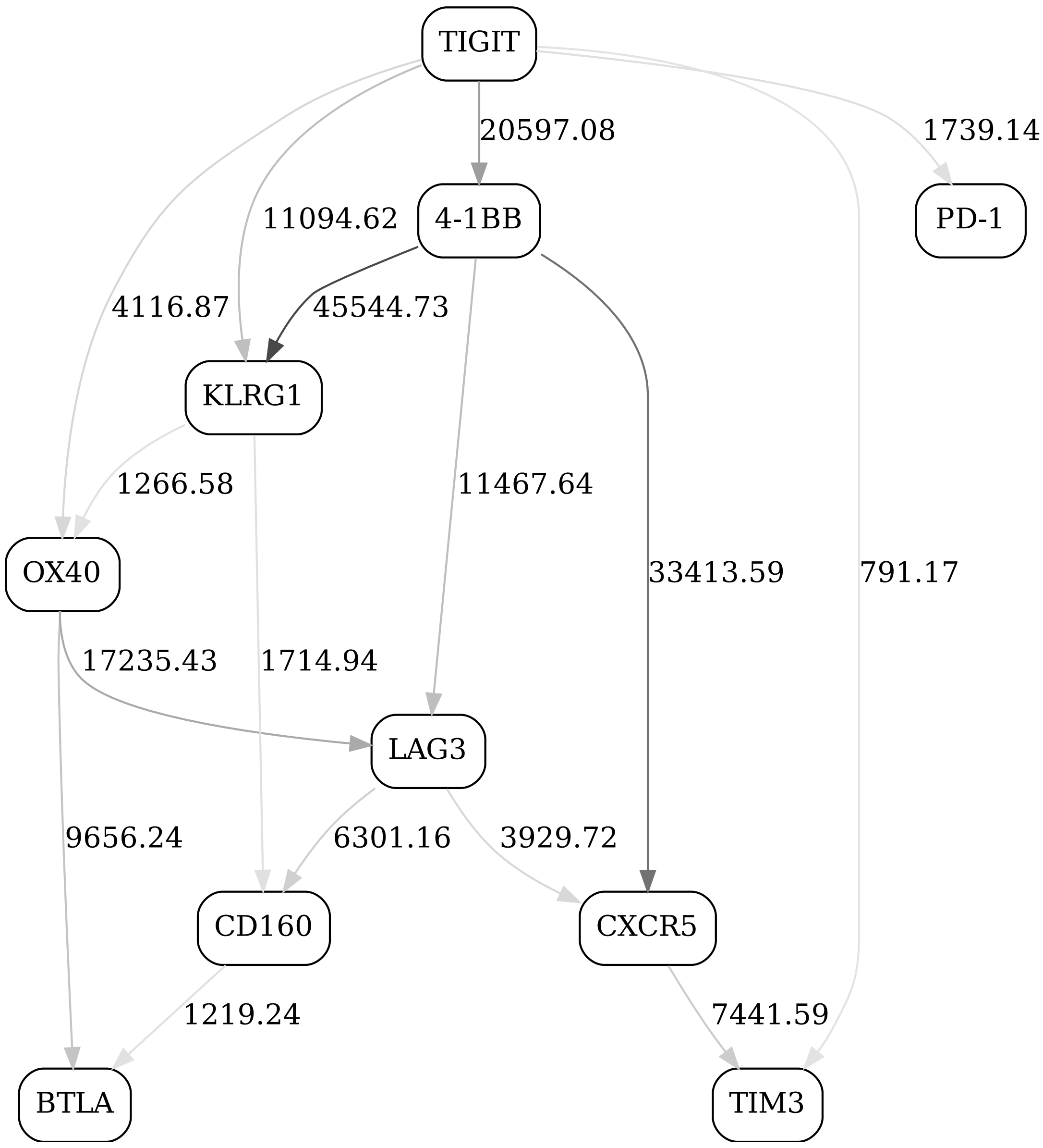


Figure 10: Checkpoint immune signaling network panel, non-naive CD8, day 1, responders

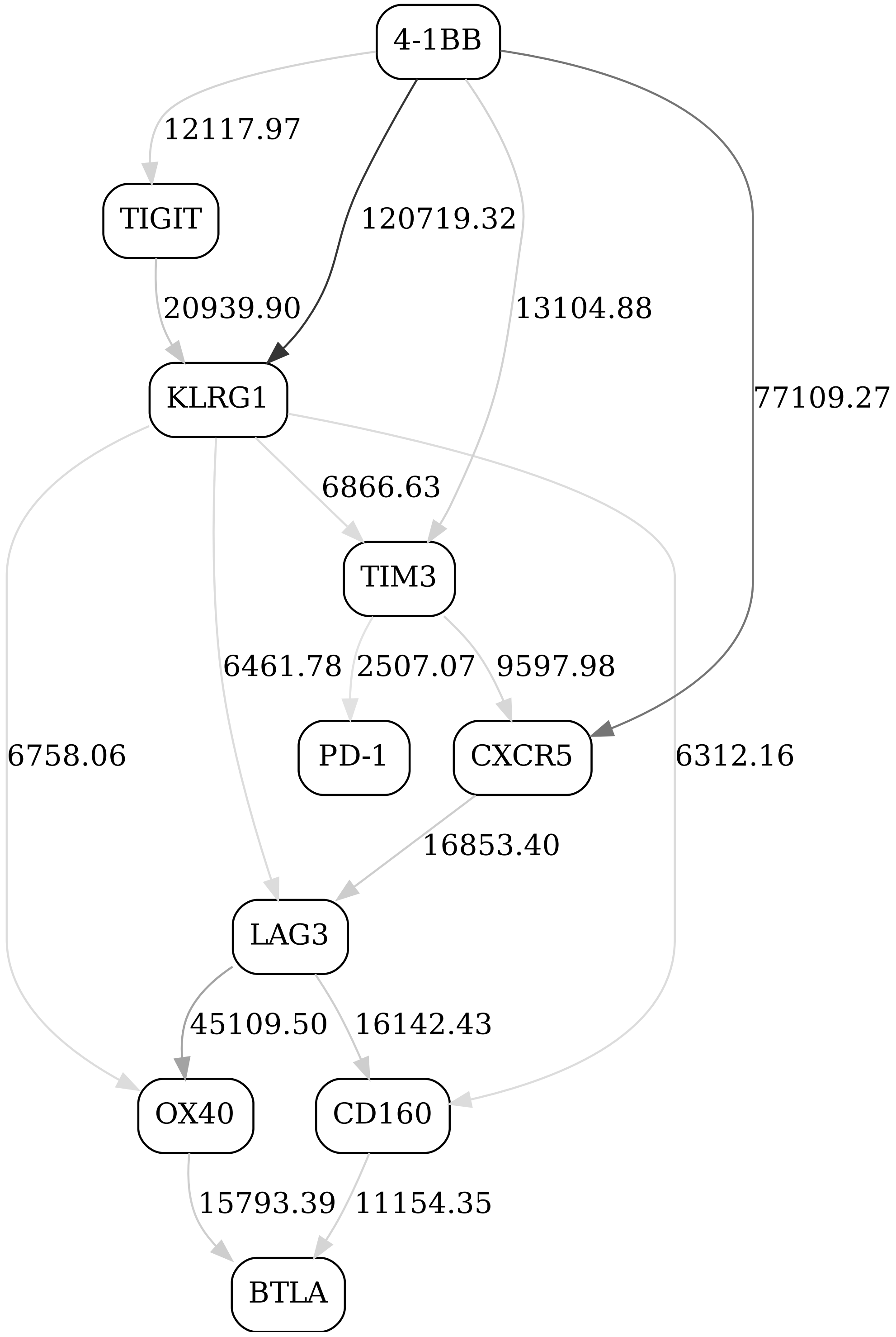


Figure 11: Checkpoint immune signaling network panel, non-naive CD8, day 1, non-responders

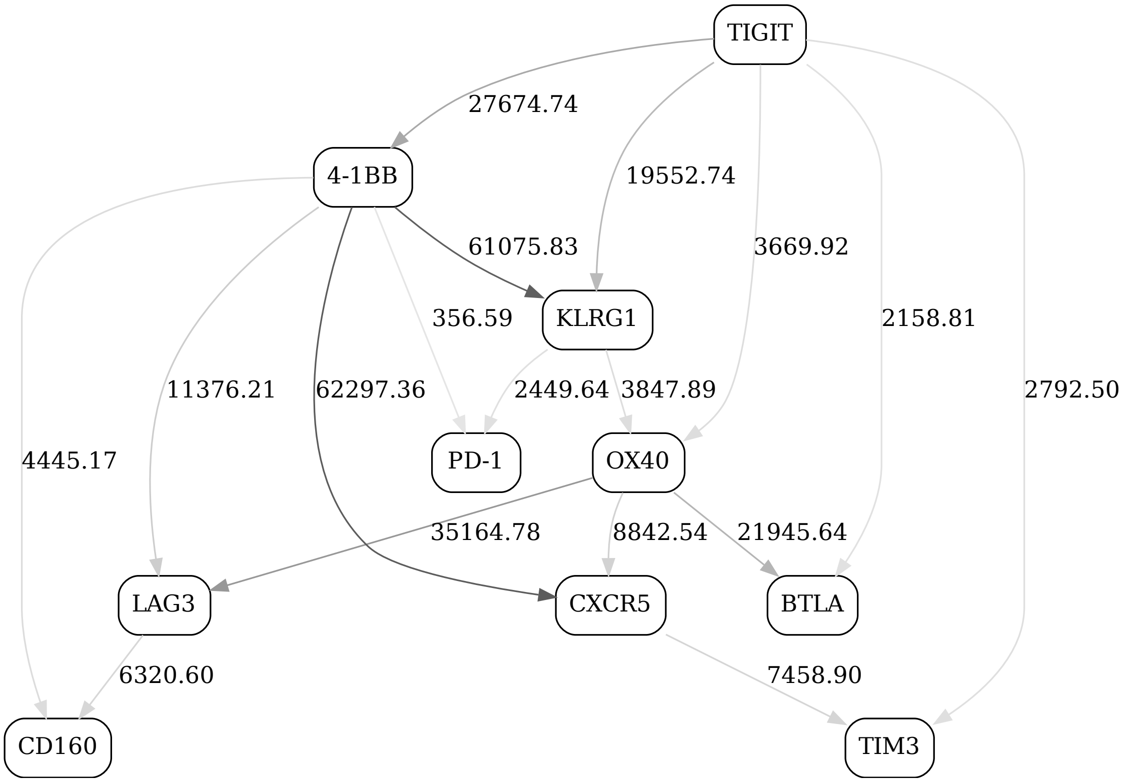


Figure 12: Checkpoint immune signaling network panel, non-naive CD8, day 21, responders

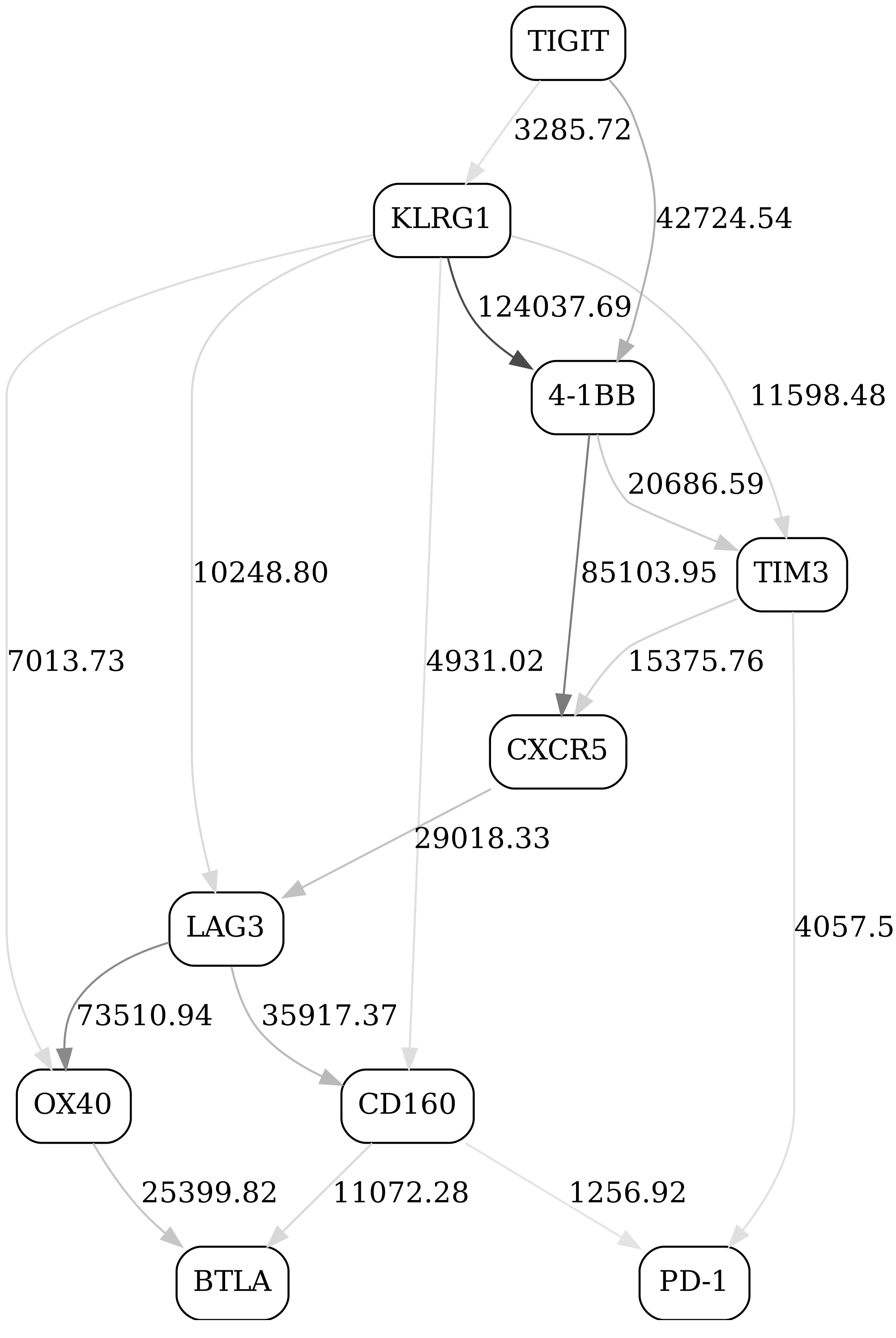


Figure 13: Checkpoint immune signaling network panel, non-naive CD8, day 21, non-responders

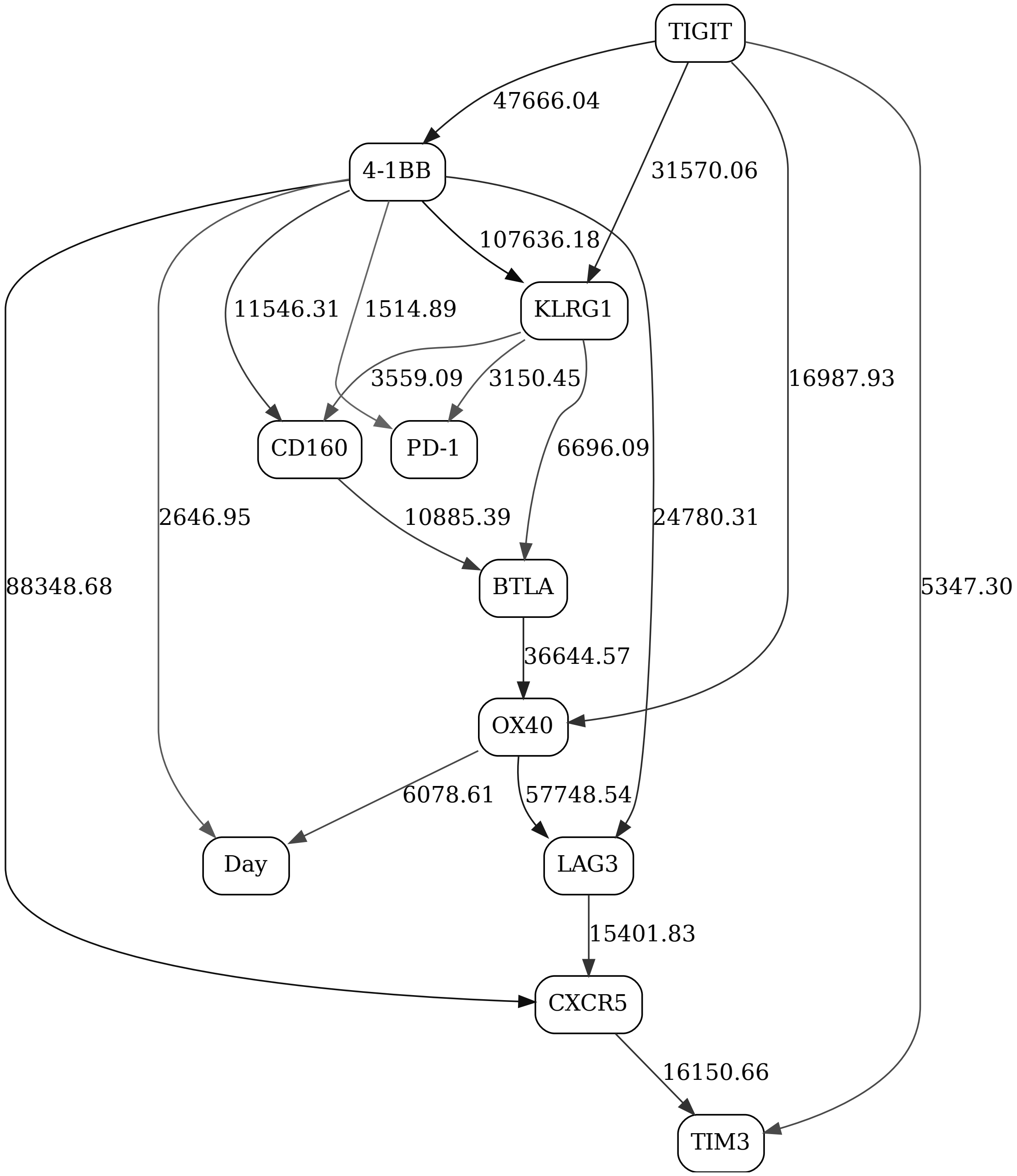


Figure 14: Checkpoint immune signaling network panel, non-naive CD8, day contrast (2-state "Day" variable), responders

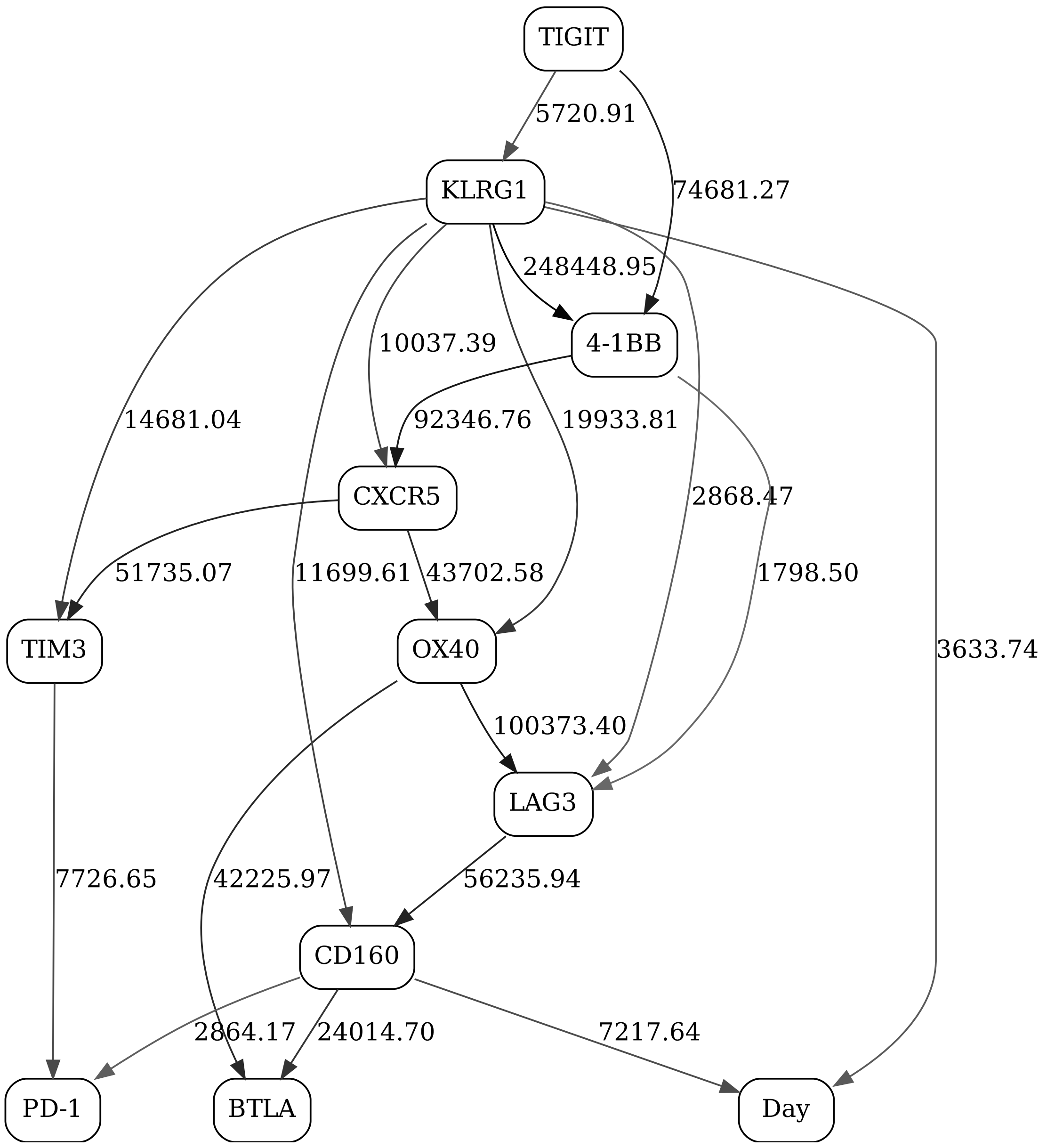


Figure 15: Checkpoint immune signaling network panel, non-naive CD8, day contrast (2-state "Day" vari-able), non-responders

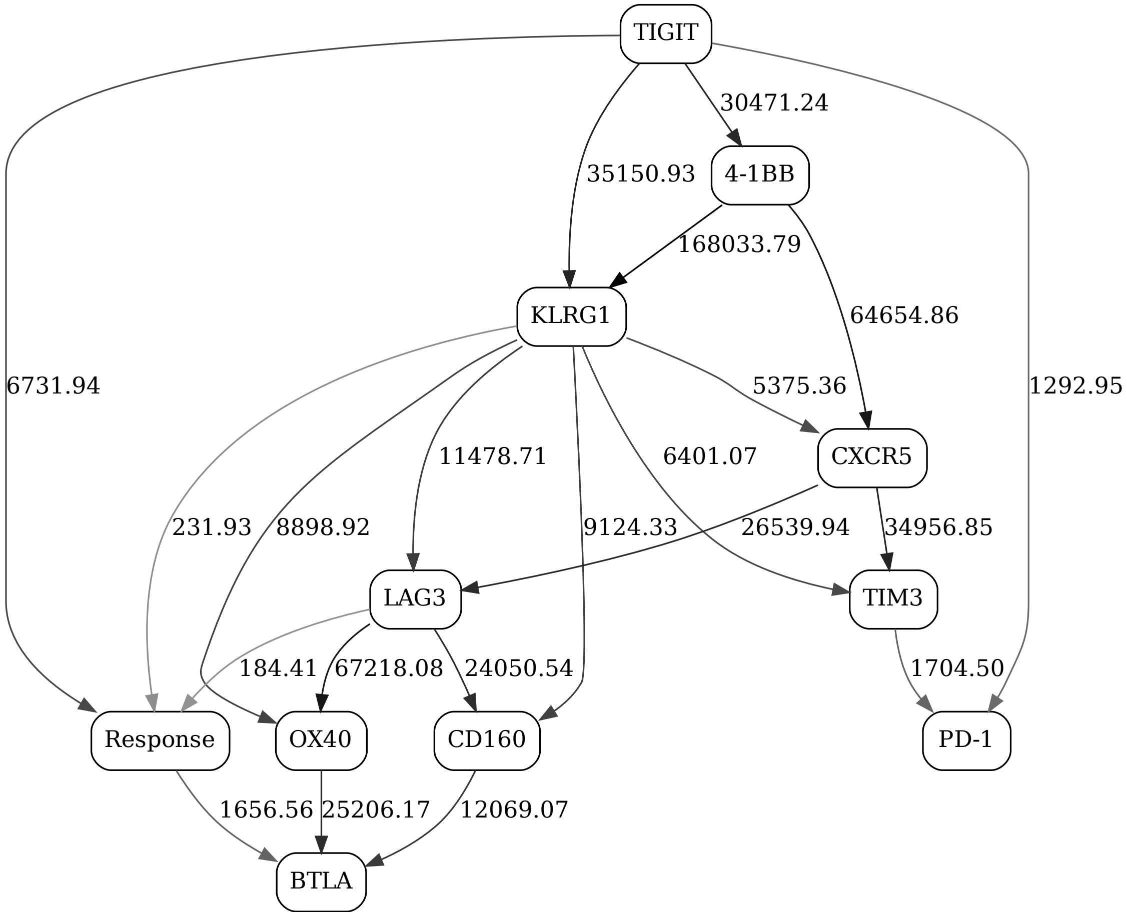


Figure 16: Checkpoint immune signaling network panel, non-naive CD8, day 1, response contrast (2-state "Response" variable)

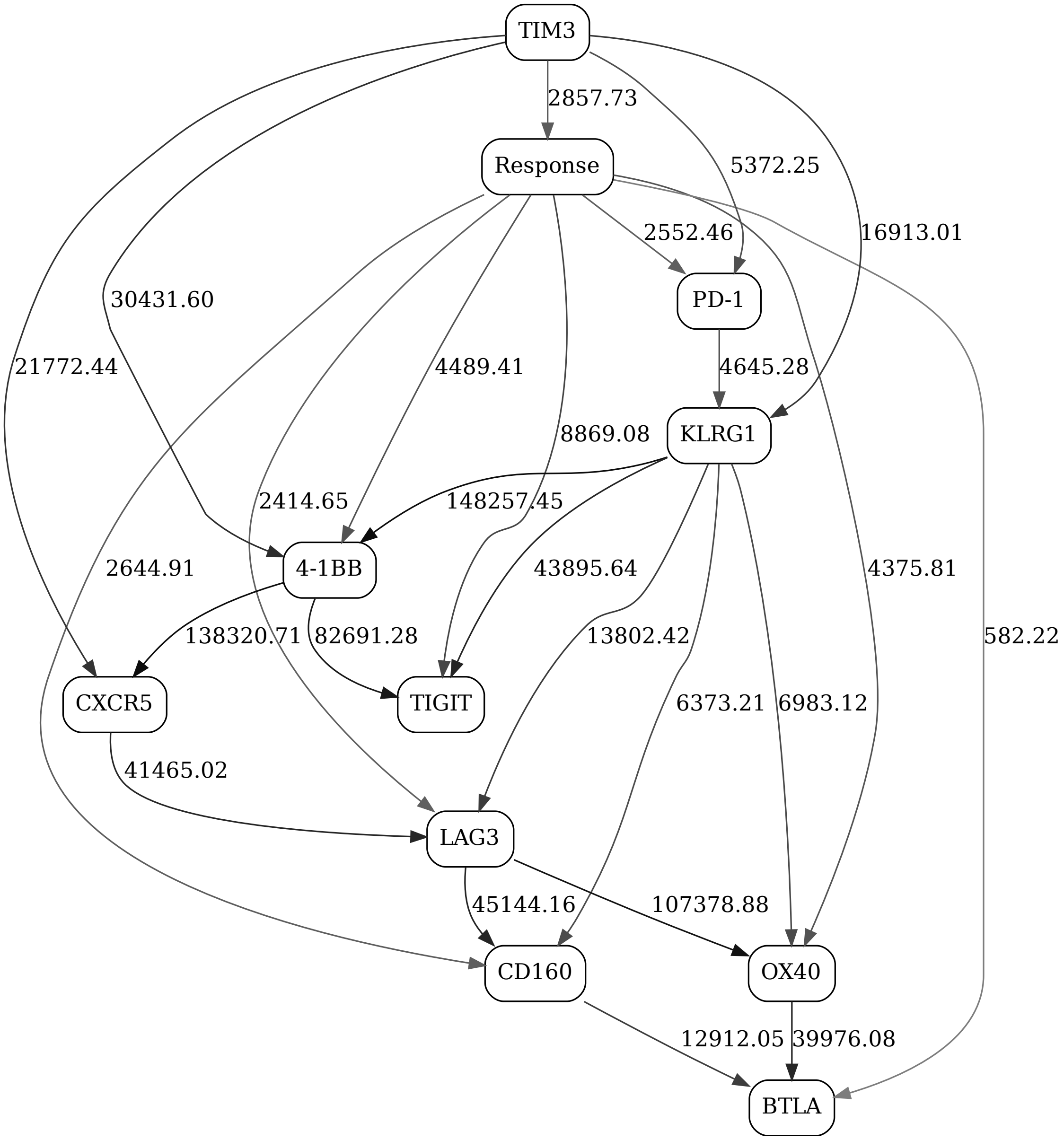


Figure 17: Checkpoint immune signaling network panel, non-naive CD8, day 21, response contrast (2-state "Response" variable)

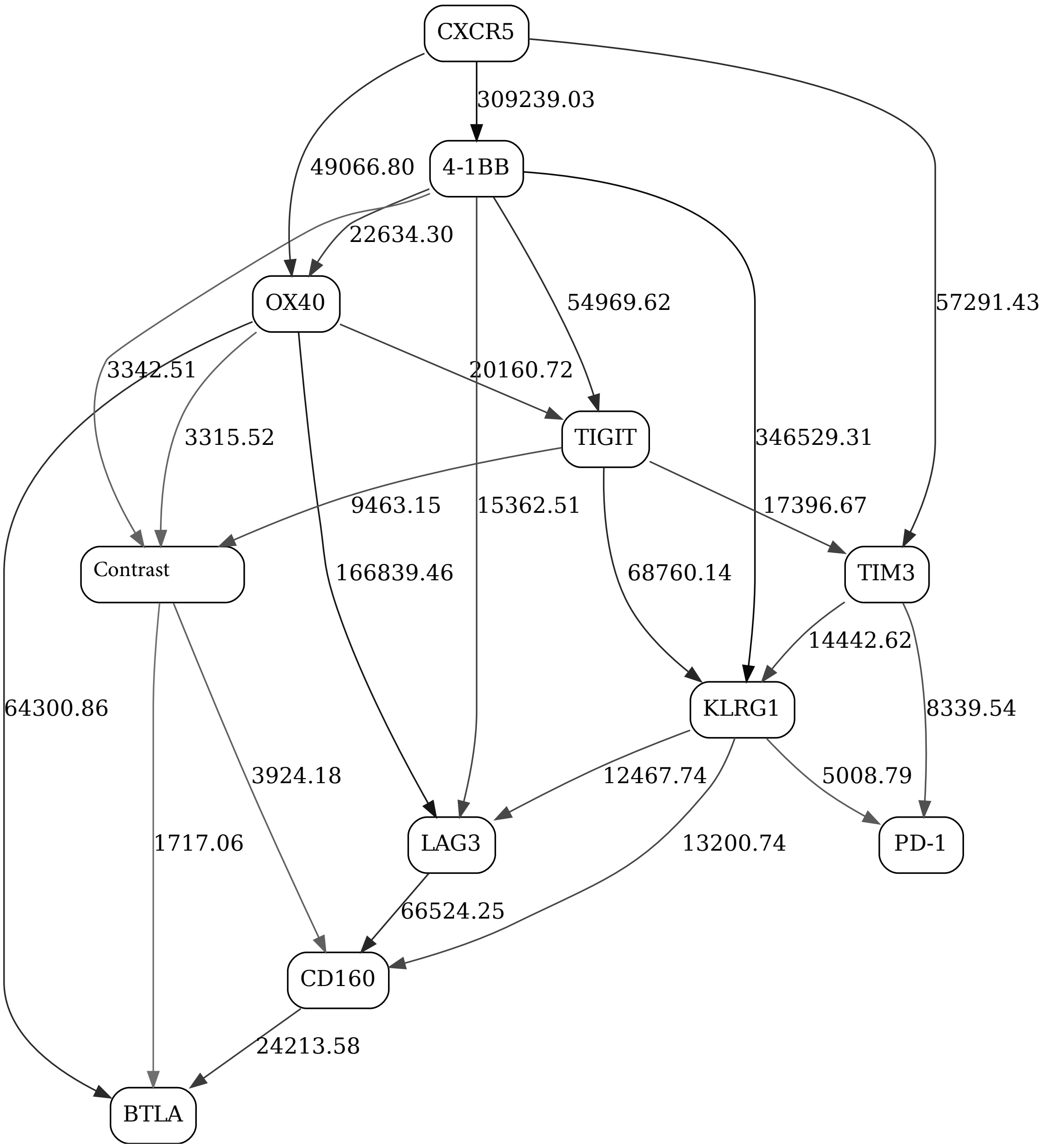


Figure 18: Checkpoint immune signaling network panel, non-naive CD8

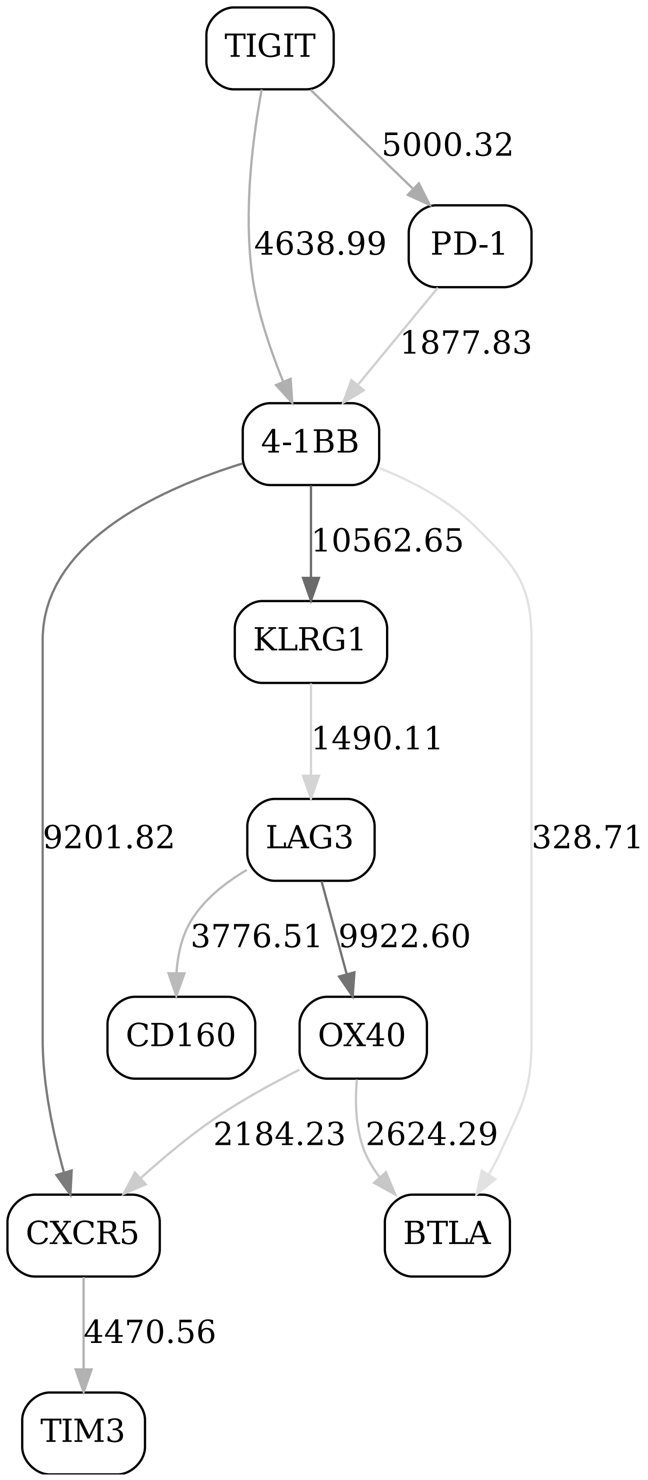


Figure 19: Checkpoint immune signaling network panel, naive CD8, day 1, responders

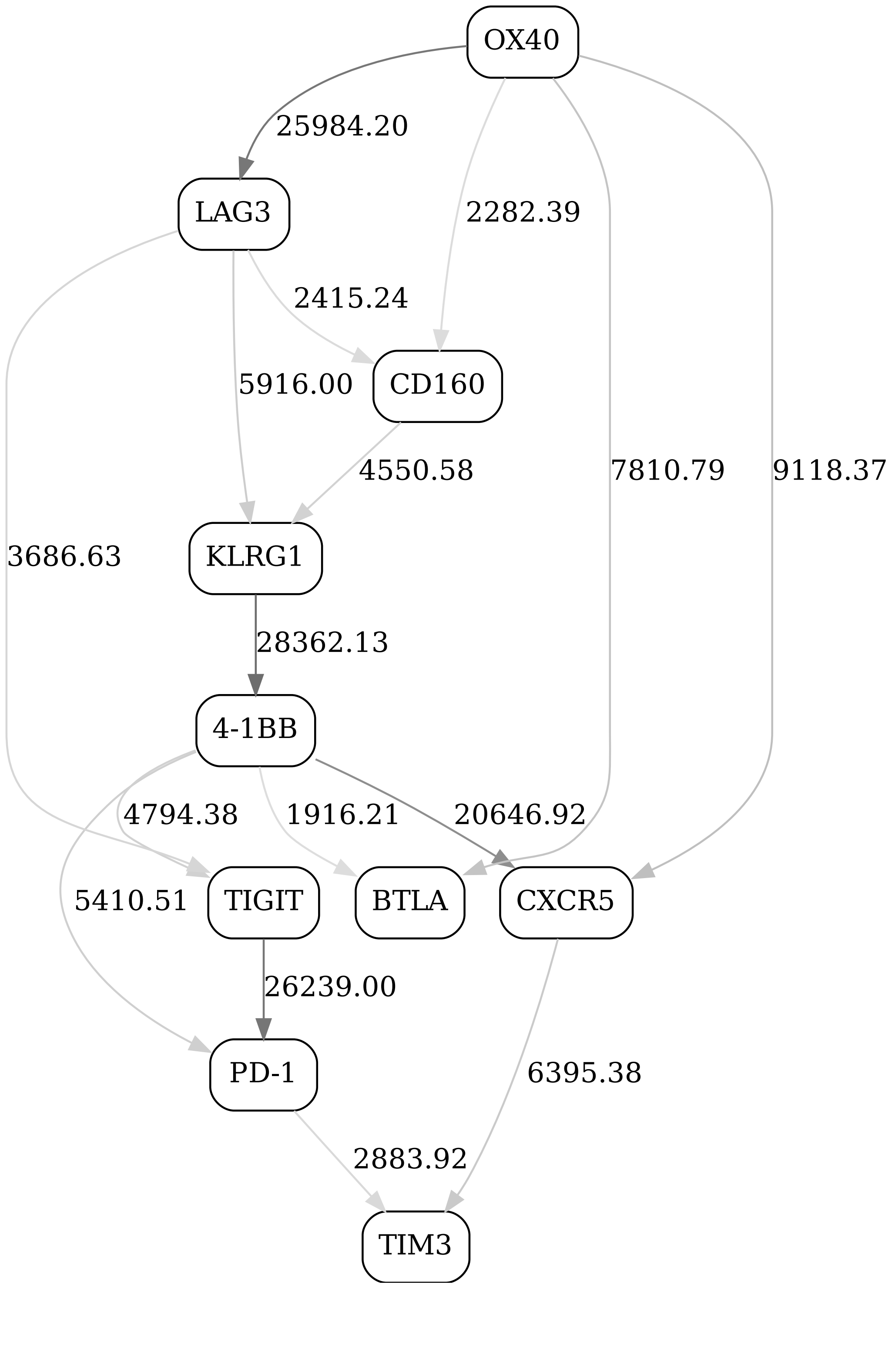


Figure 20: Checkpoint immune signaling network panel, naive CD8, day 1, non-responders

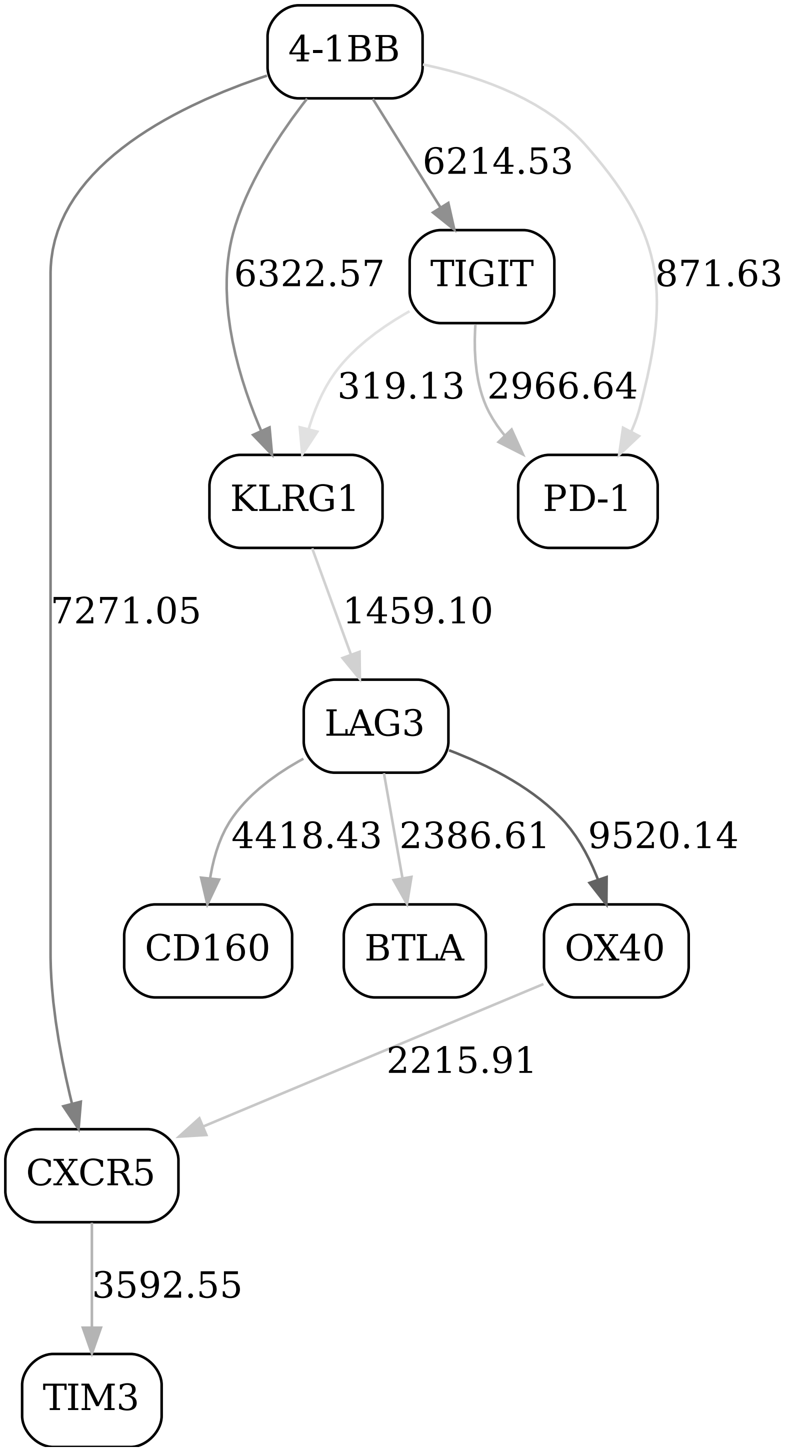


Figure 21: Checkpoint immune signaling network panel, naive CD8, day 21, responders

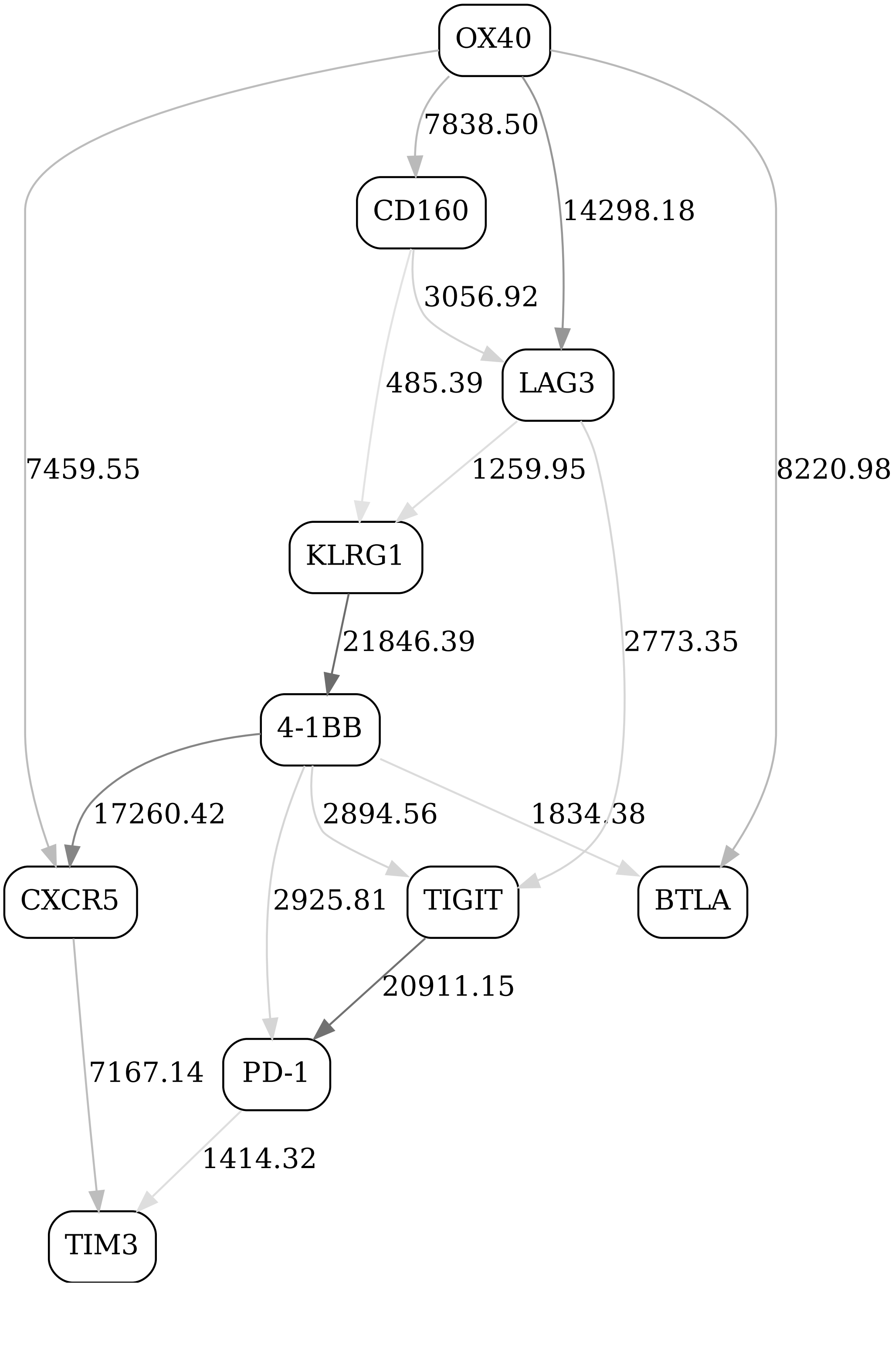


Figure 22: Checkpoint immune signaling network panel, naive CD8, day 21, non-responders

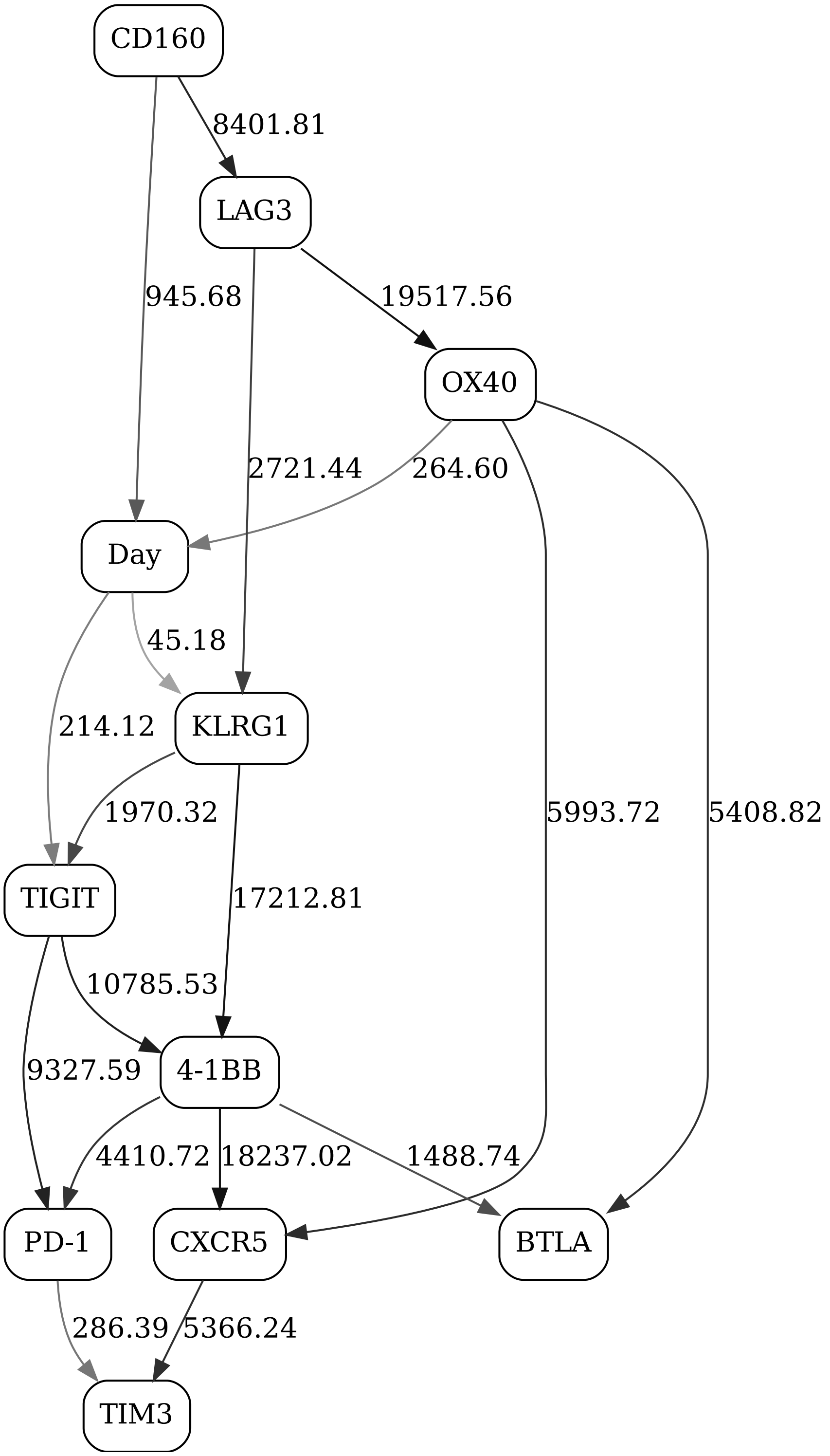


Figure 23: Checkpoint immune signaling network panel, naive CD8, day contrast (2-state "Day" variable), responders

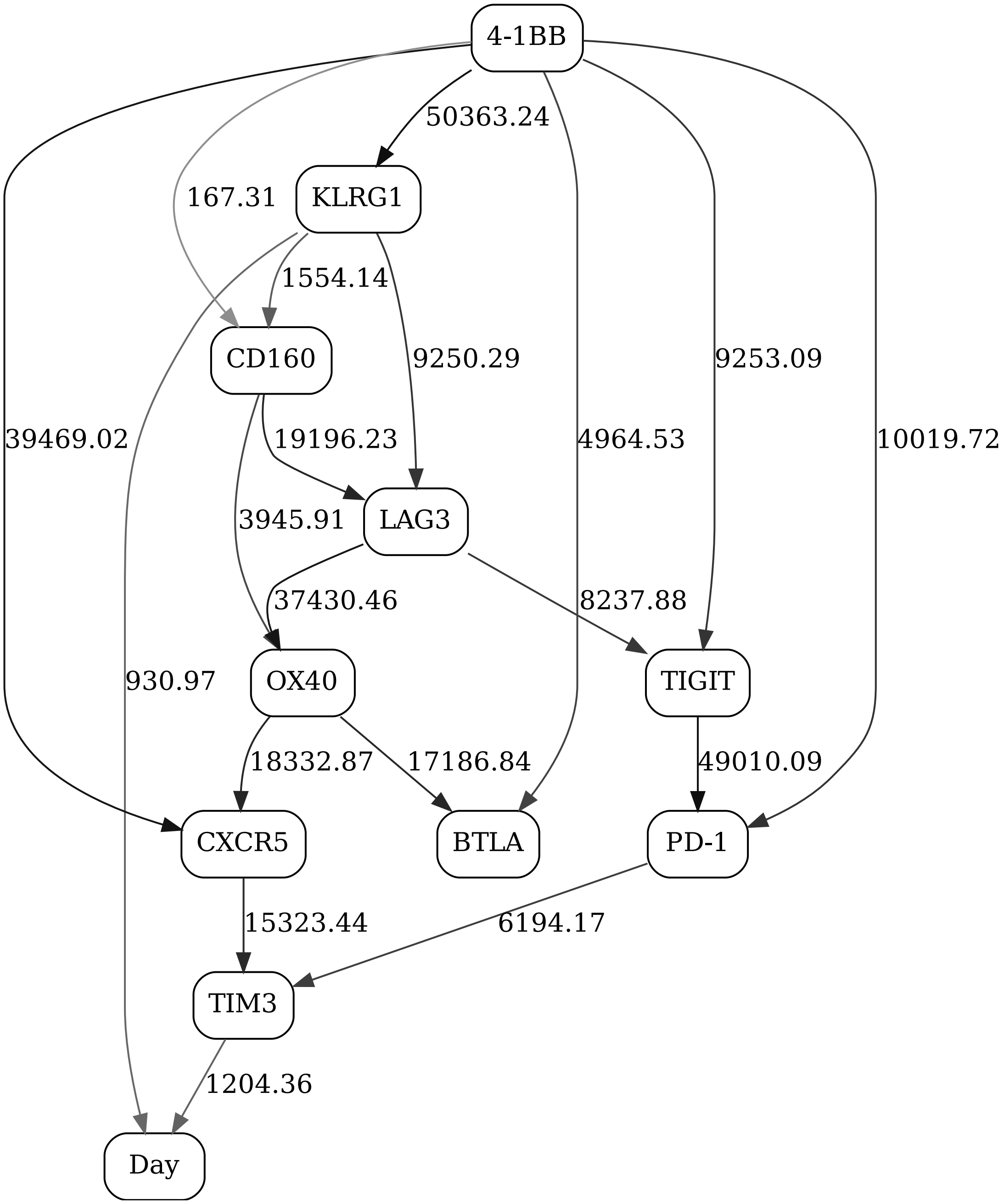


Figure 24: Checkpoint immune signaling network panel, naive CD8, day contrast (2-state "Day" variable), non-responders

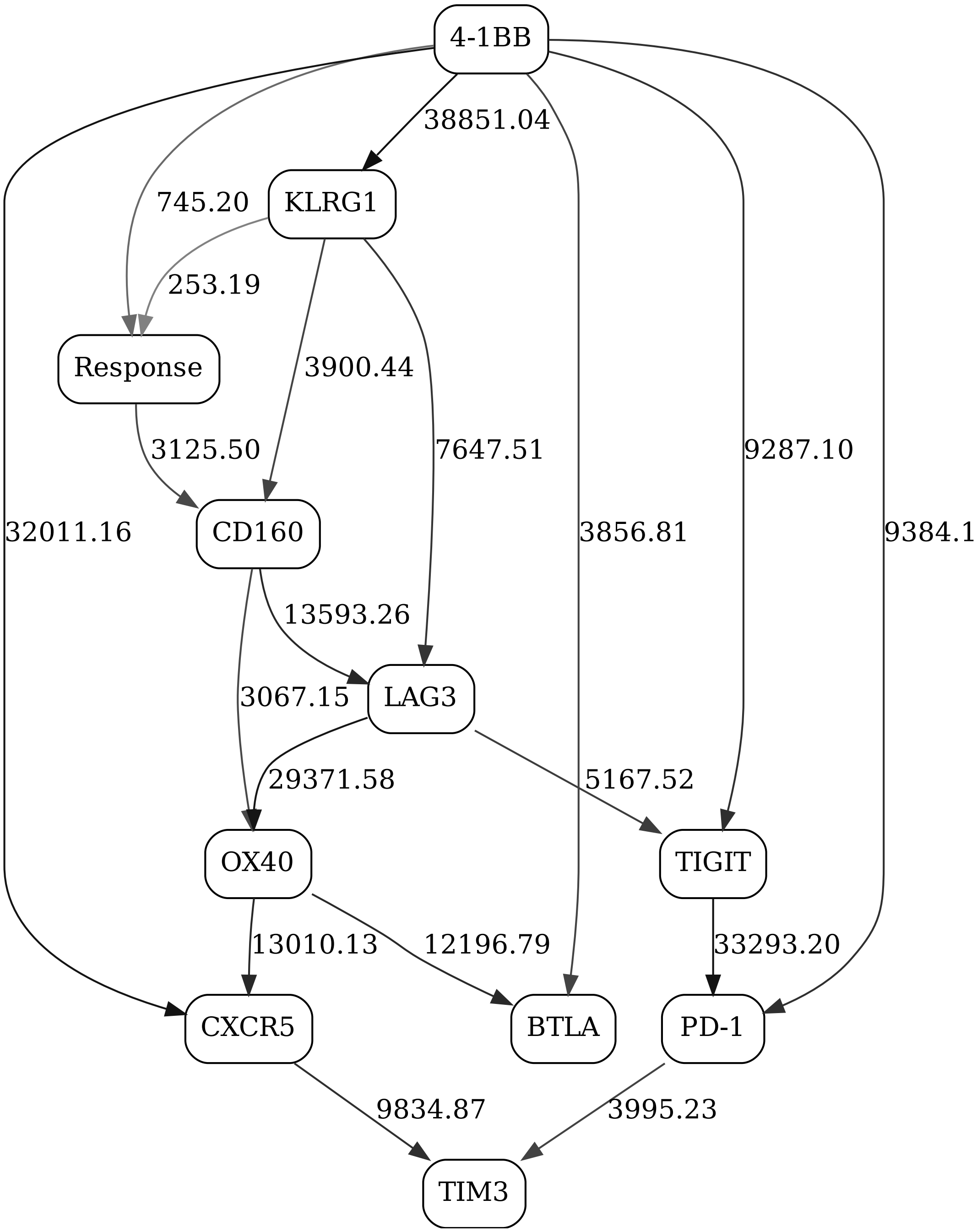


Figure 25: Checkpoint immune signaling network panel, naive CD8, day 1, response contrast (2-state "Response" variable)

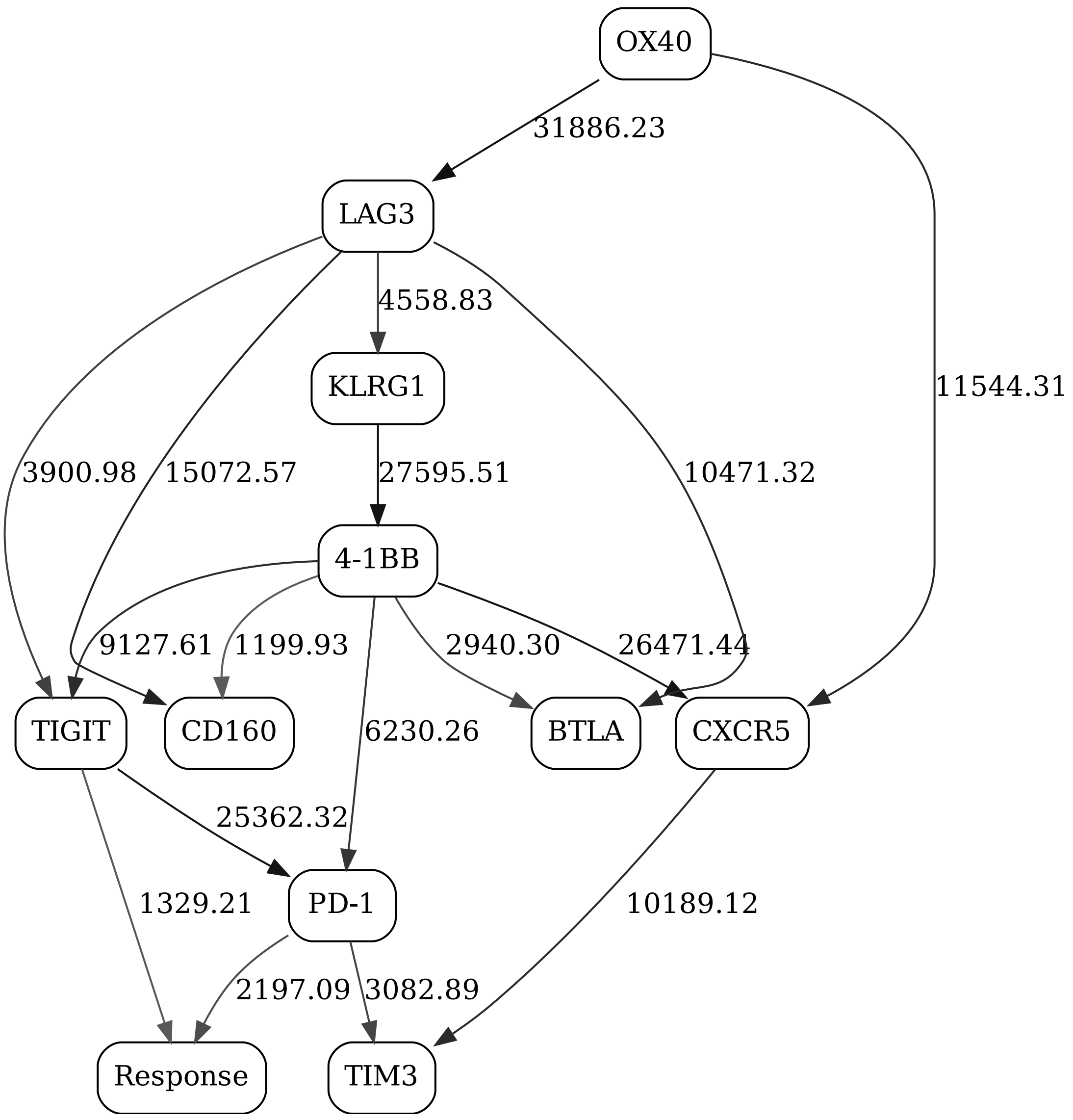


Figure 26: Checkpoint immune signaling network panel, naive CD8, day 21, response contrast (2-state "Response" variable)

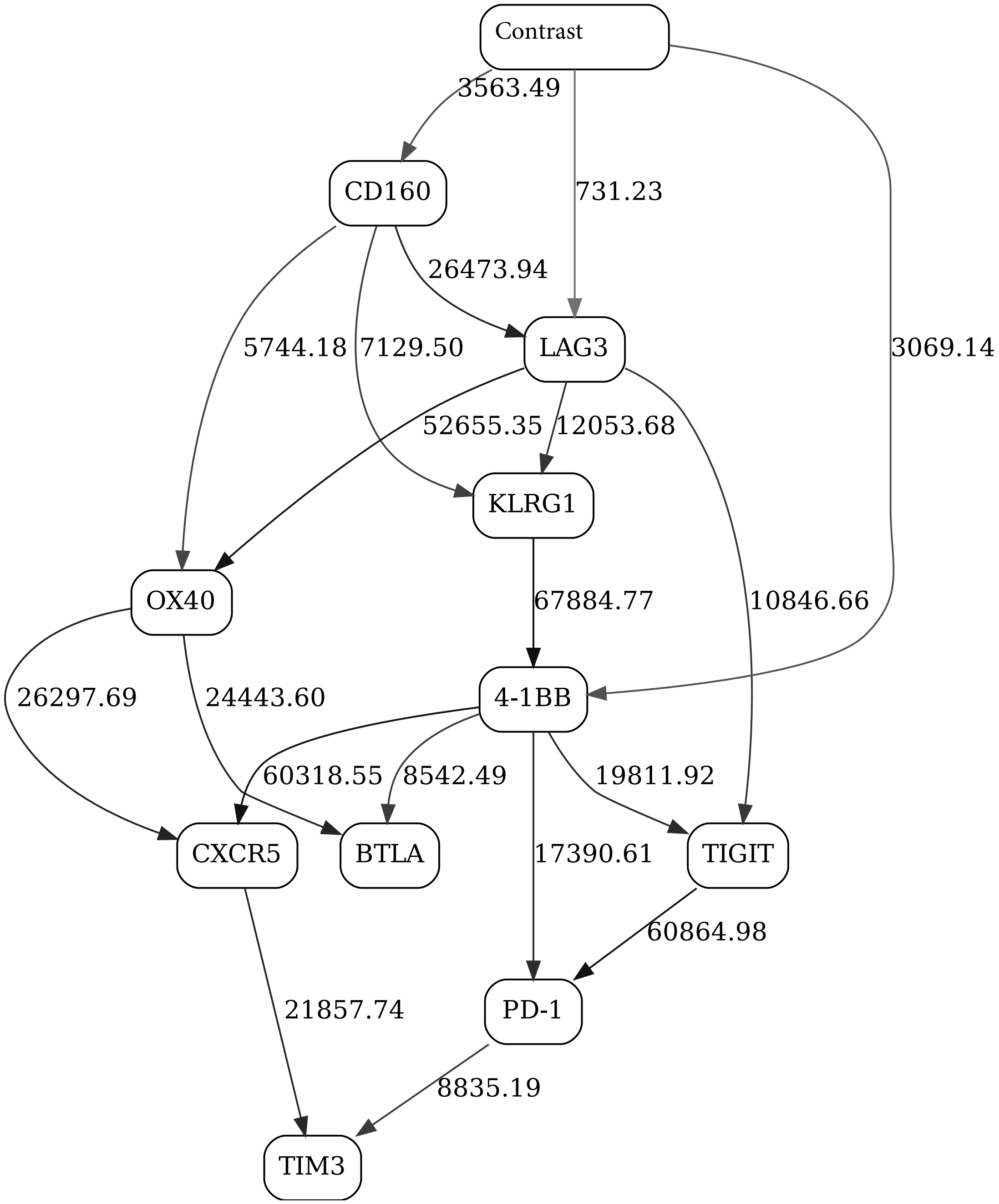


Figure 27: Checkpoint immune signaling network panel, naive CD8

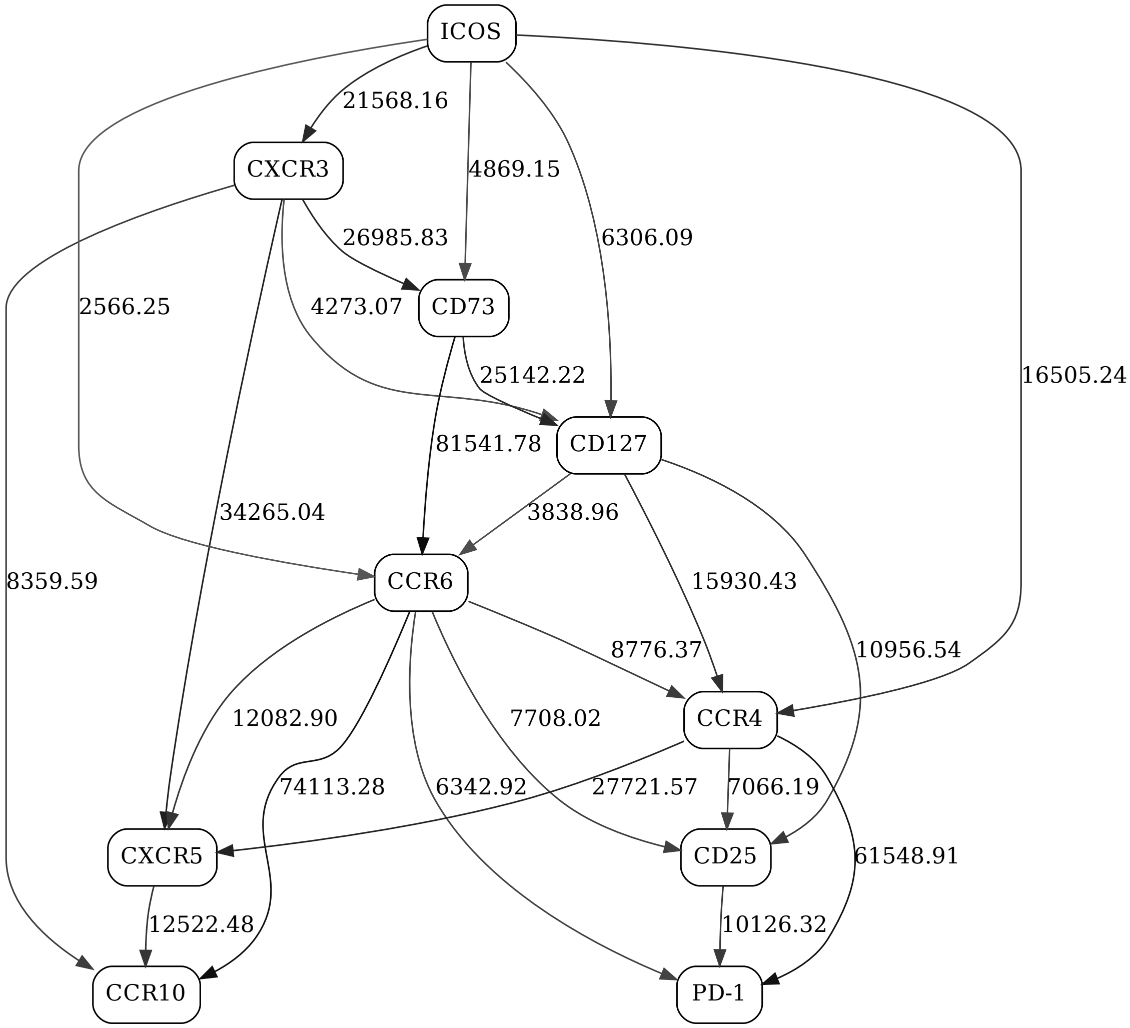


Figure 28: Adaptive immune signaling network panel, non-naive CD4, day 1, responders

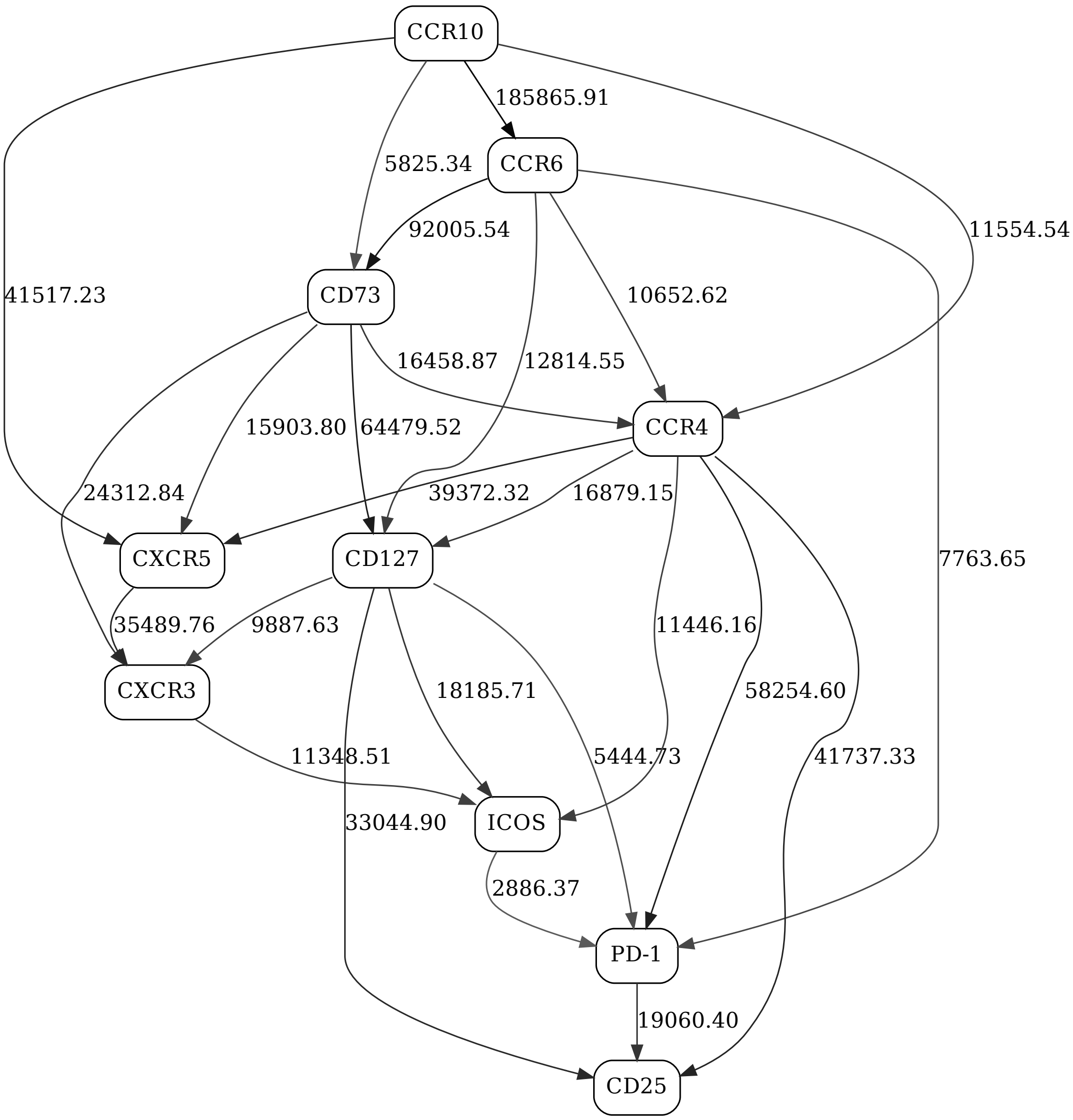


Figure 29: Adaptive immune signaling network panel, non-naive CD4, day 1, non-responders

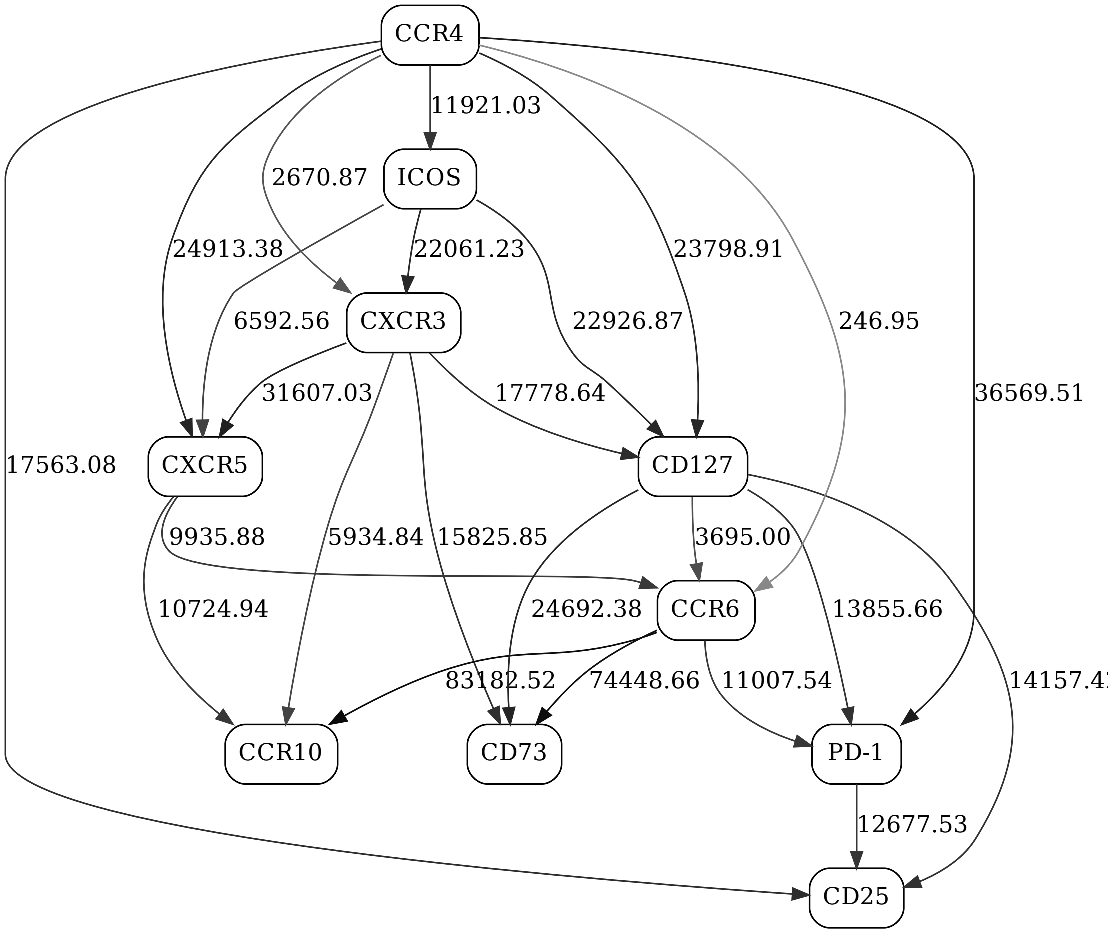


Figure 30: Adaptive immune signaling network panel, non-naive CD4, day 21, responders

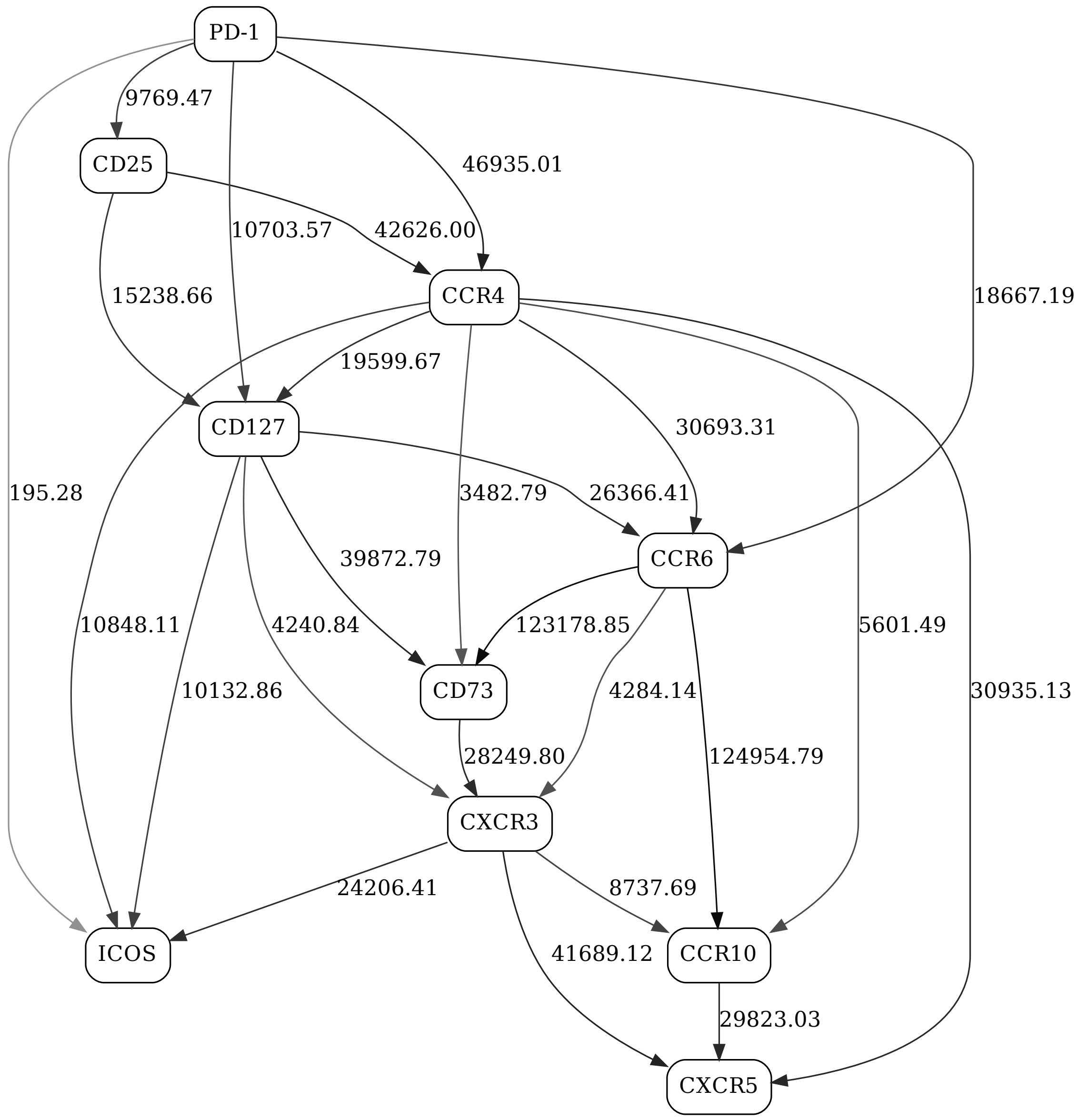


Figure 31: Adaptive immune signaling network panel, non-naive CD4, day 21, non-responders

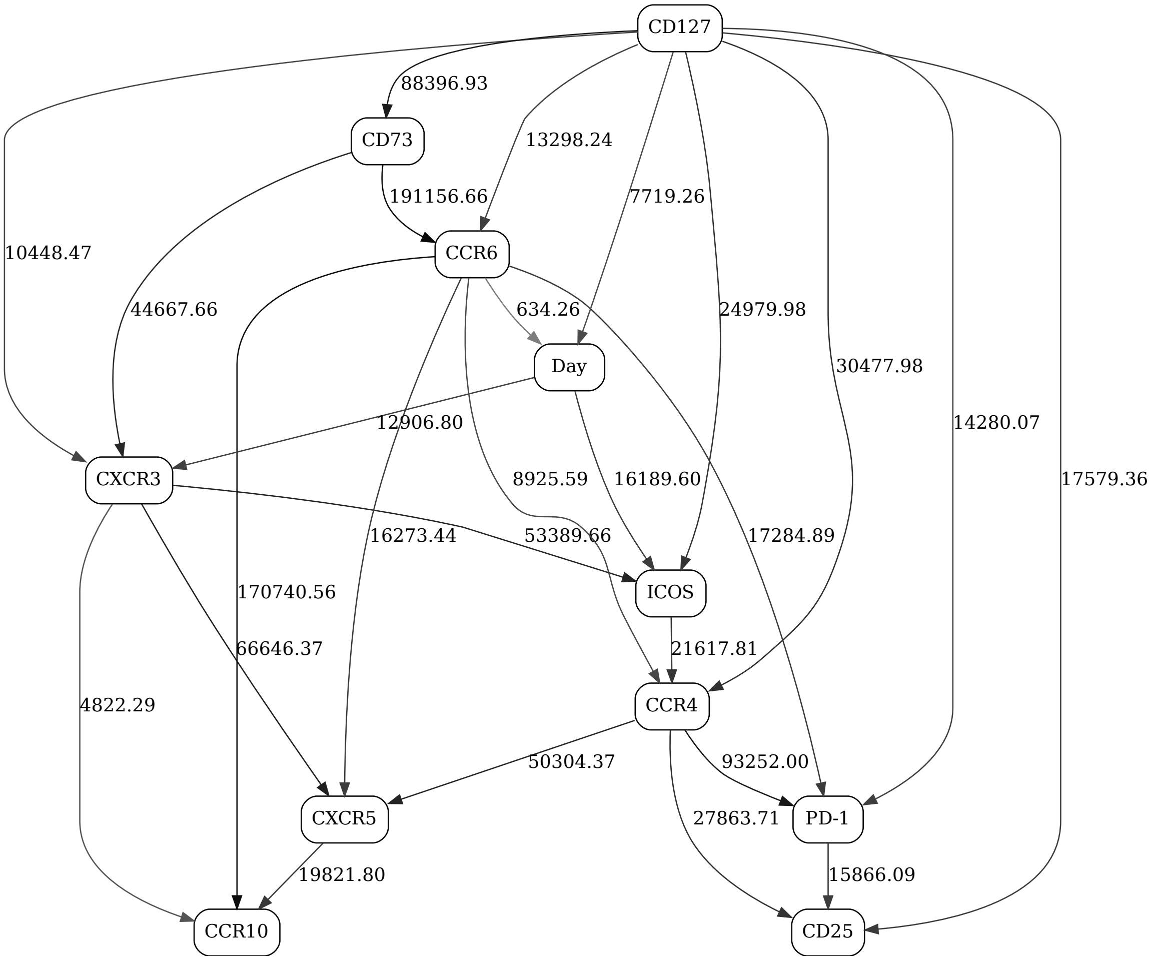


Figure 32: Adaptive immune signaling network panel, non-naive CD4, day contrast (2-state "Day" variable), responders

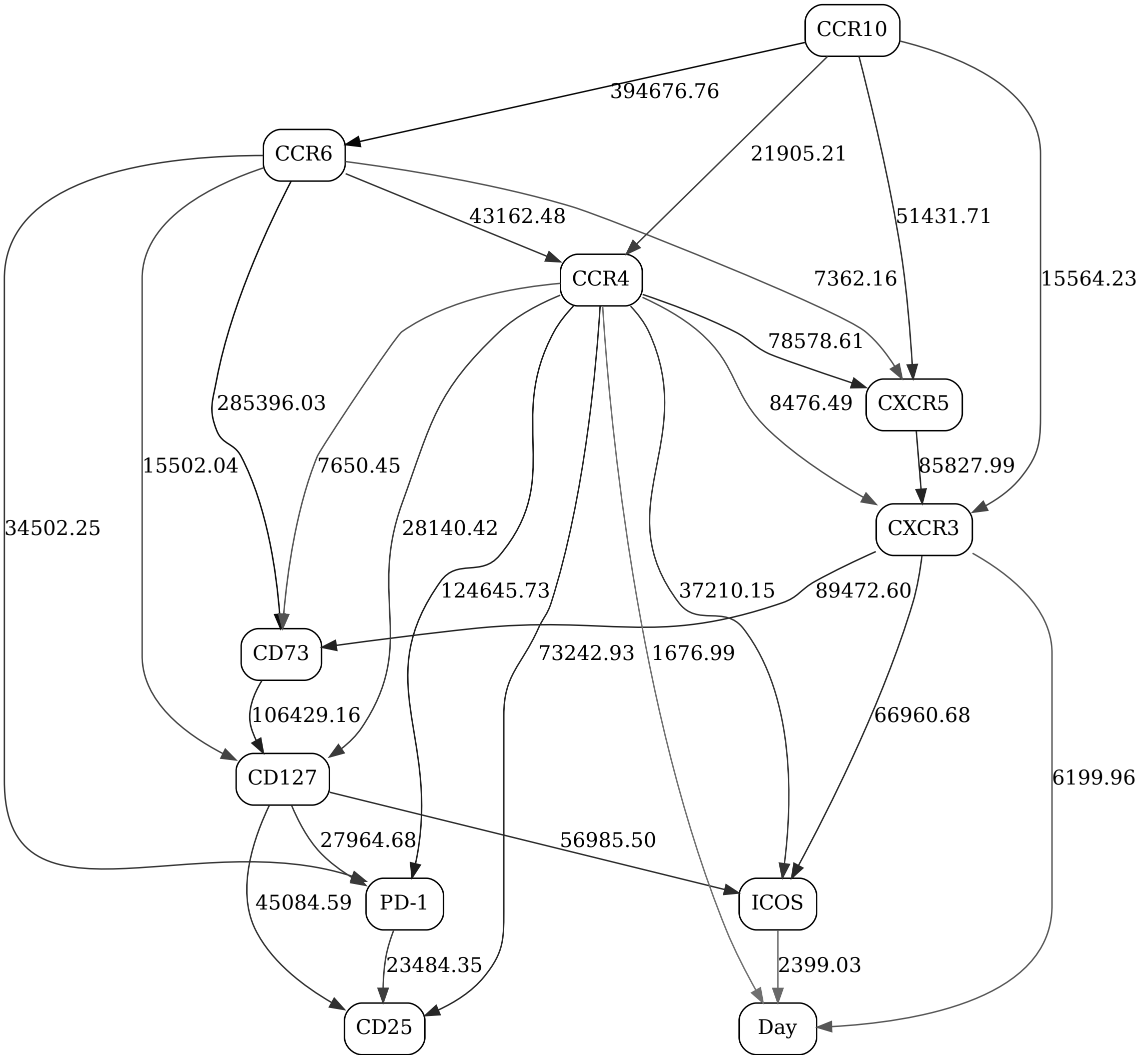


Figure 33: Adaptive immune signaling network panel, non-naive CD4, day contrast (2-state "Day" variable), non-responders

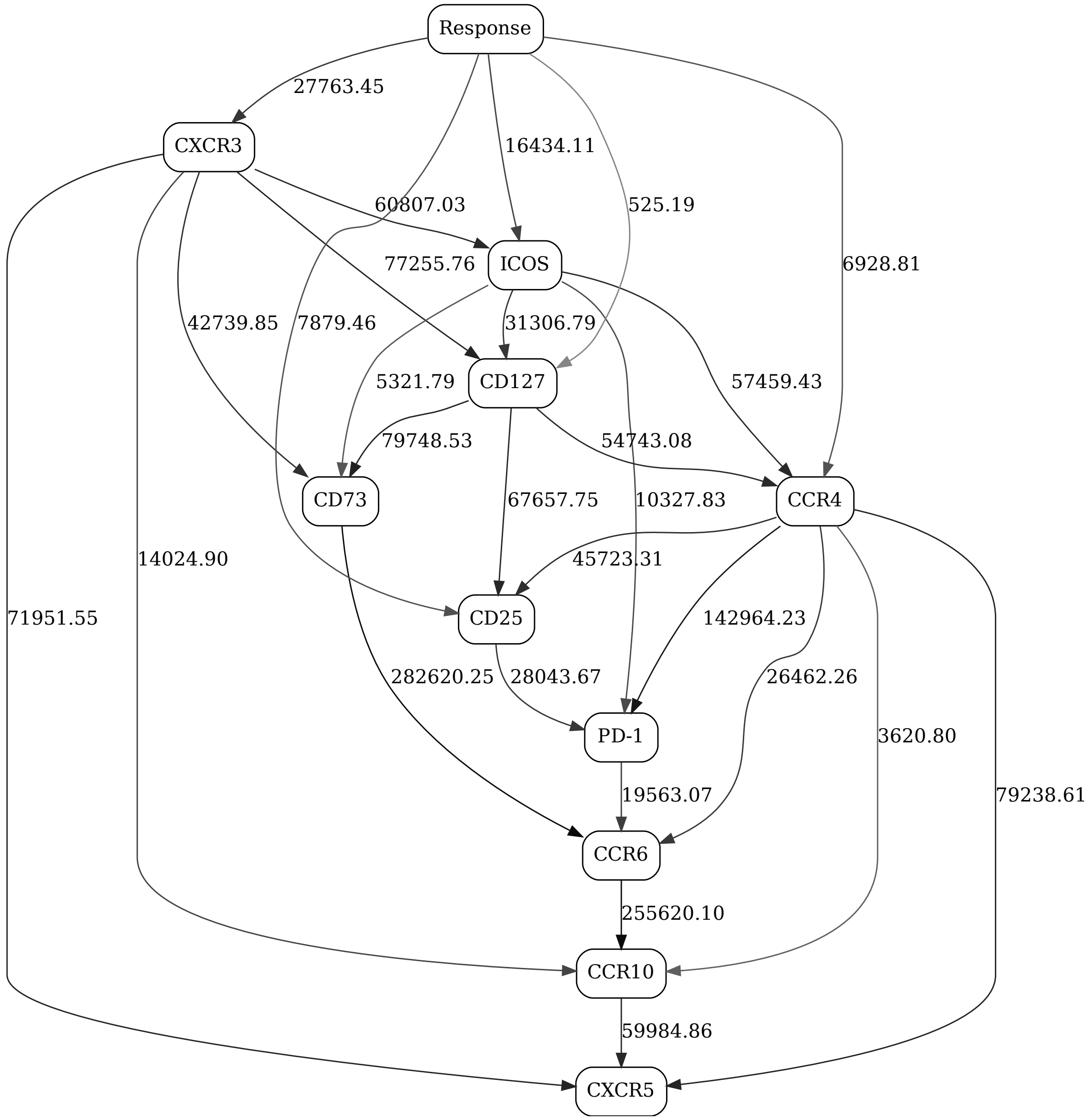


Figure 34: Adaptive immune signaling network panel, non-naive CD4, day 1, response contrast (2-state "Response" variable)

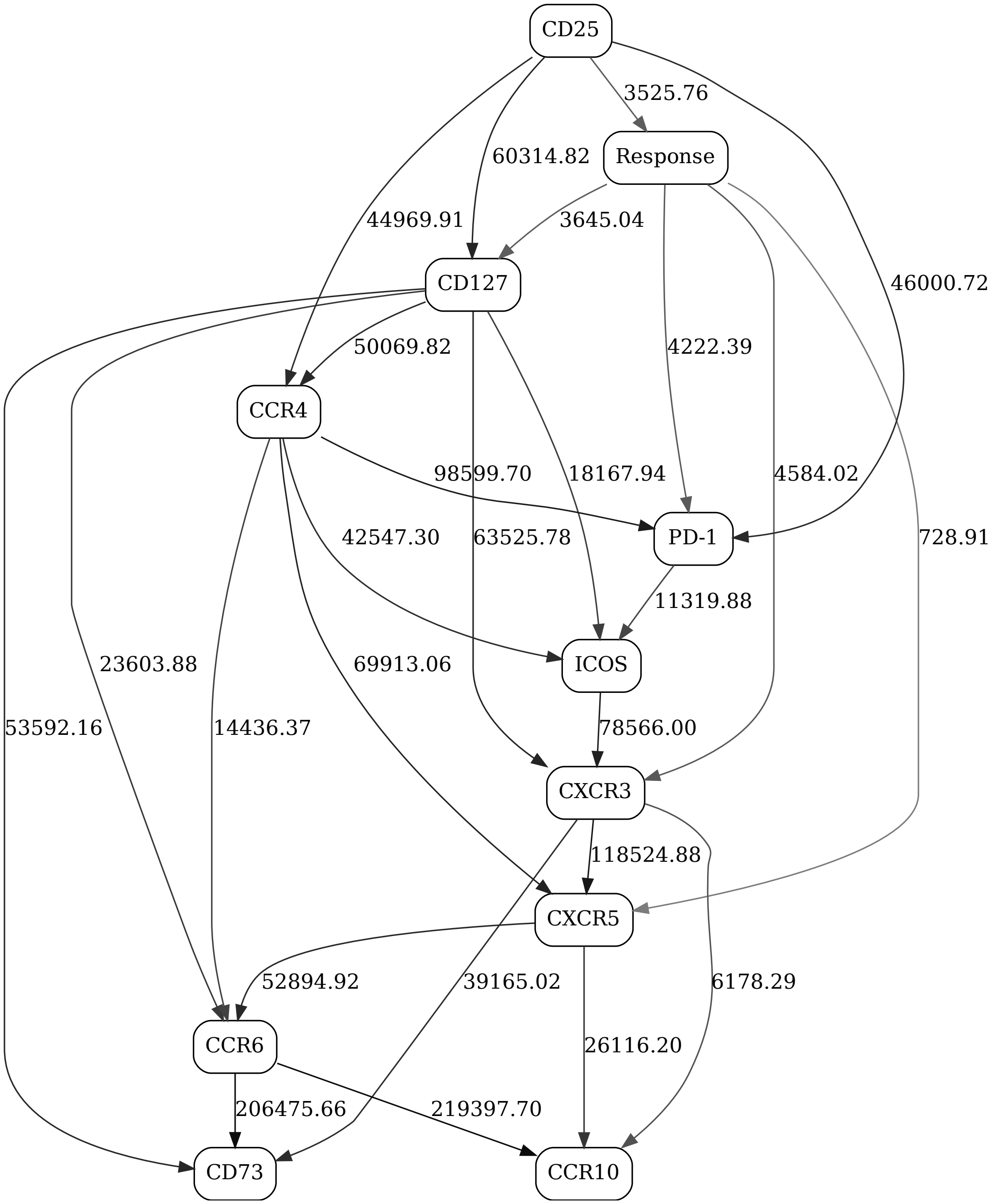


Figure 35: Adaptive immune signaling network panel, non-naive CD4, day 21, response contrast (2-state "Response" variable)

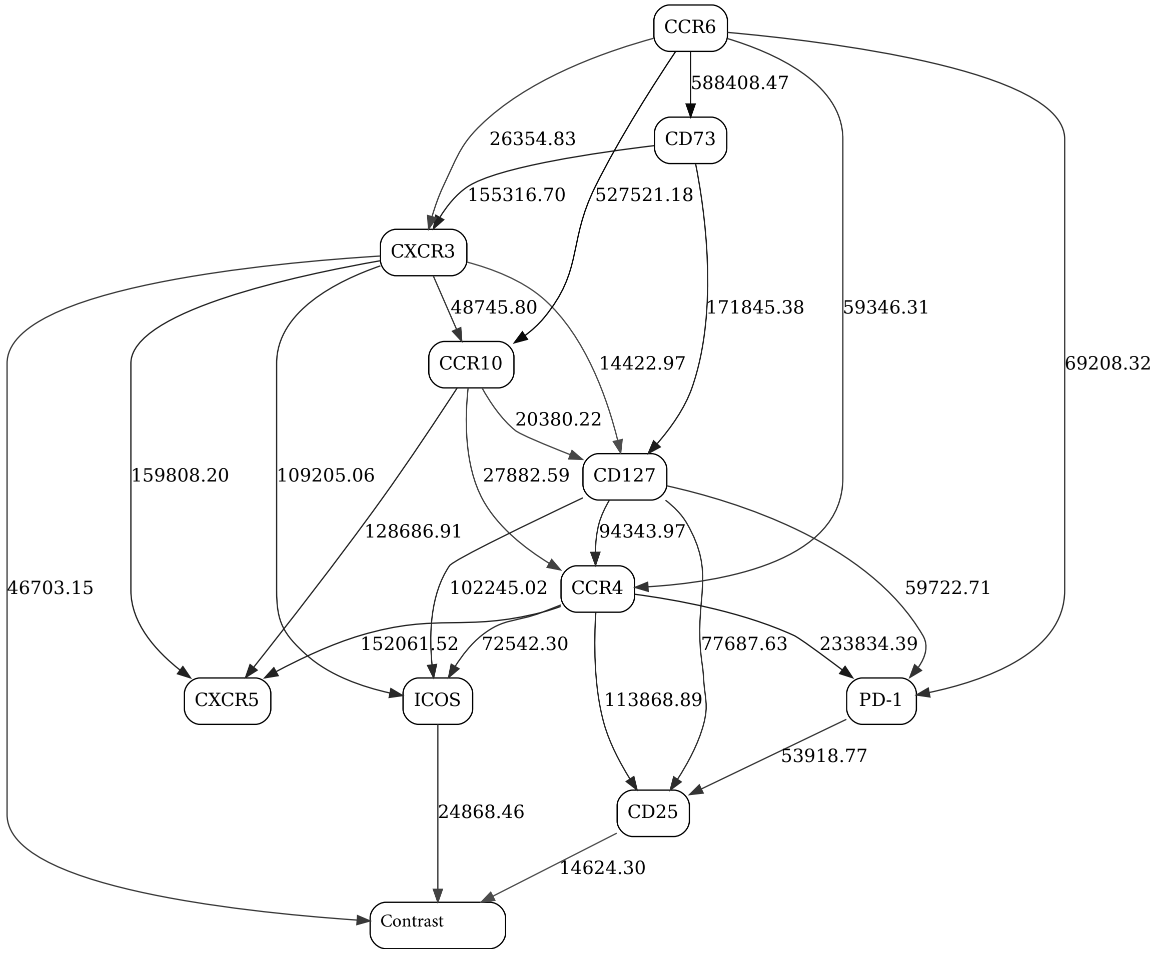


Figure 36: Adaptive immune signaling network panel, non-naive CD4

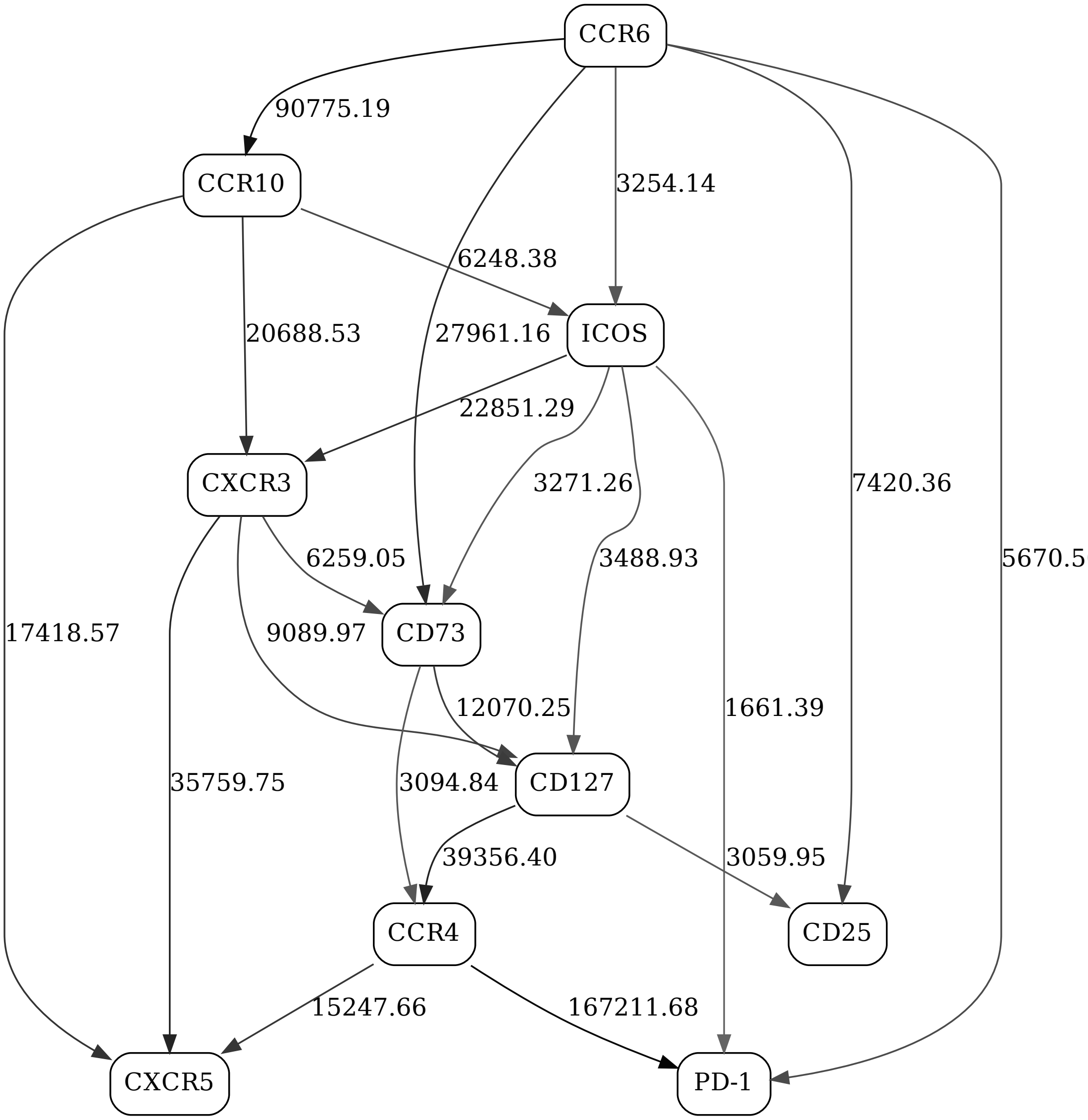


Figure 37: Adaptive immune signaling network panel, naive CD4, day 1, responders

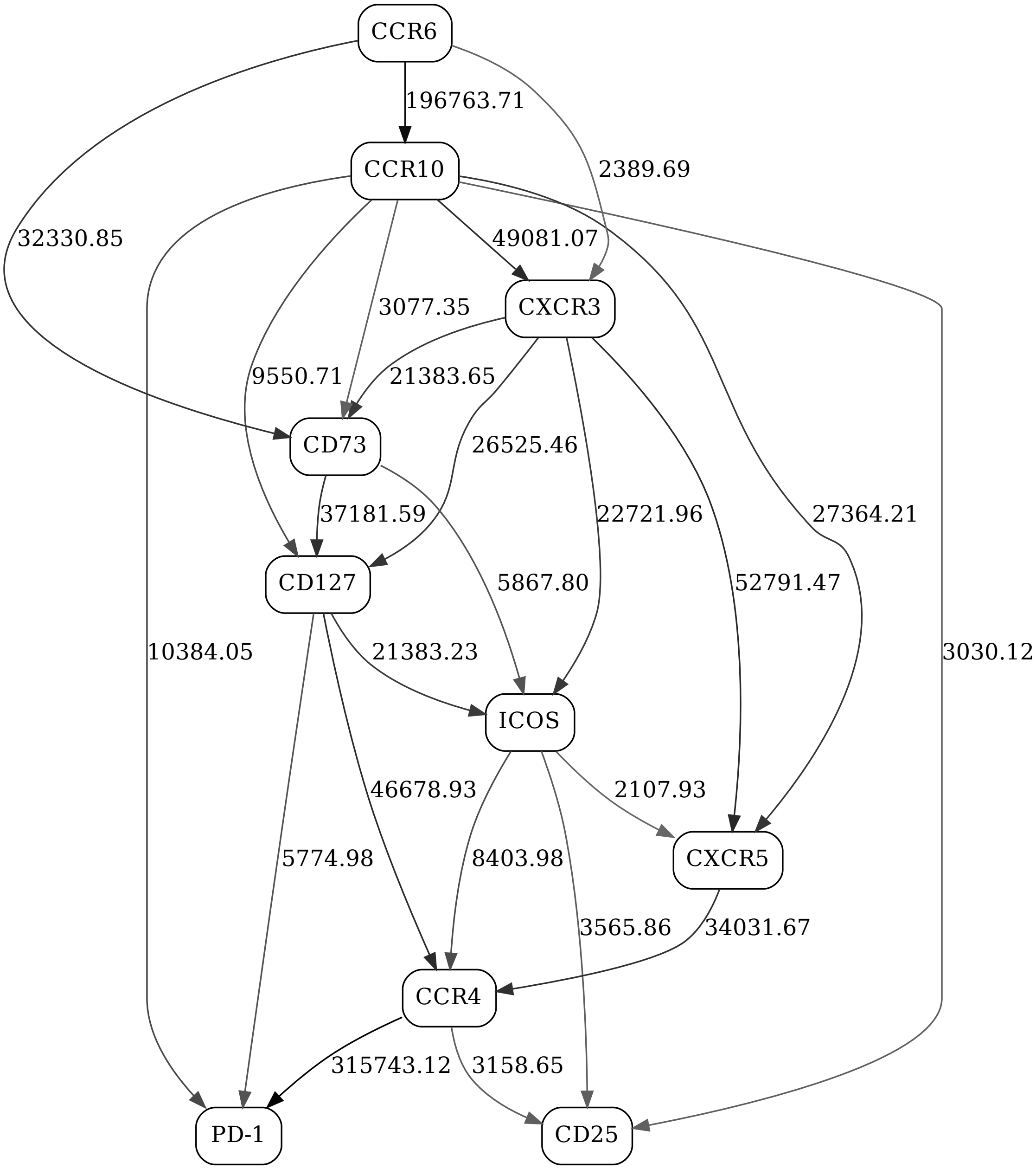


Figure 38: Adaptive immune signaling network panel, naive CD4, day 1, non-responders

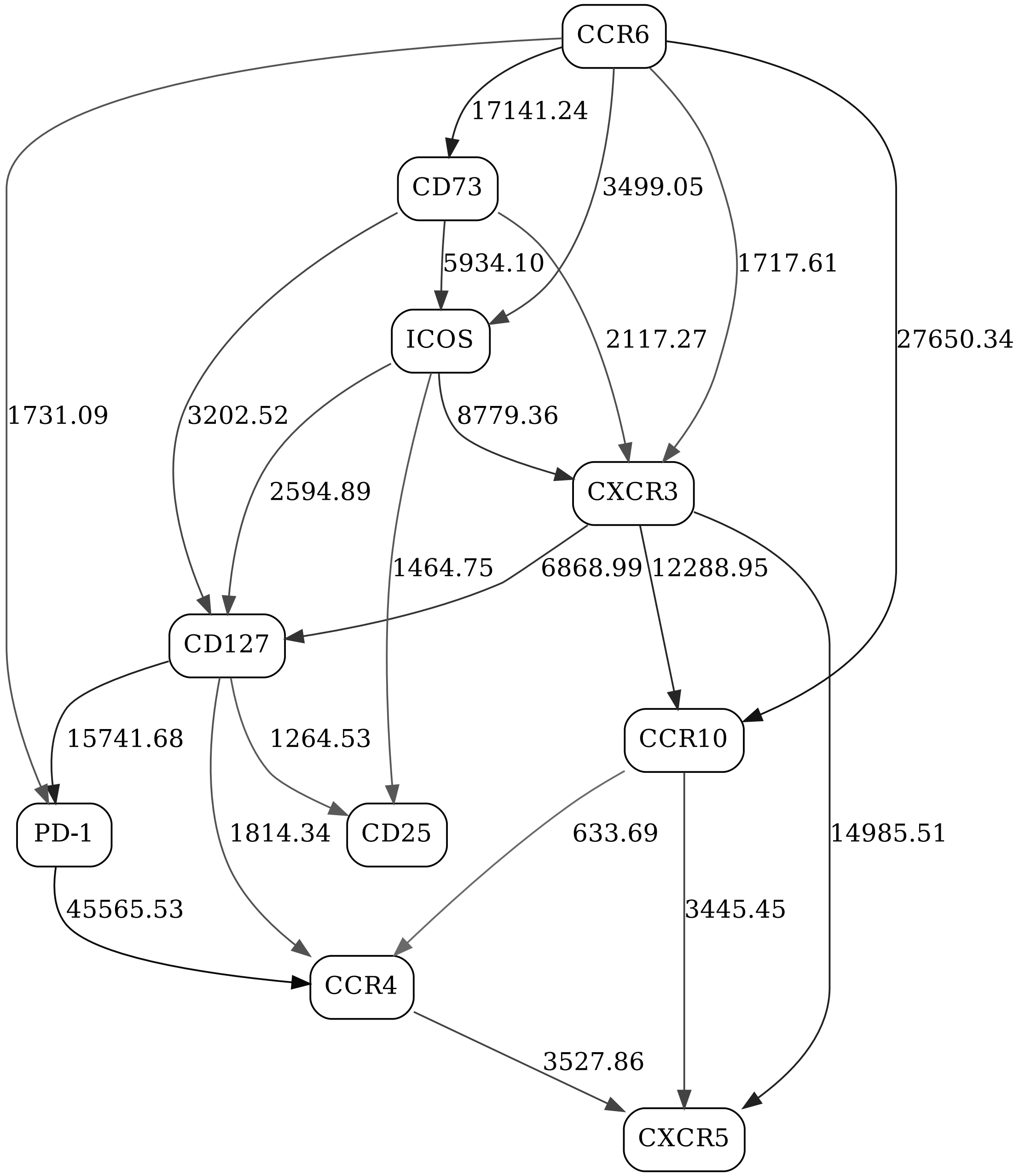


Figure 39: Adaptive immune signaling network panel, naive CD4, day 21, responders

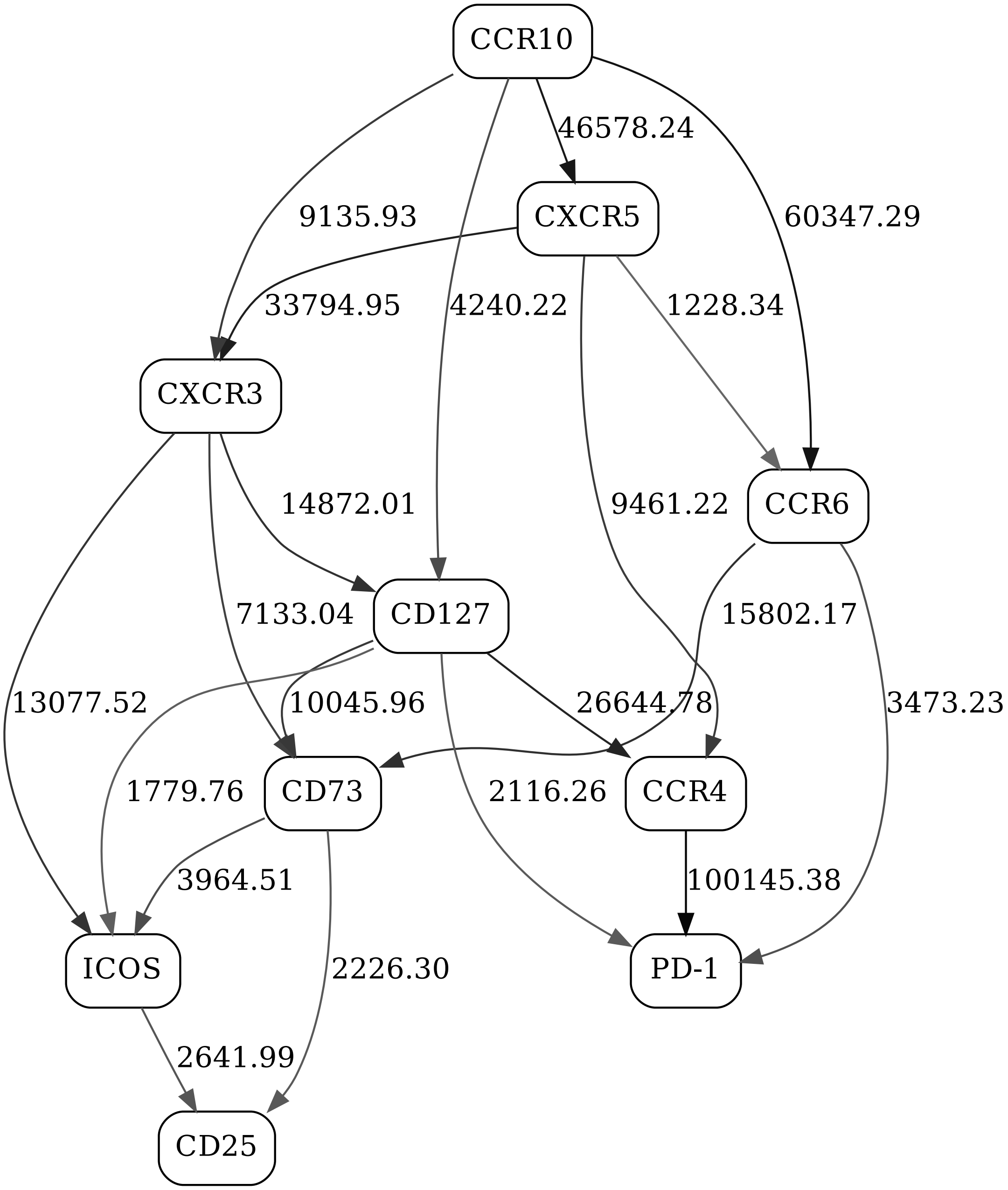


Figure 40: Adaptive immune signaling network panel, naive CD4, day 21, non-responders

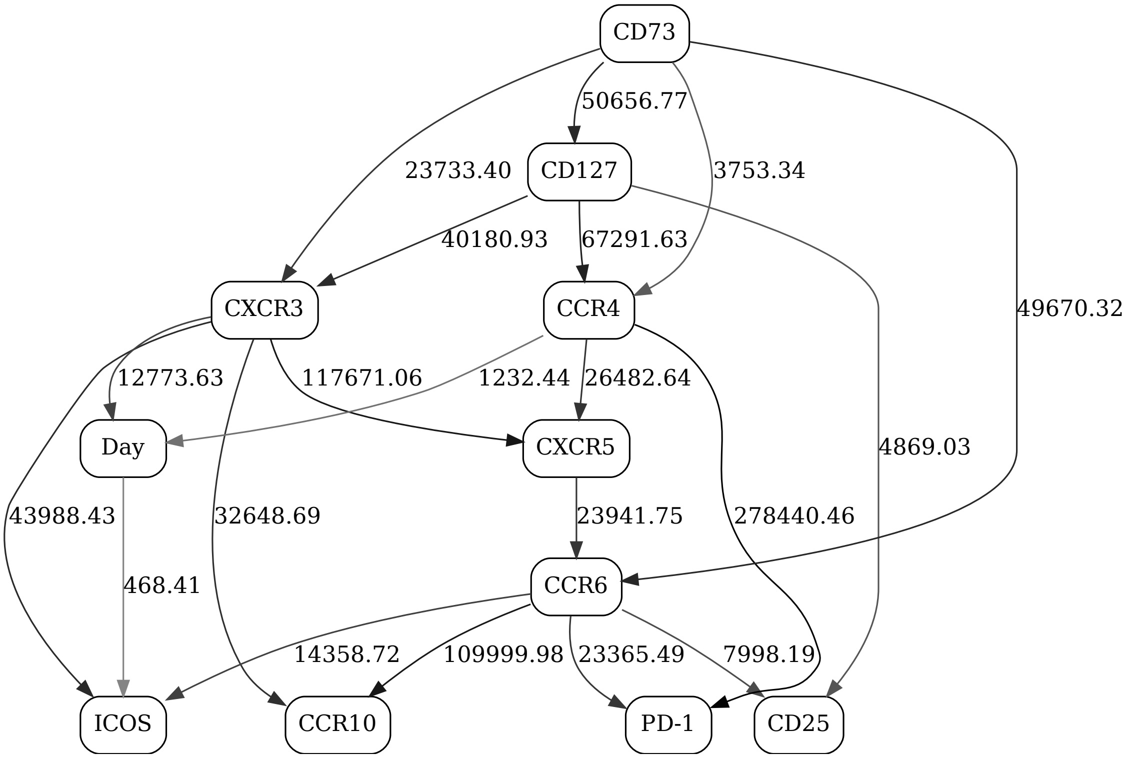


Figure 41: Adaptive immune signaling network panel, naive CD4, day contrast (2-state "Day" variable), responders

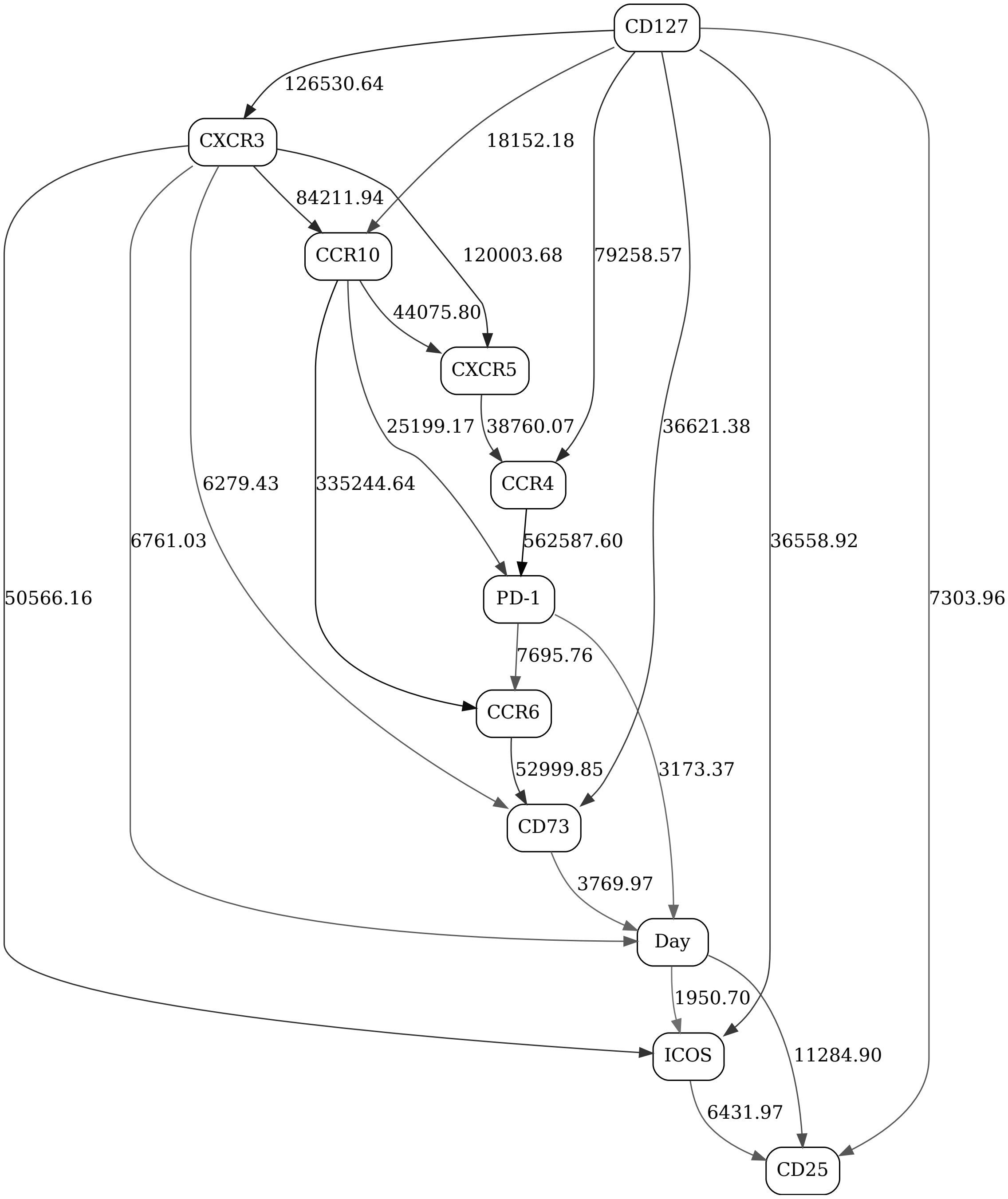


Figure 42: Adaptive immune signaling network panel, naive CD4, day contrast (2-state "Day" variable), non-responders

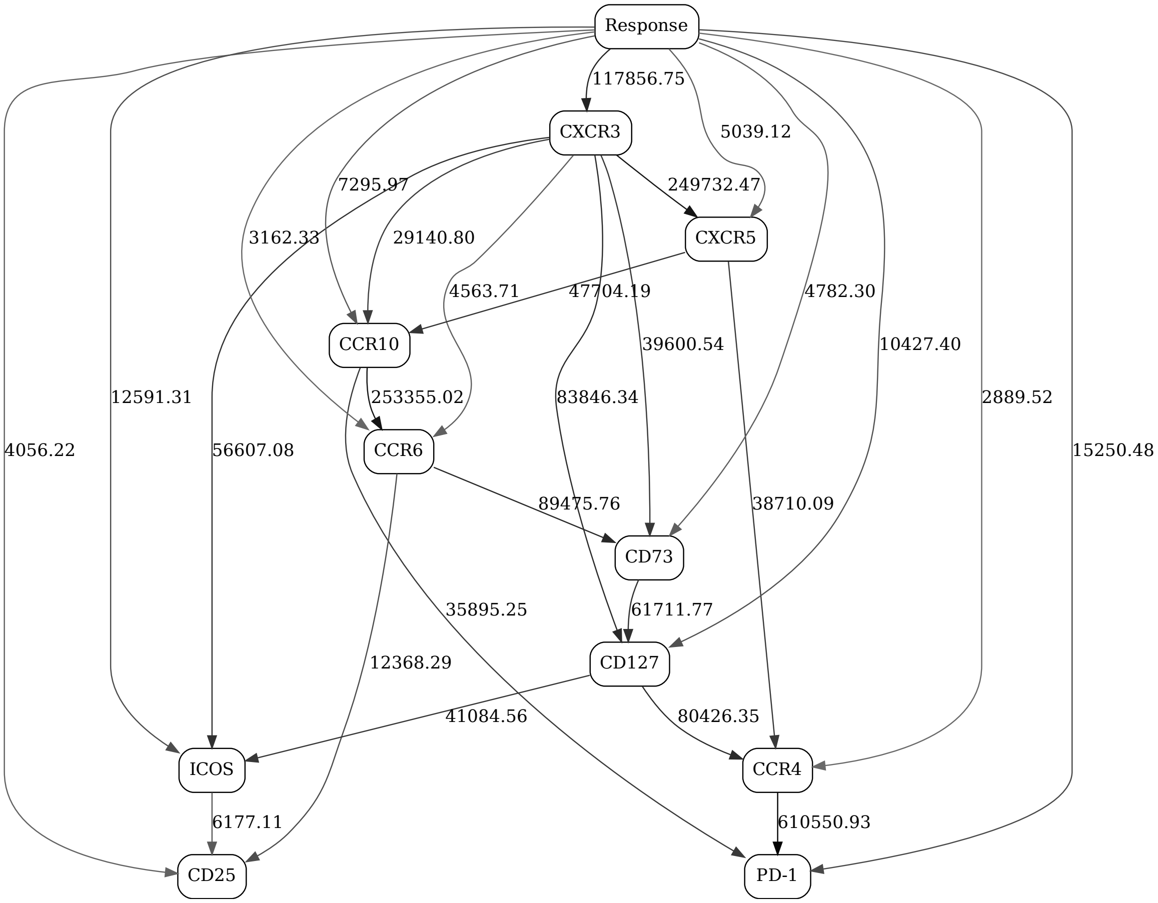


Figure 43: Adaptive immune signaling network panel, naive CD4, day 1, response contrast (2-state "Response" variable)

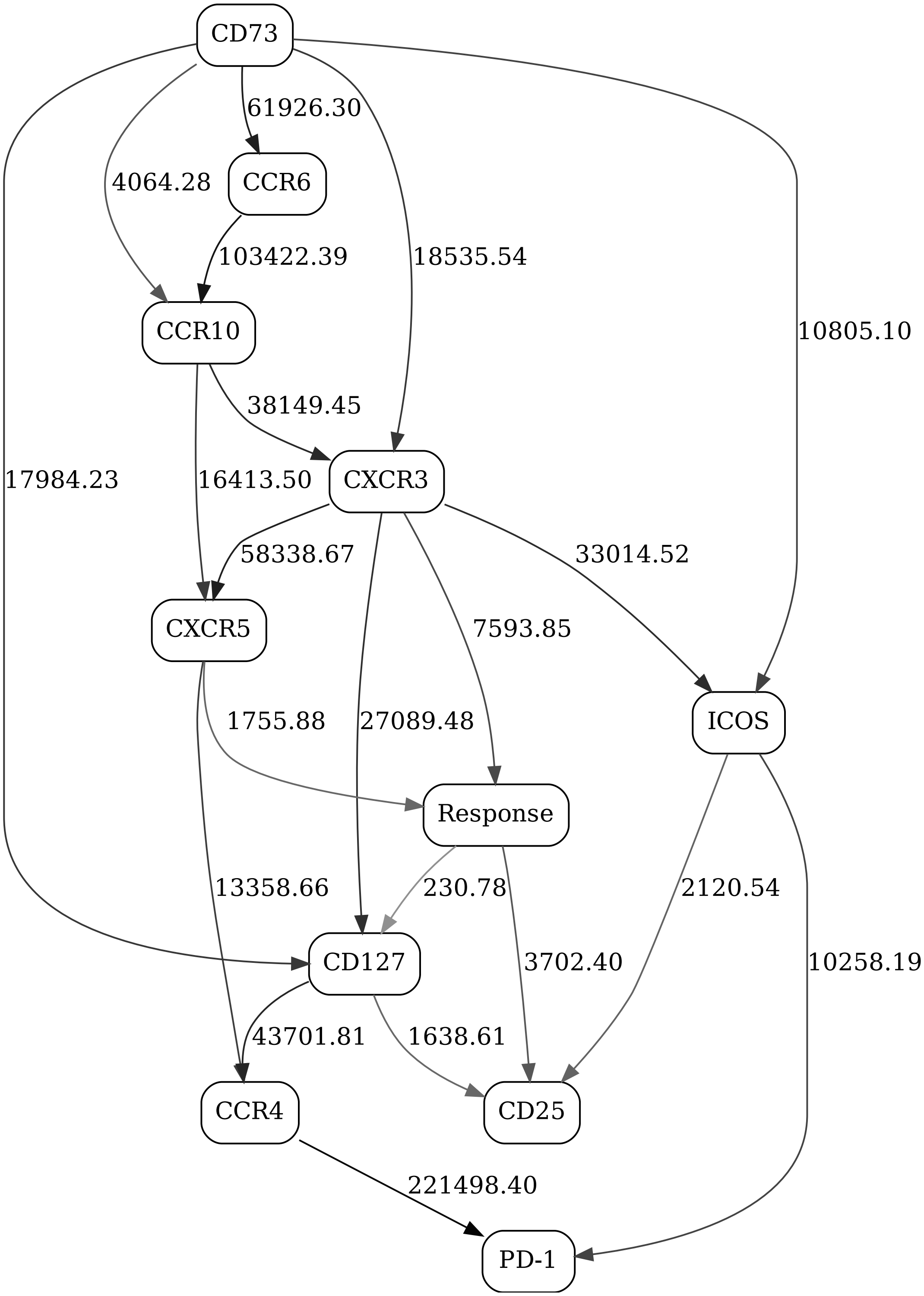


Figure 44: Adaptive immune signaling network panel, naive CD4, day 21, response contrast (2-state ”Response” variable)

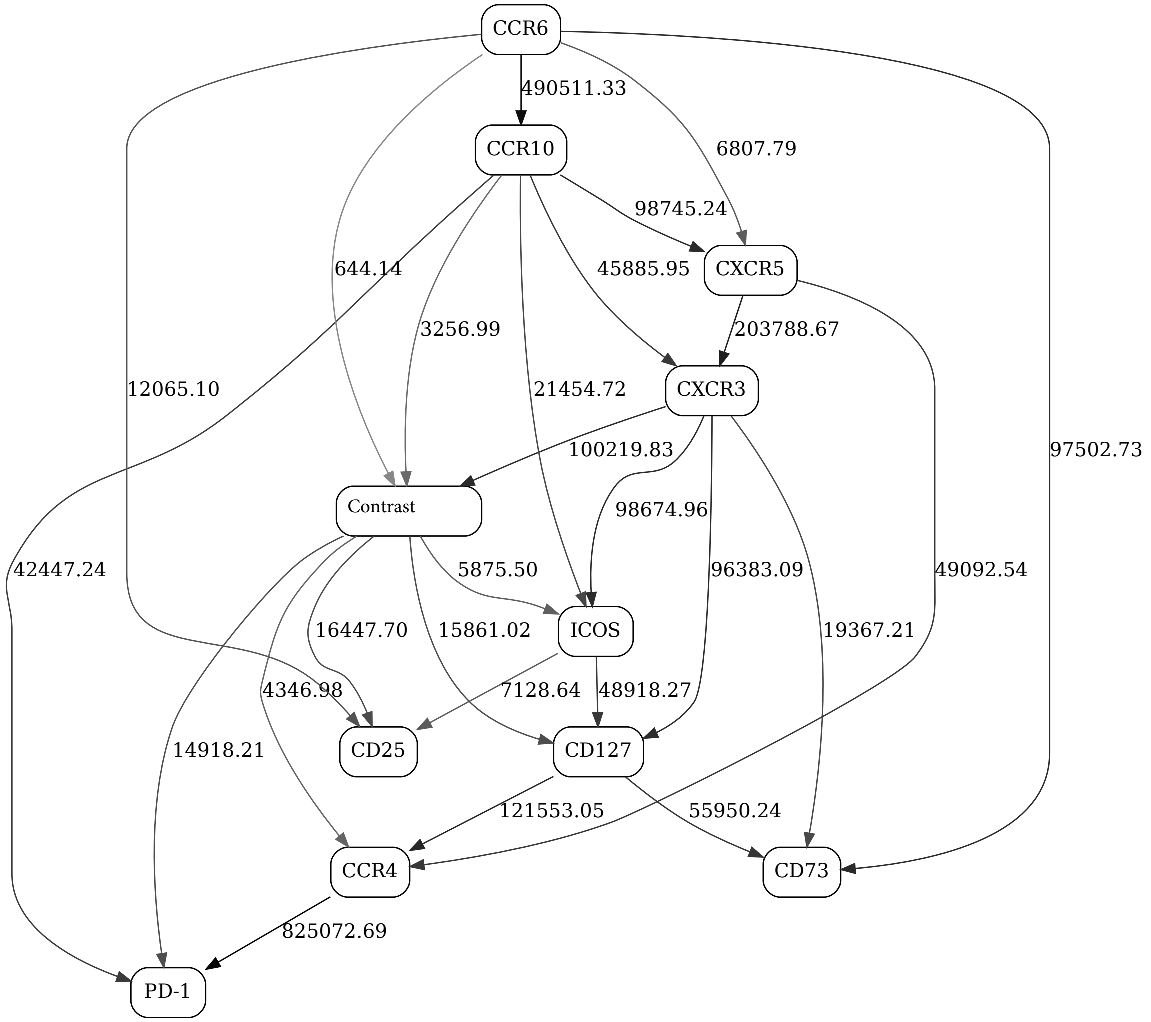


Figure 45: Adaptive immune signaling network panel, naive CD4

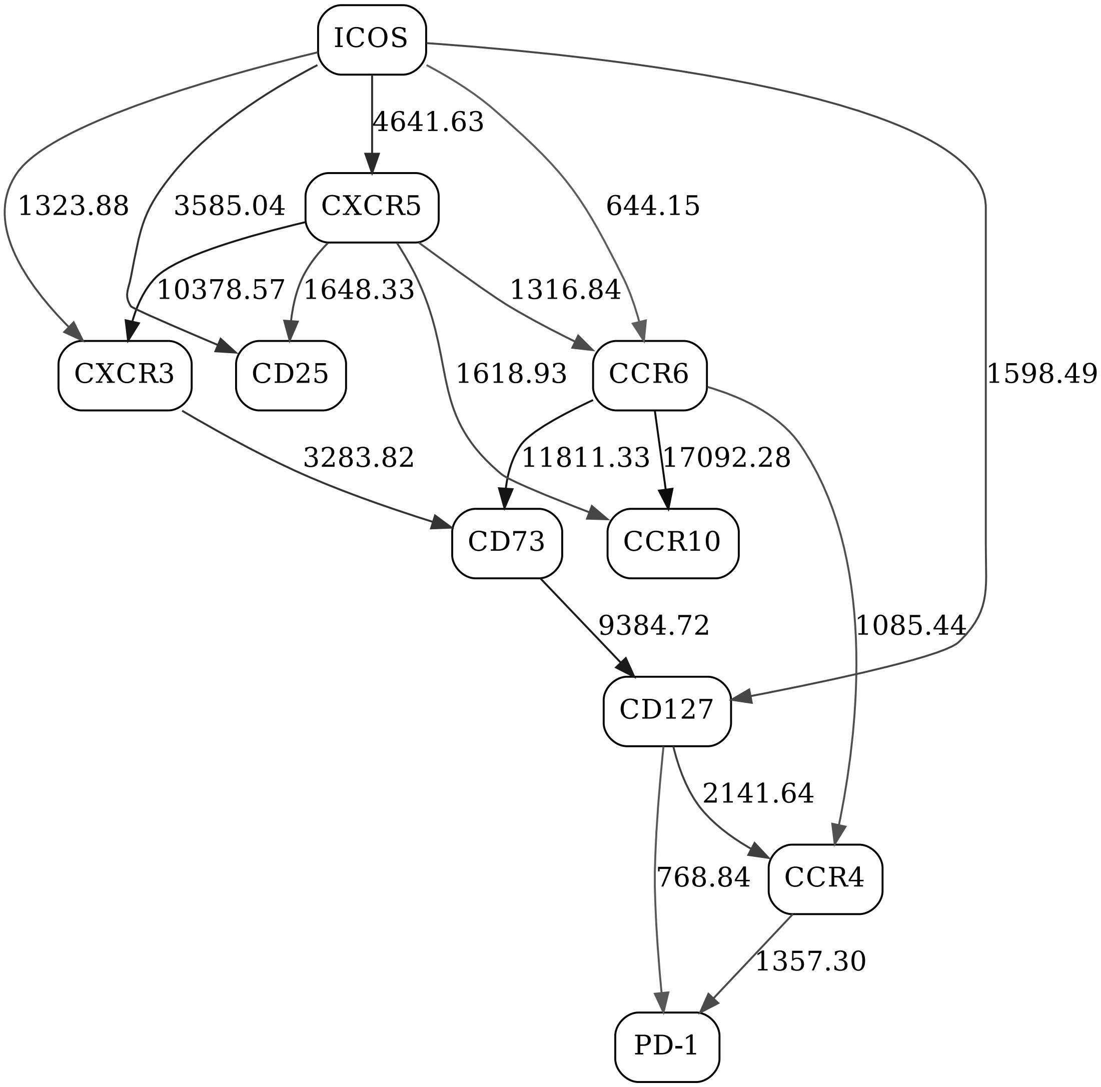


Figure 46: Adaptive immune signaling network panel, non-naive CD8, day 1, responders

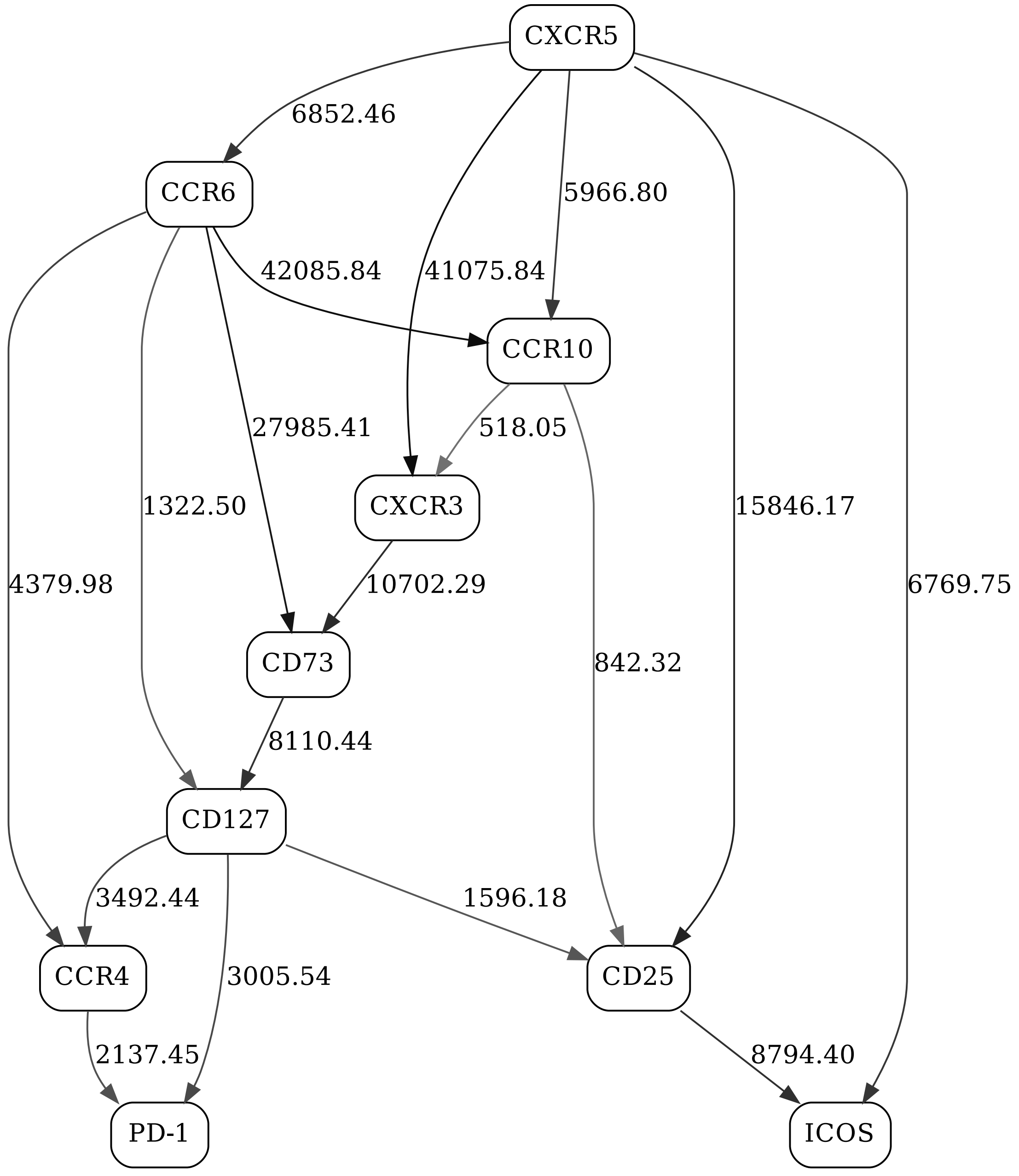


Figure 47: Adaptive immune signaling network panel, non-naive CD8, day 1, non-responders

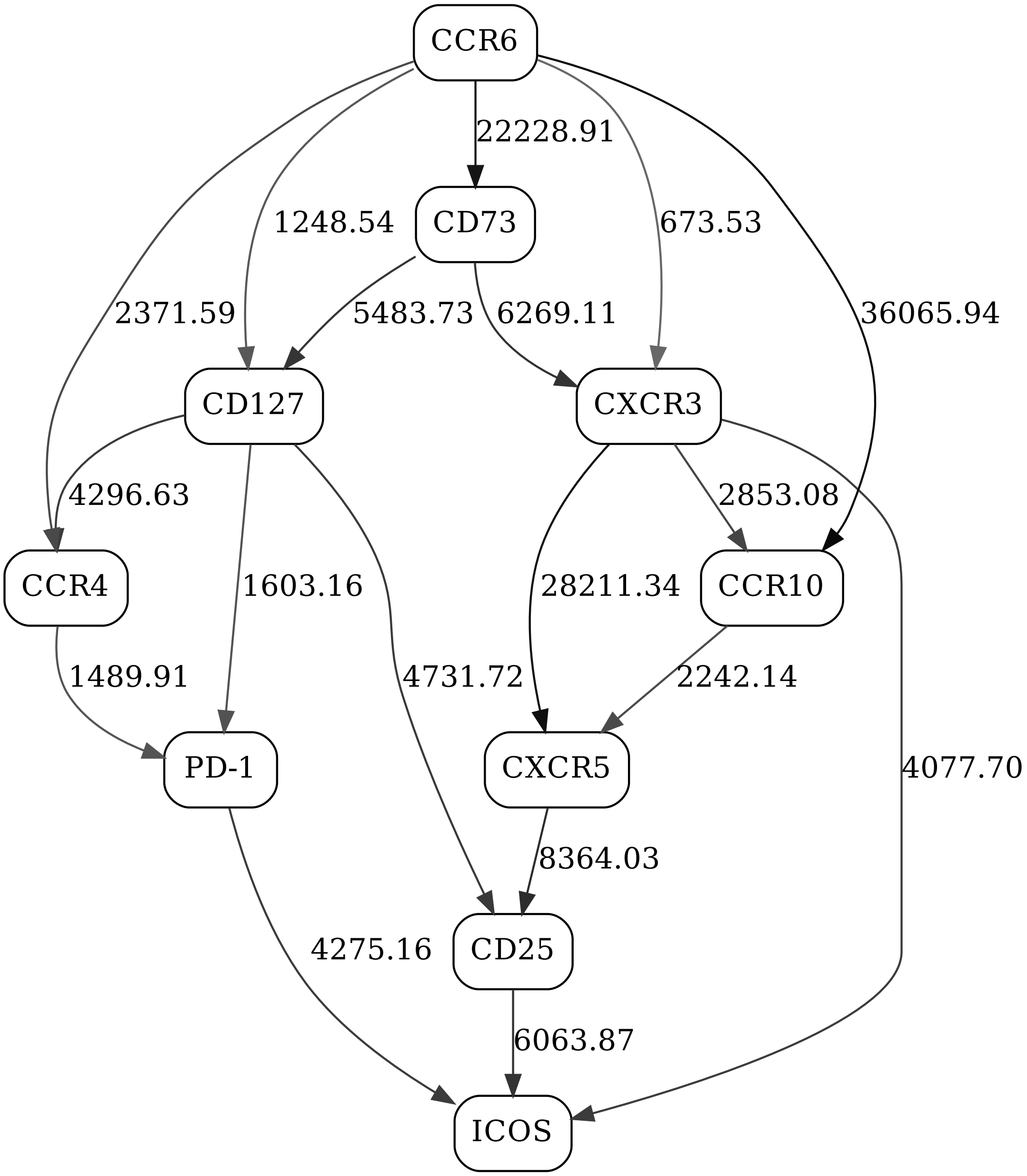


Figure 48: Adaptive immune signaling network panel, non-naive CD8, day 21, responders

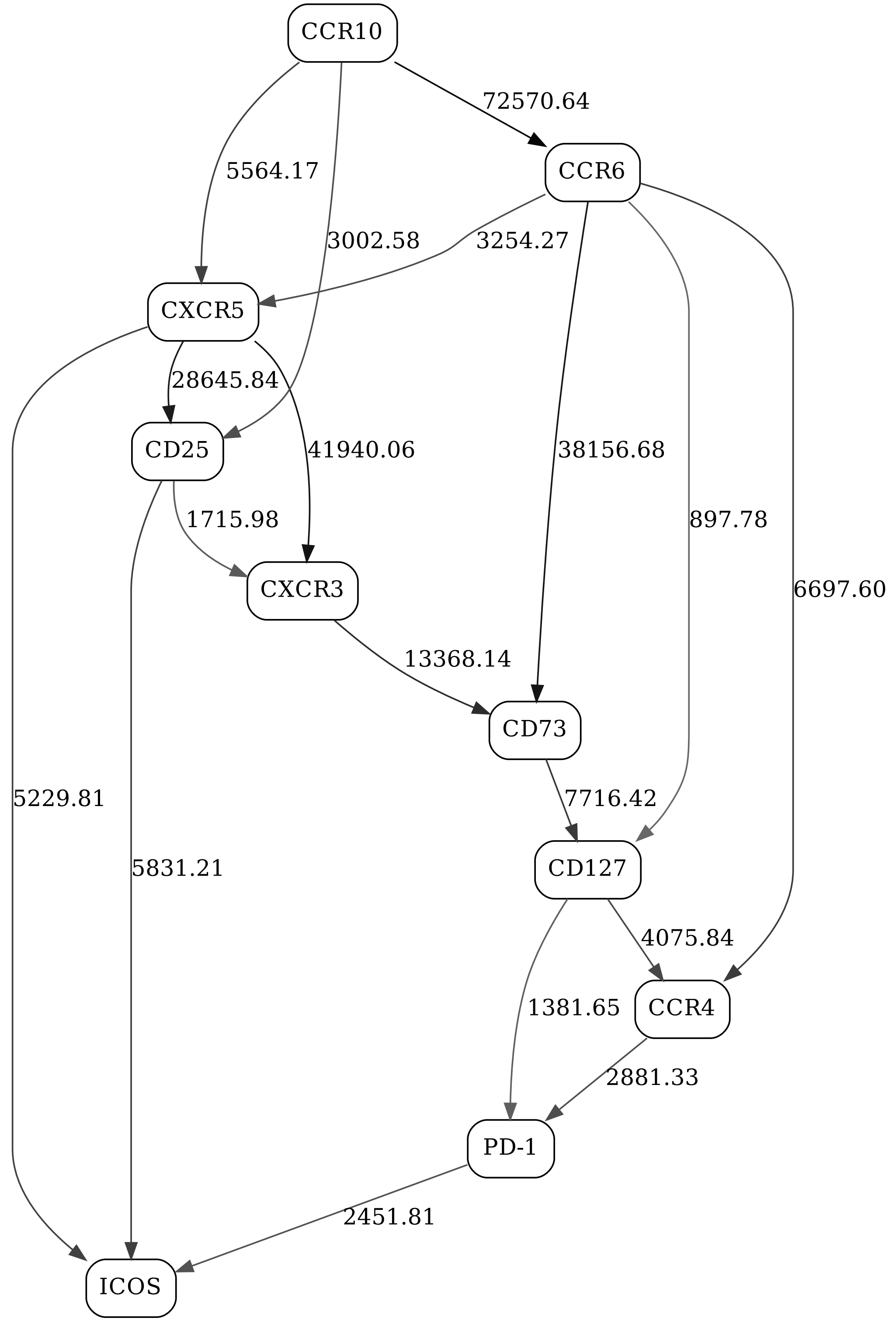


Figure 49: Adaptive immune signaling network panel, non-naive CD8, day 21, non-responders

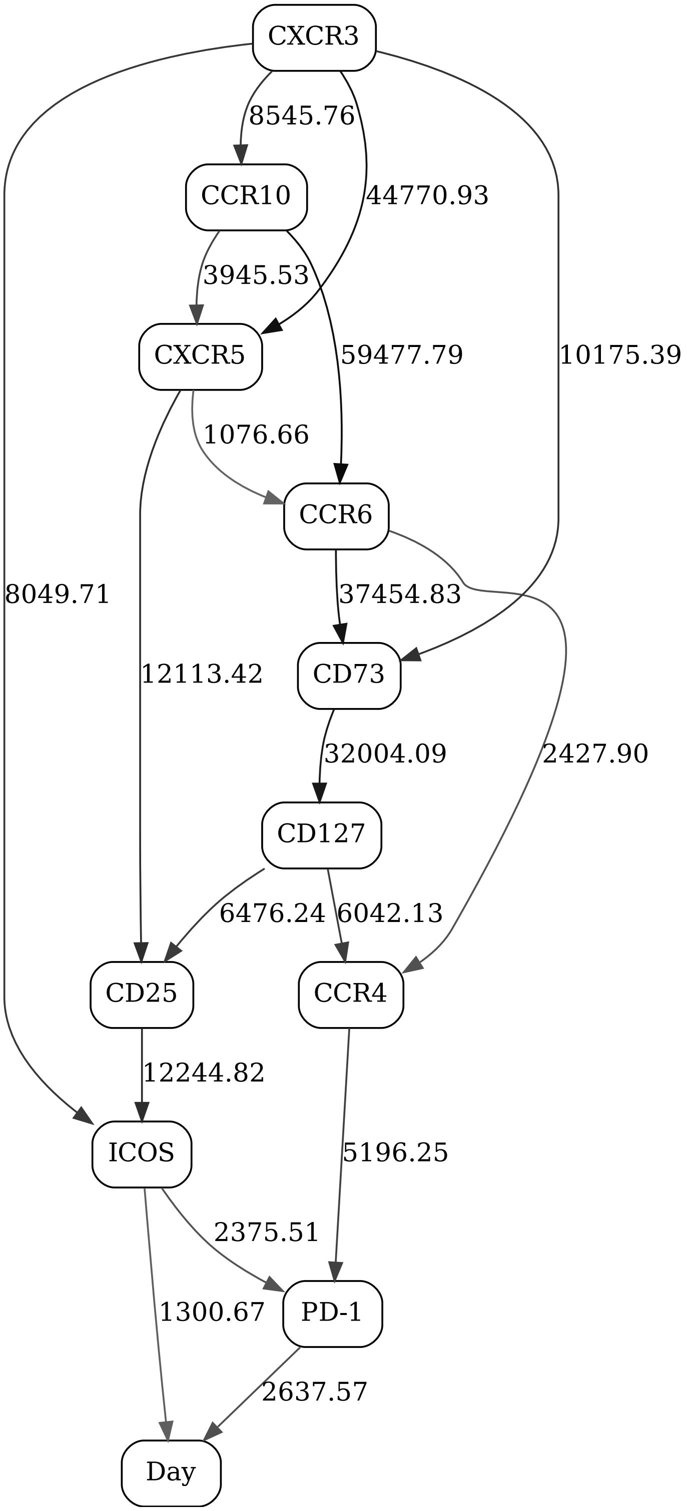


Figure 50: Adaptive immune signaling network panel, non-naive CD8, day contrast (2-state "Day" variable), responders

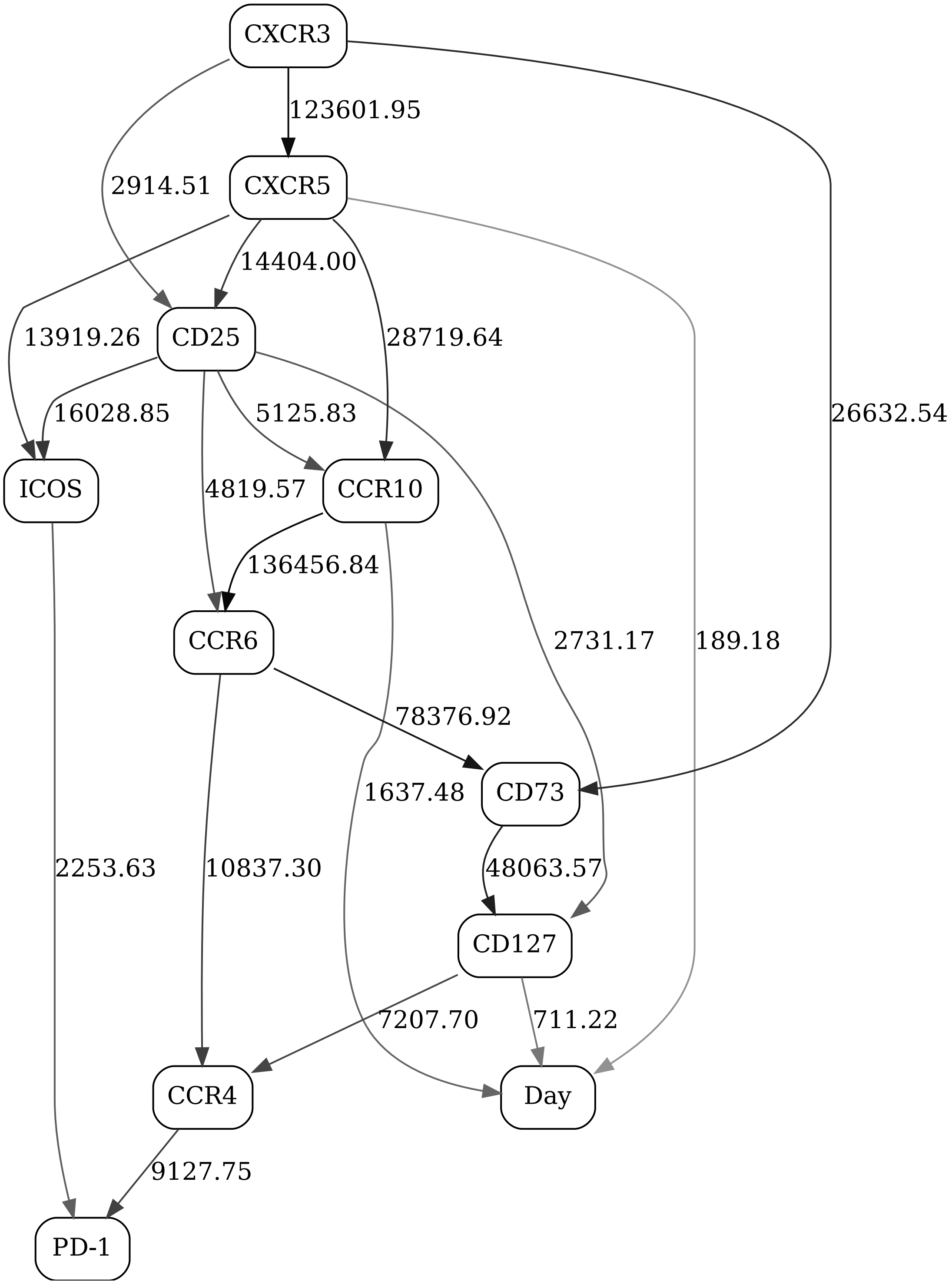


Figure 51: Adaptive immune signaling network panel, non-naive CD8, day contrast (2-state "Day" variable), non-responders

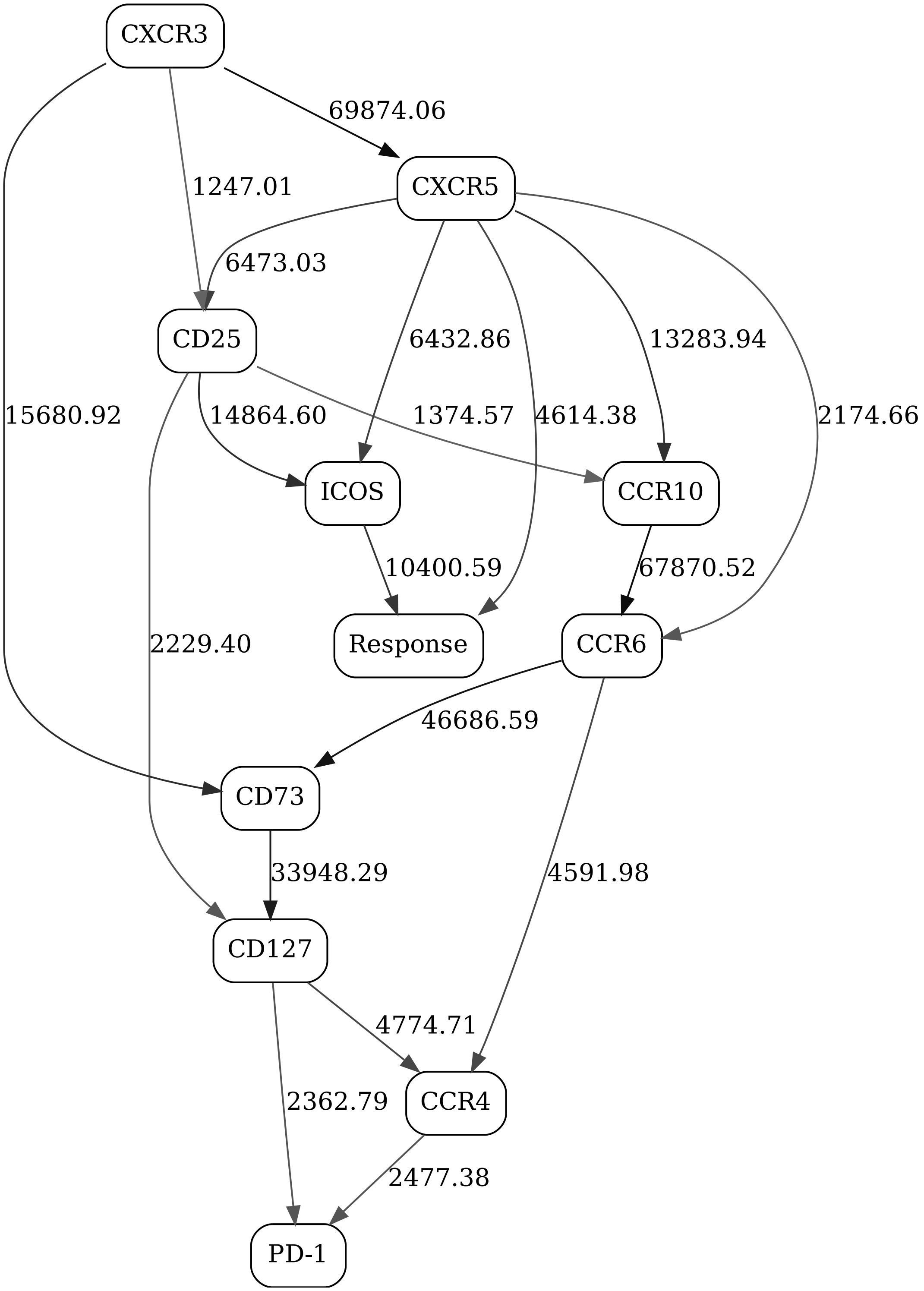


Figure 52: Adaptive immune signaling network panel, non-naive CD8, day 1, response contrast (2-state ”Response” variable)

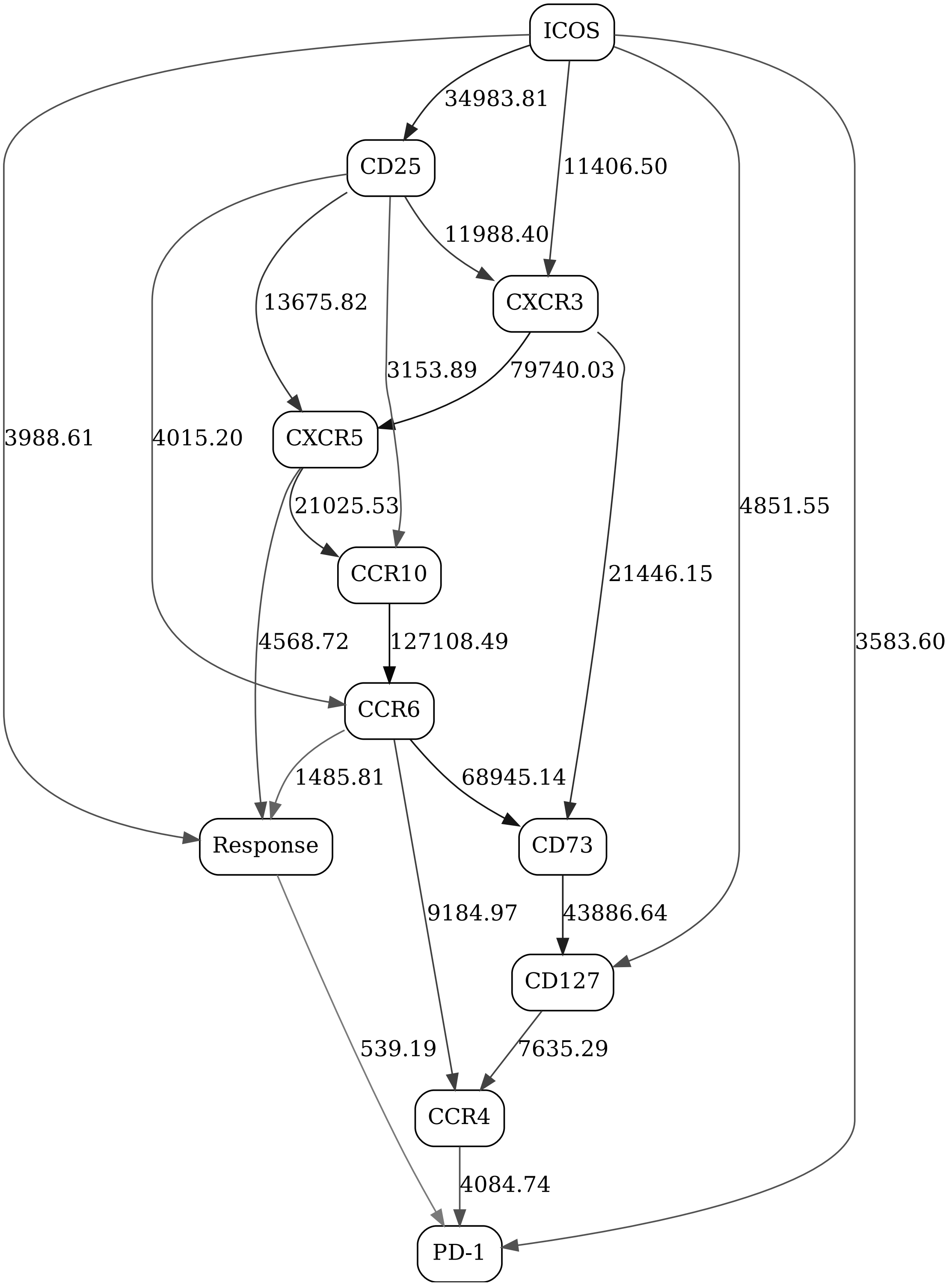


Figure 53: Adaptive immune signaling network panel, non-naive CD8, day 21, response contrast (2-state ”Response” variable)

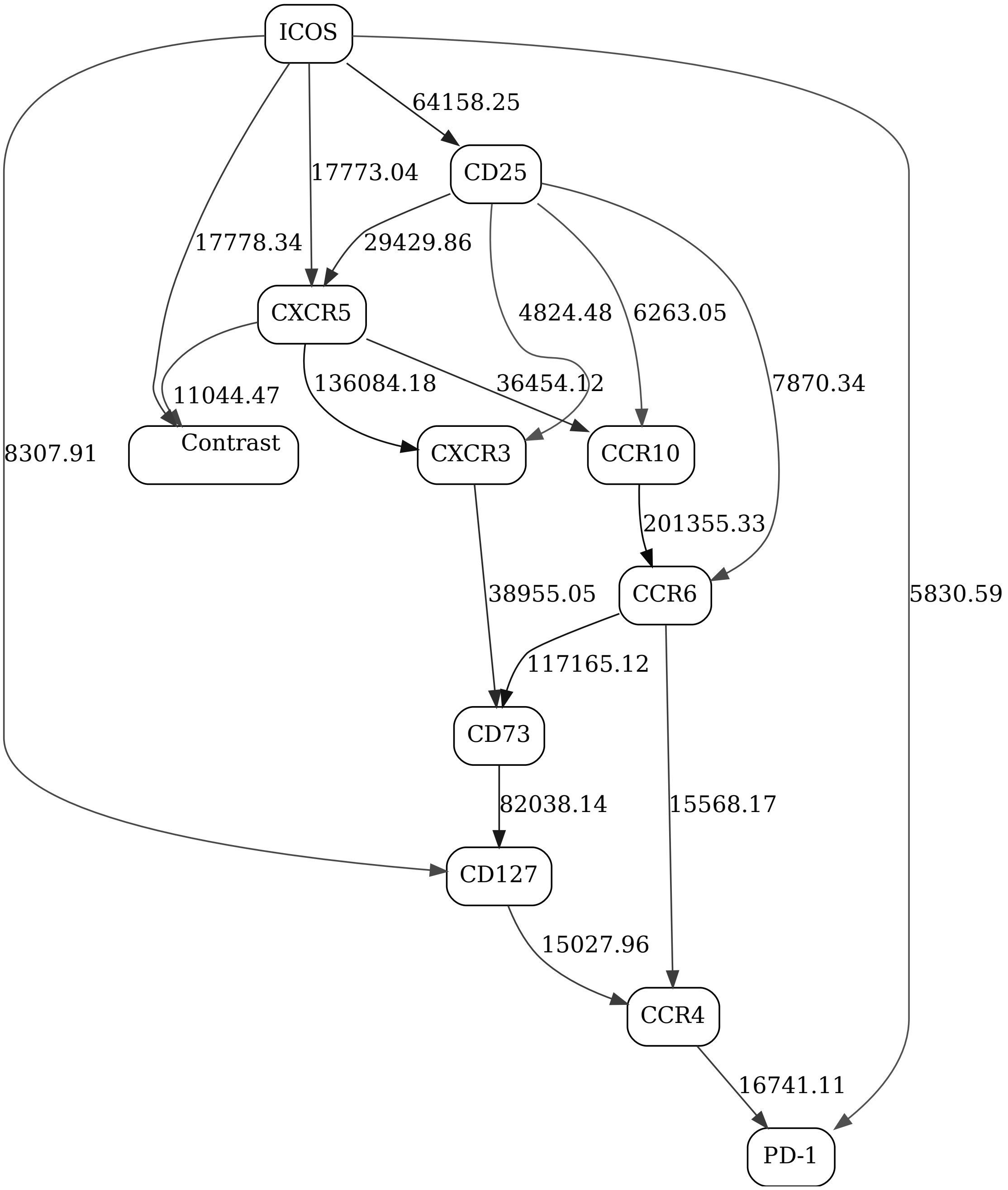


Figure 54: Adaptive immune signaling network panel, non-naive CD8

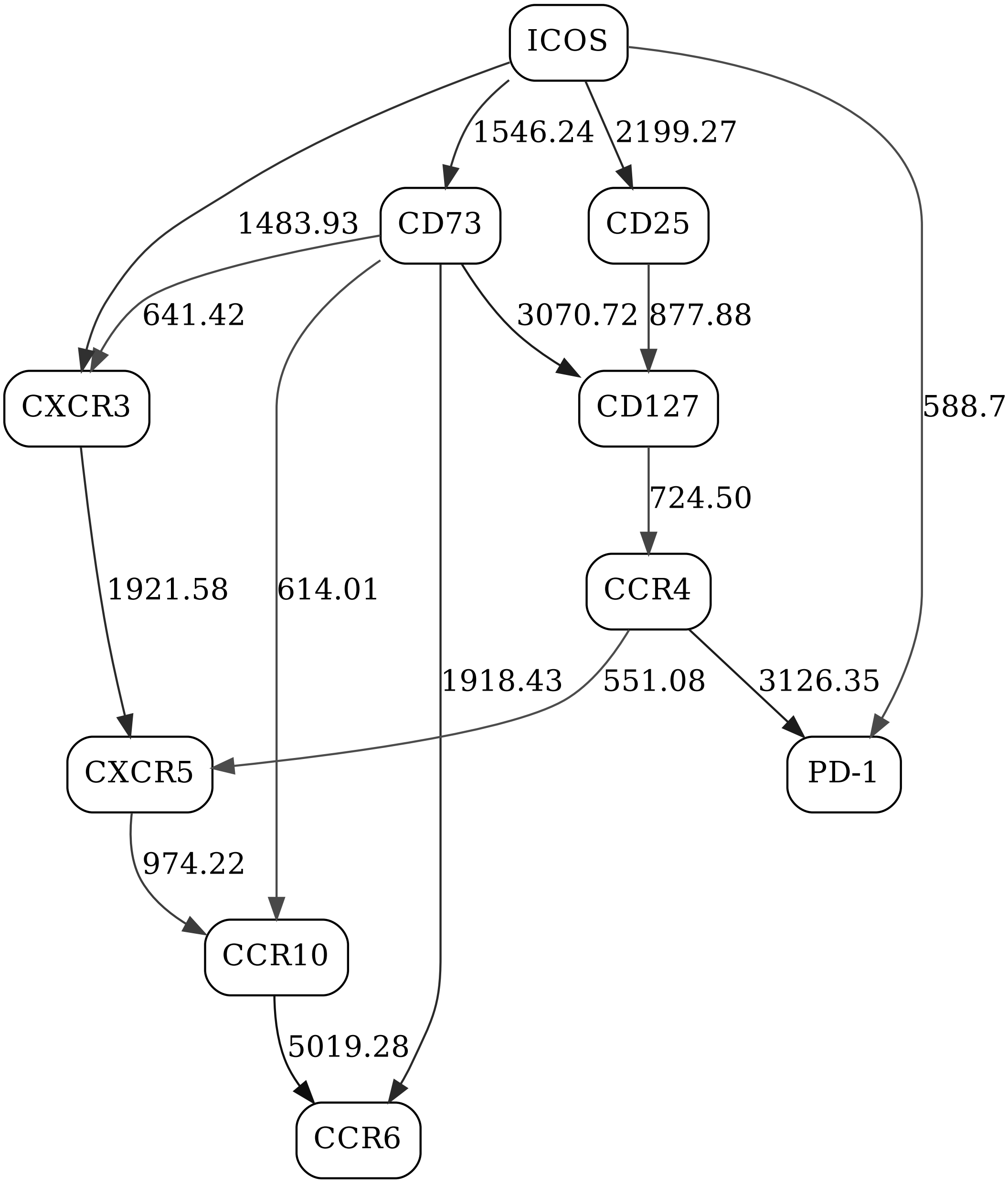


Figure 55: Adaptive immune signaling network panel, naive CD8, day 1, responders

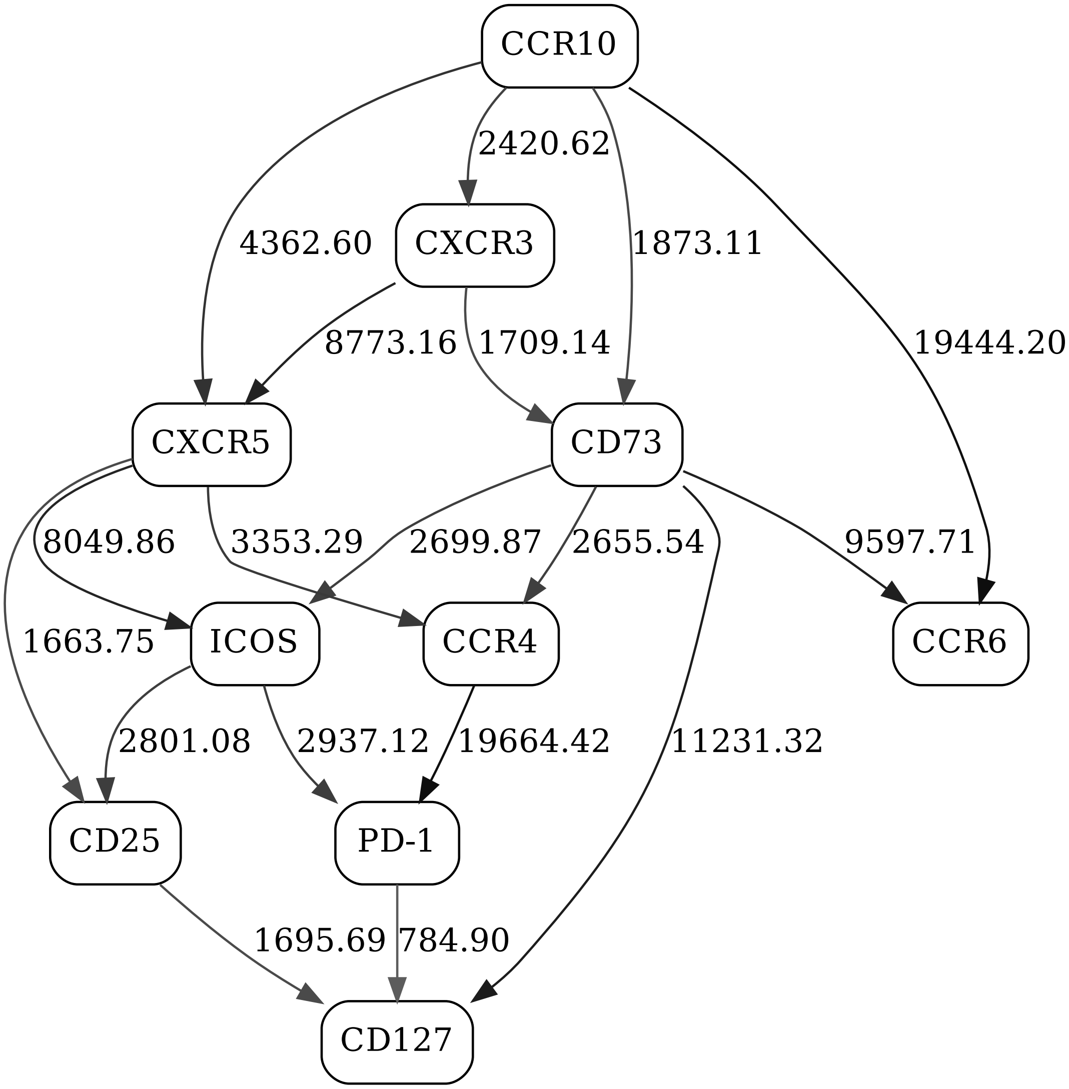


Figure 56: Adaptive immune signaling network panel, naive CD8, day 1, non-responders

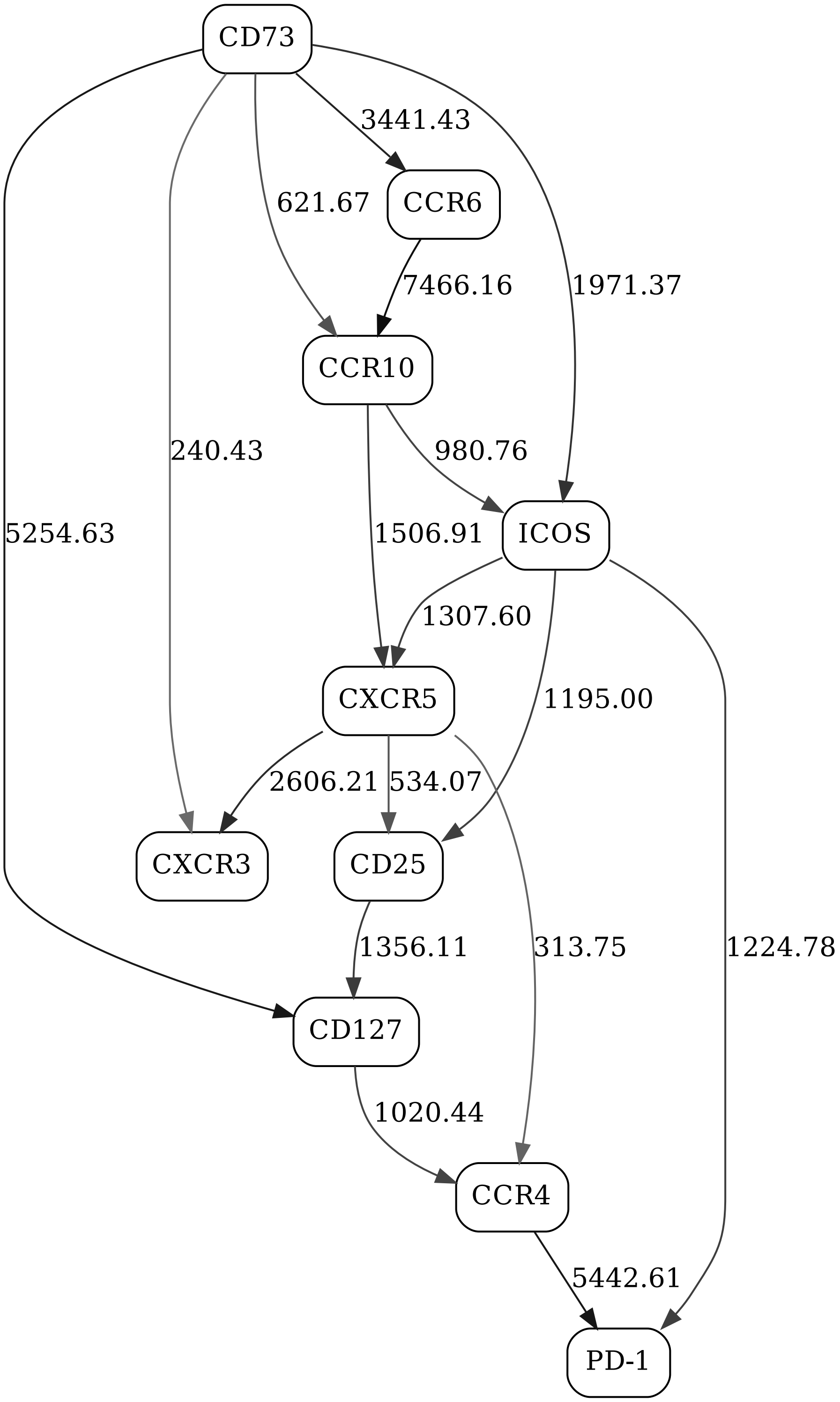


Figure 57: Adaptive immune signaling network panel, naive CD8, day 21, responders

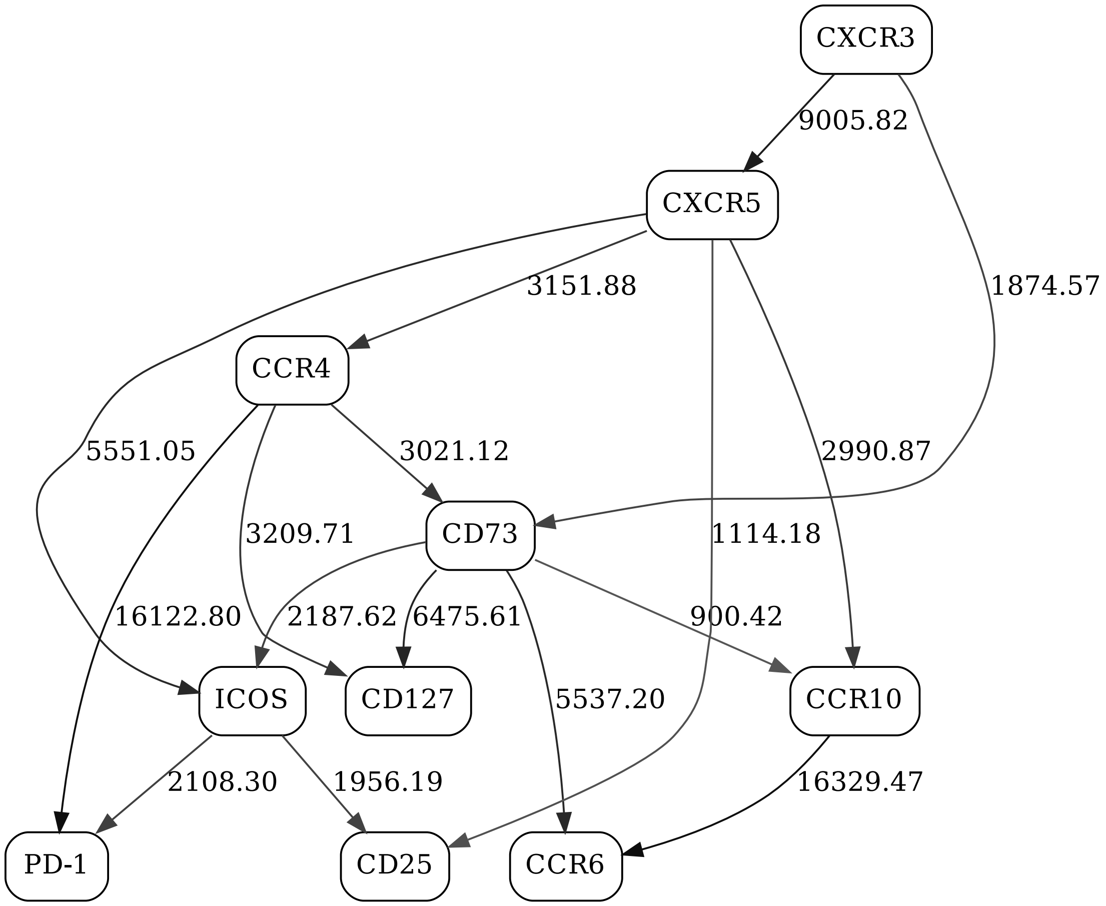


Figure 58: Adaptive immune signaling network panel, naive CD8, day 21, non-responders

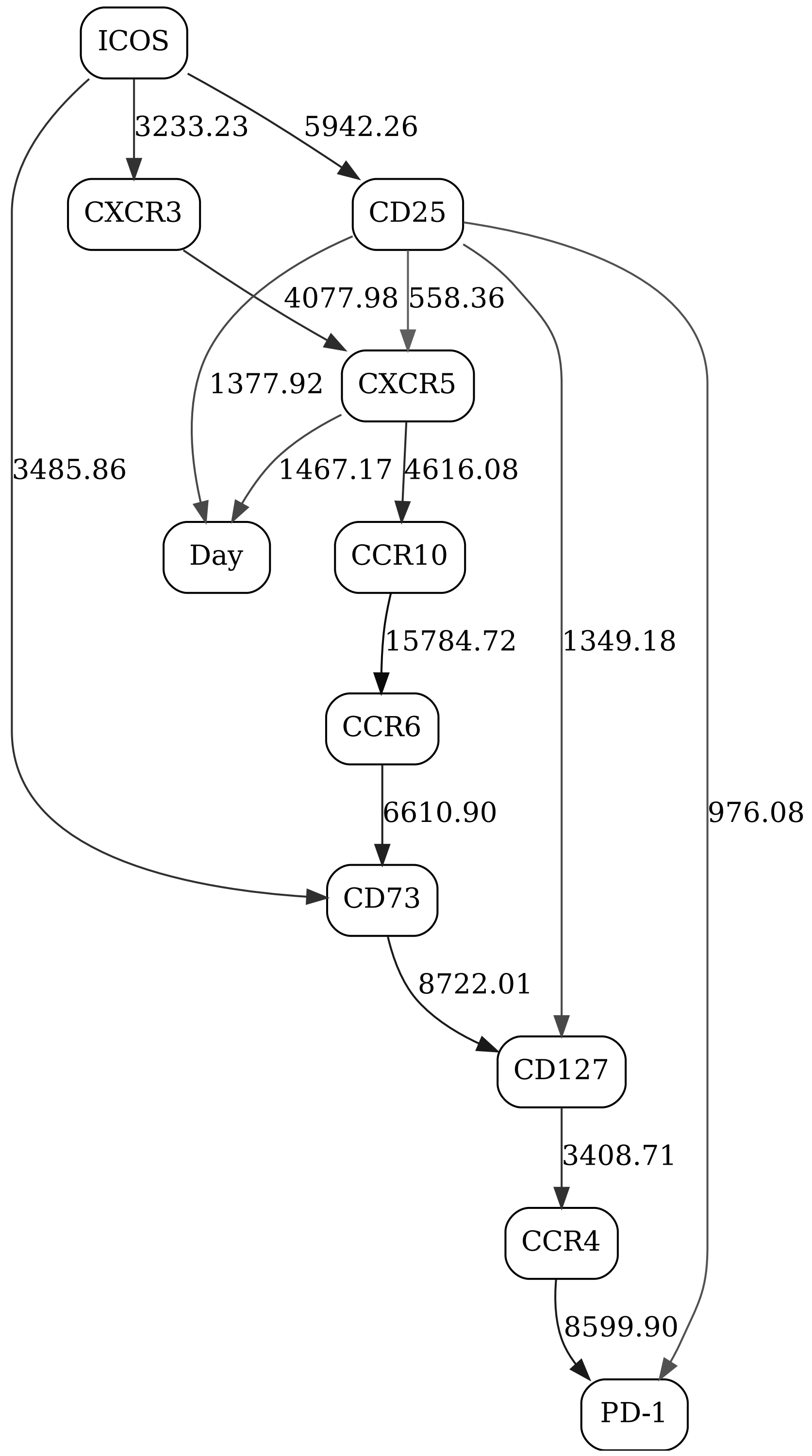


Figure 59: Adaptive immune signaling network panel, naive CD8, day contrast (2-state "Day" variable), responders

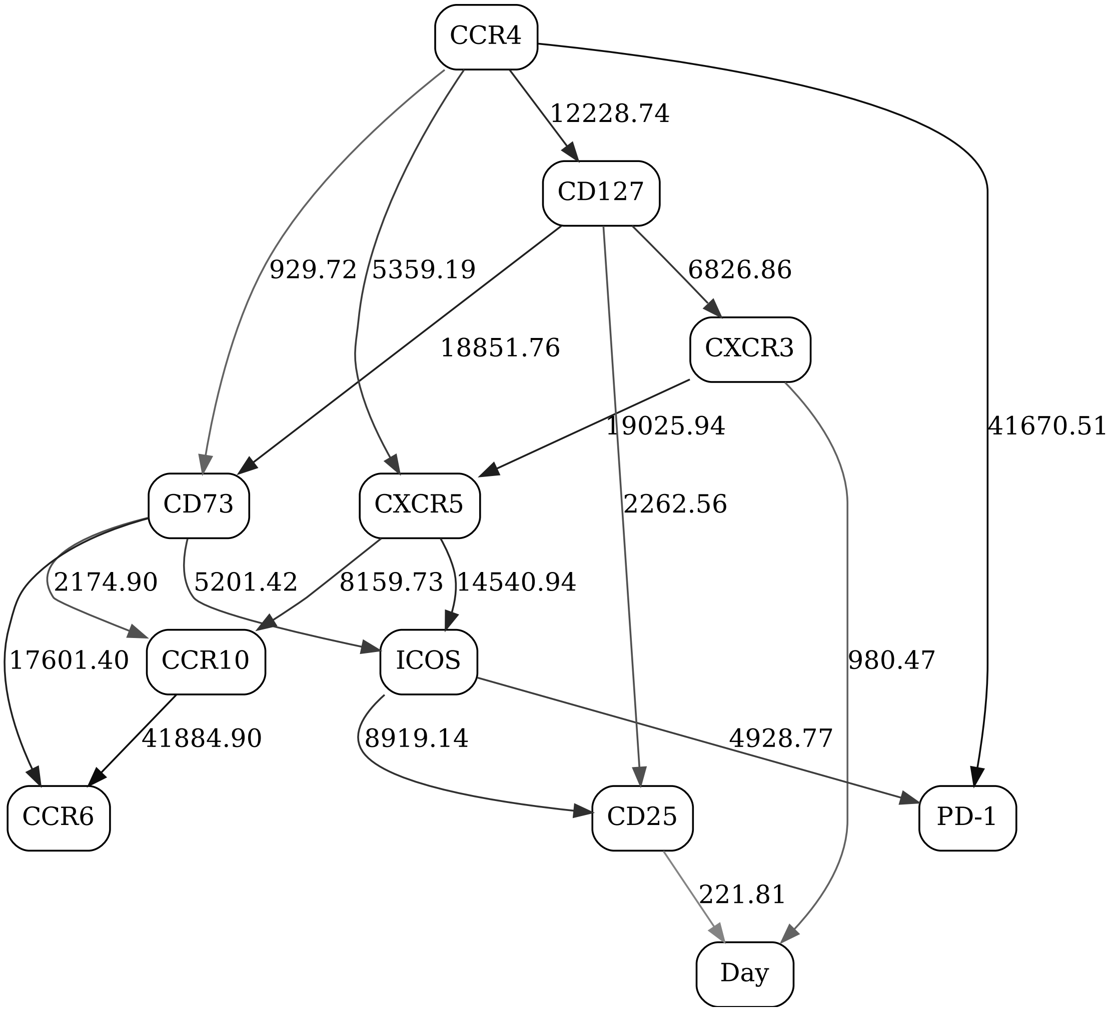


Figure 60: Adaptive immune signaling network panel, naive CD8, day contrast (2-state "Day" variable) non-responders

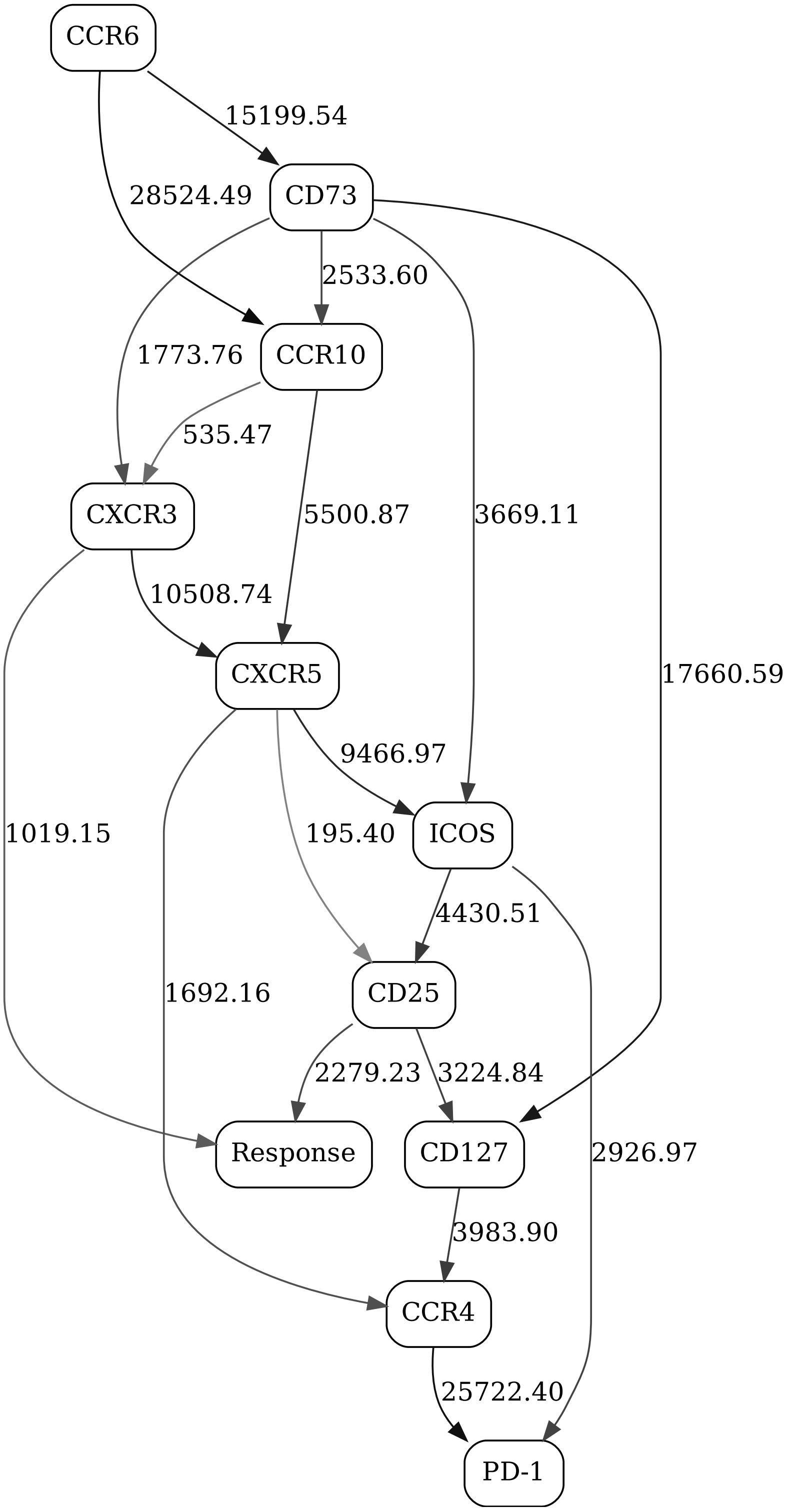


Figure 61: Adaptive immune signaling network panel, naive CD8, day 1, response contrast (2-state "Response" variable)

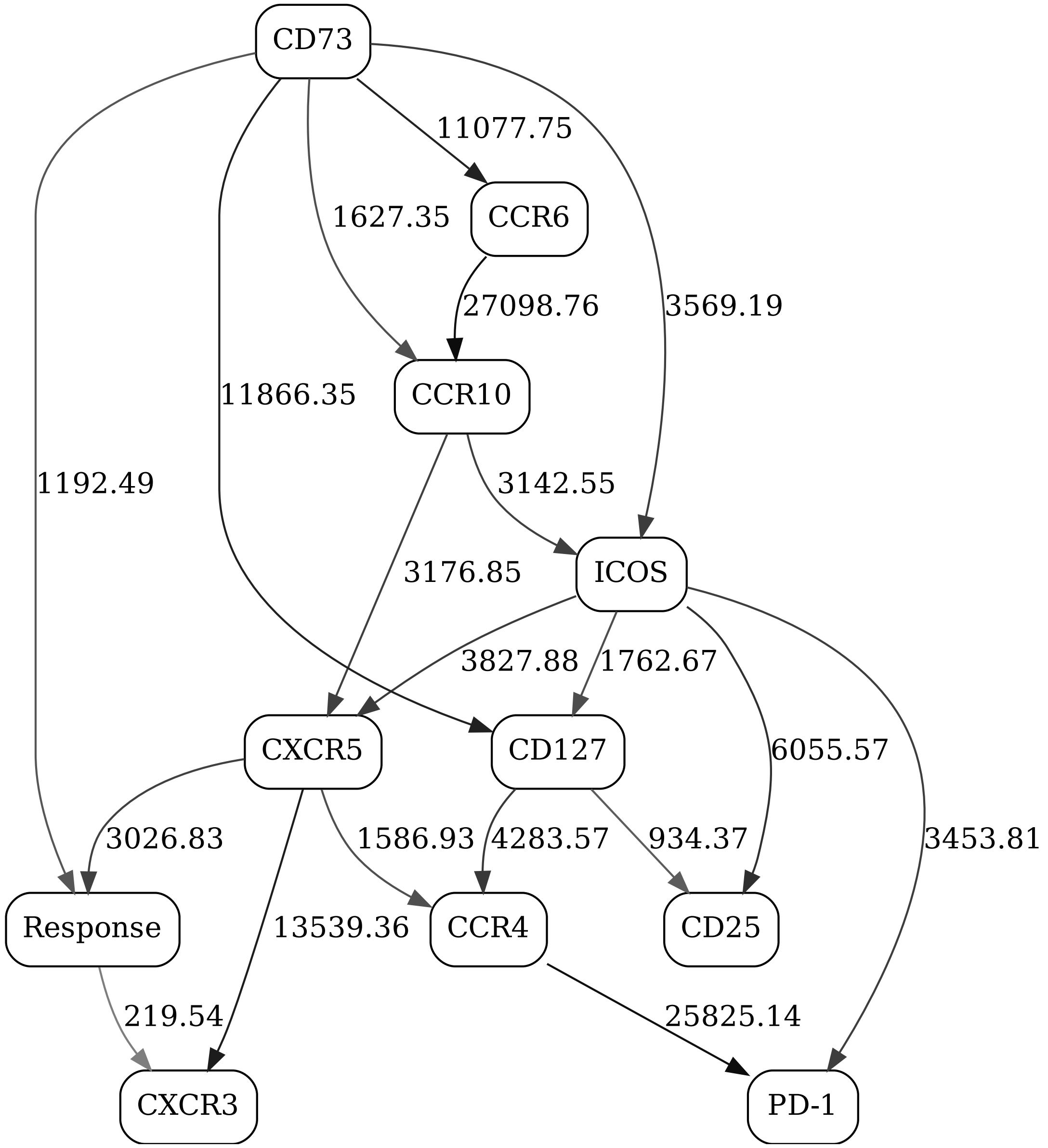


Figure 62: Adaptive immune signaling network panel, naive CD8, day 21, response contrast (2-state "Response" variable)

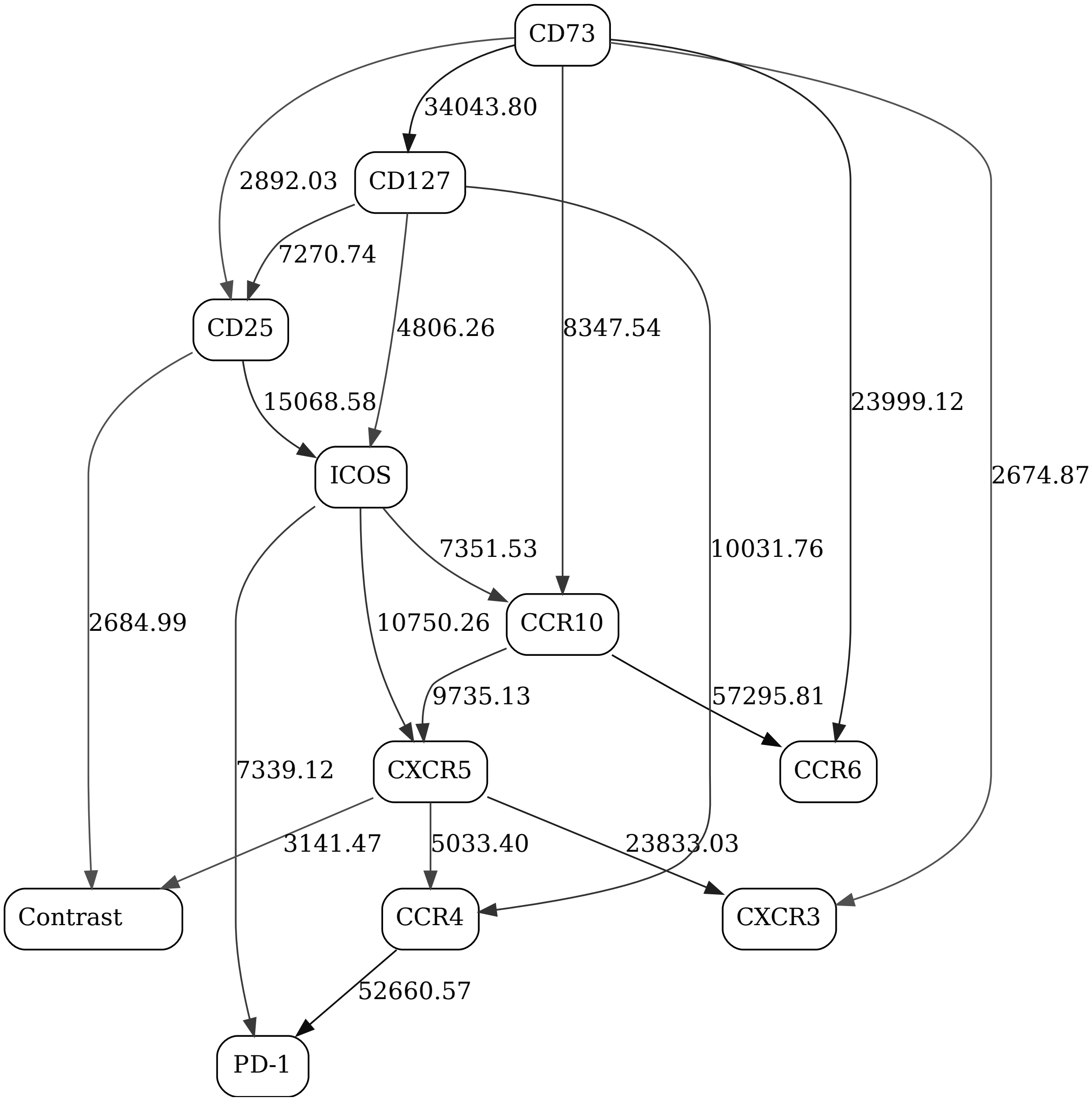


Figure 63: Adaptive immune signaling network panel, naive CD8
